# Effective Molecular Dynamics from Neural-Network Based Structure Prediction Models

**DOI:** 10.1101/2022.10.17.512476

**Authors:** Alexander Jussupow, Ville R. I. Kaila

## Abstract

Recent breakthroughs in neural-network based structure prediction methods, such as AlphaFold2 and RoseTTAFold, have dramatically improved the quality of computational protein structure prediction. These models also provide statistical confidence scores that can estimate uncertainties in the predicted structures, but it remains unclear to what extent these scores are related to the intrinsic conformational dynamics of proteins. Here we compare AlphaFold2 prediction scores with >60 μs of explicit molecular dynamics simulations of 28 one- and two-domain proteins with varying degree of flexibility. We demonstrate a strong correlation between the statistical prediction scores and the explicit motion derived from extensive atomistic molecular dynamics simulations, and further derive an elastic network model based on the statistical scores of AlphFold2 (AF-ENM), which we benchmark in combination with coarse-grained molecular dynamics simulations. We show that our AF-ENM method reproduces the global protein dynamics with improved accuracy, providing a powerful way to derive effective molecular dynamics using neural-network based structure prediction models.

## Introduction

AlphaFold2 ^1^ and RoseTTA fold ^2^ can infer protein structures based on sequence information with high accuracy. As a result, many projects have emerged that aim to improve the accessibility and usability of these structure prediction tools, most notably the AlphaFold Protein Structure Database ^3^, which currently provides predicted structures for over 200 million proteins ^4^, ColabFold ^5^, which combines fast homology search of MMseqs2 ^6^ with AlphaFold2 and RoseTTA fold, and derivations for to predict large protein complexes ^7, 8^.

AlphaFold uses the predicted Local Distance Difference Test (pLDDT) to estimate the accuracy of the predicted Cα positions (on a relative scale of 0-100) with experimental structures ^1, 9, 10^, but it remains unclear to what extent these measures can be linked to physical properties, such as protein dynamics. While low pLDDT scores have been associated with intrinsically disordered regions (IDRs) ^3, 10–13^, conditionally-folded IDRs, i.e., protein regions that fold in the presence of a binding partner or post-translational modifications, can show high (>70 - 90) pLDDT scores. In this regard, AlphaFold2 often appears to favor the folded state ^14^, possibly as these are structurally over-represented in the protein database. Highly dynamic protein conformations can also be identified by analyzing the combined pLDDT score, their solvent accessibility ^15^, and amino acid sequences that usually comprise a high proportion of charged and hydrophobic residues ^14^.

AlphaFold2 also uses the predicted aligned error (PAE) as another metric to estimate positional errors for a given residue, *i*, between the predicted and experimental structures that are aligned relative to residue *j* (capped at 31.75 Å) ^1, 3^, an approach adapted from superposition-based metrics ^16, 17^. PAE scores can be represented as a non-symmetric matrix related to the model confidence for the relative position and orientation of different protein segments. It is thus possible that the PAE scores could be used to determine rigid cores and protein domains ^11, 18^. Qualitative comparisons with atomistic molecular dynamics (aMD) simulations seem to confirm these observations ^19^. Recent studies have also explored other approaches to gain information about protein dynamics by using the variation in the predicted structures with AlphaFold2 ^20^, but a systematic strategy to infer protein dynamics from AlphaFold prediction scores is still lacking.

Here we compare AlphaFold prediction scores for a set of 28 different single and two-domain proteins against aMD simulations, with the systems chosen to capture proteins with varying degree of flexibility. Although aMD simulations provide an accurate basis for the structural characterization of the conformational space, highly dynamic proteins, e.g., with intrinsically disordered regions (IDRs) or flexible domains, remain challenging and limited in system size and the timescales needed to capture the major conformational changes ^21, 22^. To address this challenge, we propose a framework to convert the AlphaFold scores into elastic network models (ENMs) ^23–25^. The core assumption behind an ENM is that the protein dynamics can be adequately described by fluctuation around an initial, native structure and can be used to propagate effective molecular dynamics of the system at a low computational cost.

ENMs are often used in combination with coarse-grained MD (cgMD) simulations. To sample the large-scale conformational dynamics of biomolecules with cgMD models, multiple heavy atoms are grouped into individual interaction centers or beads, reducing the system’s degrees of freedom and allowing for larger computational time- and size-scales. However, these approximations also lead to a reduction in accuracy, particularly for proteins, which is partially accounted for by the restraints from the ENMs ^26^. Among the various coarse-grained approaches, the MARTINI force field ^24, 27–29^ has proven to be highly versatile due to its balance between chemical specificity and performance ^26, 30^. Initially developed for lipids ^27^, the MARTINI model was later extended to include parameters for proteins ^24^, DNA ^31^, RNA ^32^, carbohydrates ^33^, as well as many small molecules ^34^. The latest MARTINI3 version ^29^ has expanded available bead types, resulting in more accurate and detailed representations of functional groups, although both former and current models still struggle to properly balance the challenging protein-protein and protein-solvent interactions. In this regard, increased protein-water (P-W) interactions can improve the agreement with experimental data ^35–39^. Due to the introduced approximations, the MARTINI model also encounters difficulties in preserving secondary and tertiary structure elements of proteins, which is why it is often combined with ENMs ^23–25^ or Gō-type models ^40, 41^ in which the harmonic bonds are substituted by Lennard–Jones interactions.

After establishing a link between protein dynamics by benchmarking data from around 62 μs of aMD simulations and the statistical AlphaFold scores (pLDDT, PAE), we develop here an AlphaFold-optimized ENM (AF-ENM) and show that it can be used in combination with MARTINI cgMD simulations to increasing their usability for dynamical proteins.

## Results

### The correlation between AlphaFold scores and protein dynamics

To probe how the pLDDT and PAE scores correlate with the protein conformational dynamics, we performed aMD simulations of 28 different proteins (Fig. 1, Table S1). Each protein was simulated for 2 μs, except the single domain proteins (protein 24-26), which were simulated for 1 μs. Moreover, we used a 10.5 μs SAXS-reweighted ensemble for the two-domain model of the highly flexible heat shock protein 90 (NTD-MD construct of Hsp90, protein 28) from a previous study ^42^, leading to benchmarking data set of 61.5 μs aMD simulation. The aMD simulations were performed using the a99SB-disp force field, as it accurately captures both folded and disordered protein states ^43^. To assess the degree of flexibility of individual residues, we used the average standard deviation of the C_α_-distances to the 20 nearest amino acids (Øσ_d,20_). Unlike the widely used *root-mean-square-fluctuation* (RMSF) metric, the Øσ_d,20_ does not depend on structural alignment, and is therefore well suited for (highly flexible) multidomain proteins. To evaluate the global protein dynamics, we also used the standard deviation of all C_α_ distances (σ_d_), which can be represented as a matrix similar to the PAE scores. Overall, groups of residues with higher Øσ_d,20_, and σ_d_ values can be interpreted as regions with high conformational flexibility.

**Figure 1:**
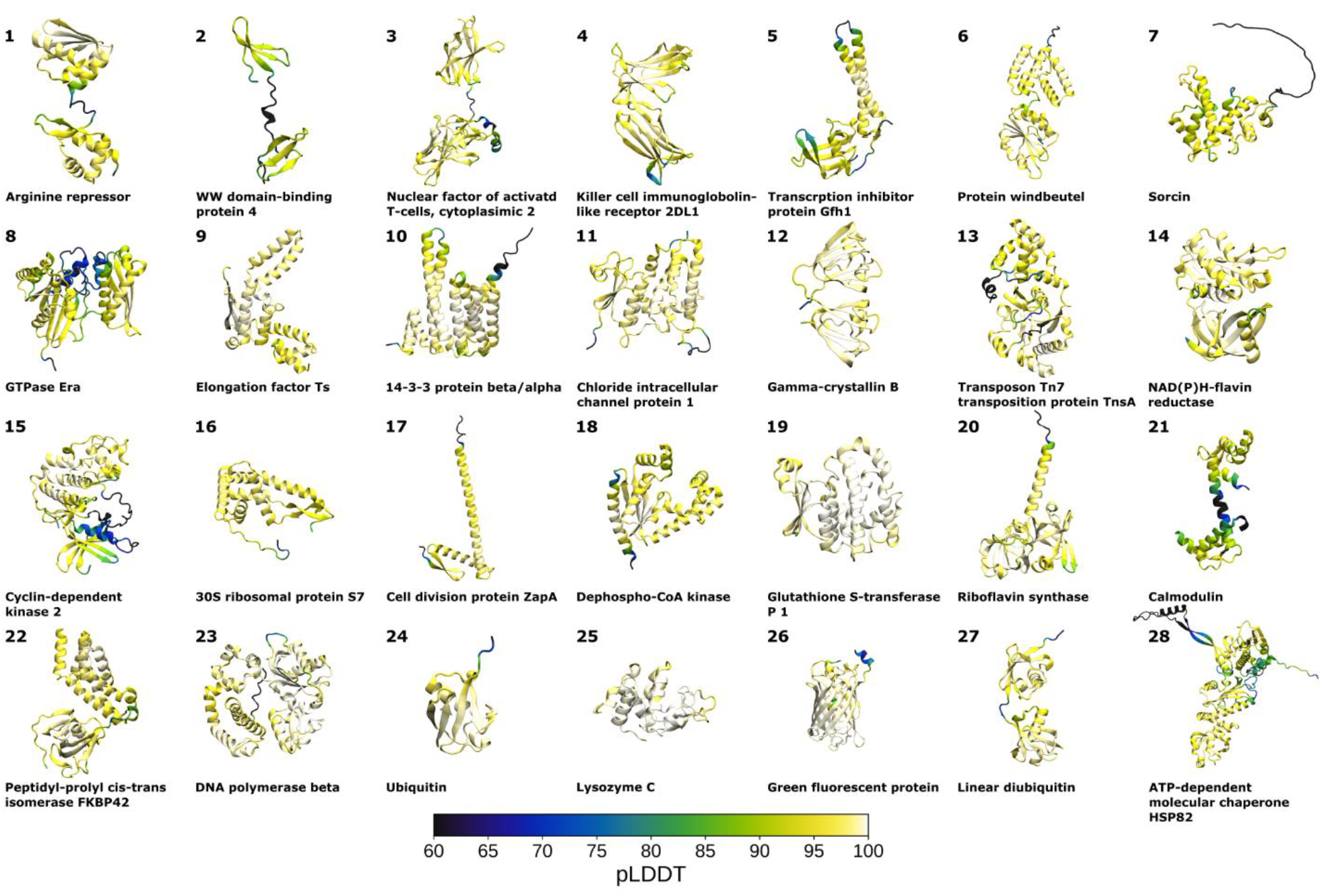
The structure of 28 proteins used to compare statistical scores of AlphaFold with dynamics derived from atomistic MD simulations. The proteins are colored based on their pLDDT scores, with pLDDT scores <= 60 (dark blue), and residues with a pLDDT ~100 (white).

A comparison between the PAE (in blue) and σ_d_ (in red) matrices, as well as the pLDDT and Øσ_d,20_ scores, for the 28 systems are shown in Fig. 2, with a more detailed analysis of the individual proteins in Fig. S1-S28. We note that high pLDDT scores correspond to low Øσ_d,20_ values, with an overall correlation coefficient *R* of 0.65, with a 1 Å increase in the Øσ_d,20_ corresponding to a 9 unit decrease of the pLDDT score. The overall linear regression across all tested proteins is:

**Figure 2.**
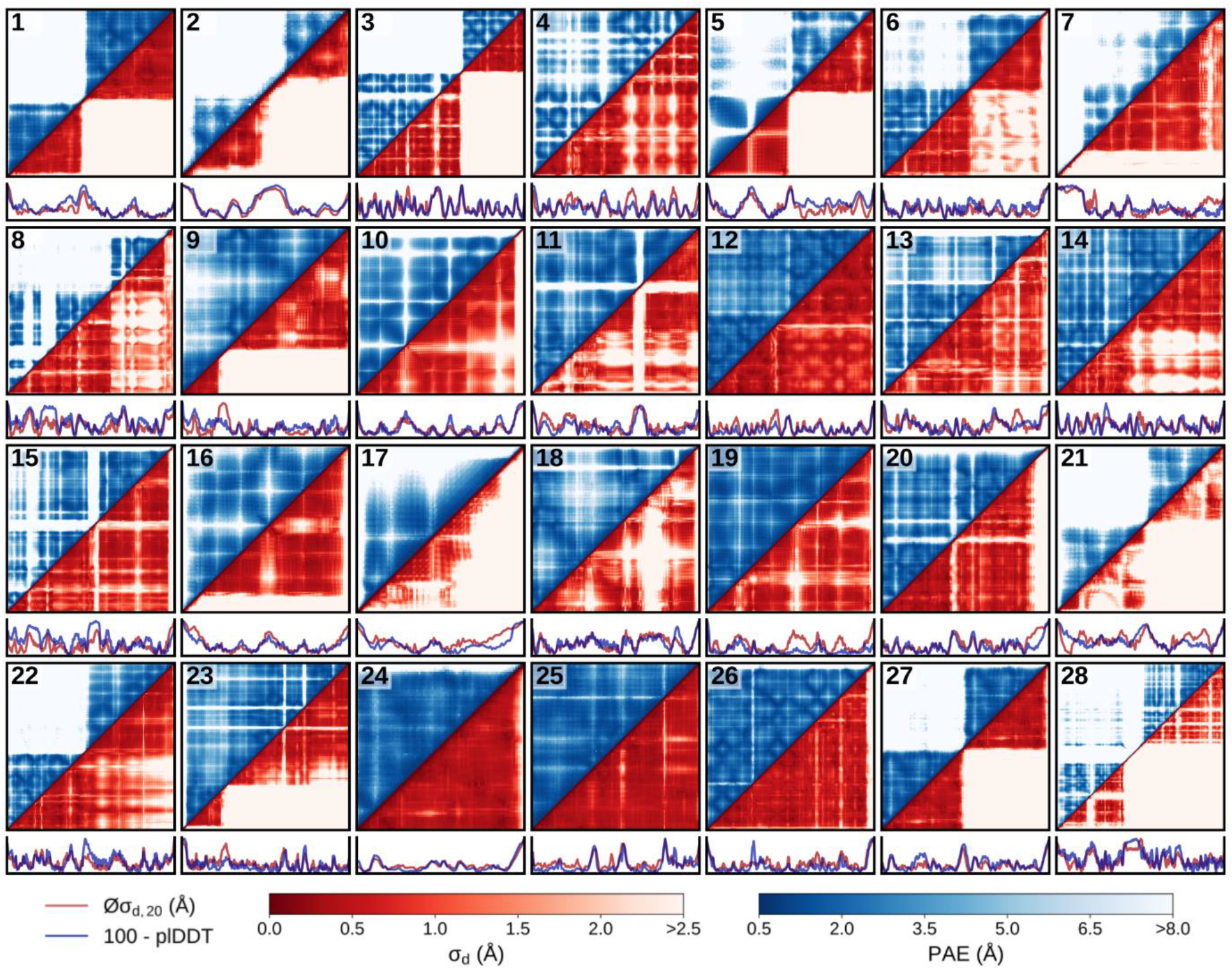
Overview of AlphaFold scores and protein dynamics metrics for all studied systems. A fully symmetric matrix indicates perfect correlation between the AlphaFold and MD data. The figure is divided in 28 subfigures for each of the proteins shown in Fig. 1. The top matrices show a comparison between (symmetrized) PAE matrices (blue) against the standard deviation of all C_α_ distances σ_d_ (red). The PAE scores range between 0.5 to 8.0, while the threshold for σ_d_ is set to <2.5 Å to enable better comparisons between the different systems. The bottom graphs compare the pLDDT scores (blue) and average standard deviation of the Cα-distances to the 20 closest amino acids Øσ_d,20_ (red). See Fig. S1-S28 for a detailed analysis of the individual systems.

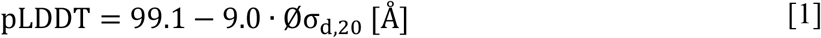

We observe that the R values for the regressions of individual systems vary between 0.24 to 0.99, with a median value of 0.74 (Table S2). While the slopes vary from −1.3 to −21.7 (median of −10.3), the intercept range is between 94.1-106.3 (median of 101.3). Regions with low pLDDT scores generally correspond to high Øσ_d,20_ values, particularly for disordered proteins regions, flexible loops, and linker regions. However, the relative heights of the peaks do not always align (e.g., Fig. S4a). Our analysis thus suggests that the pLDDT scores can be used to identify more flexible regions within a given protein rather than comparing the absolute dynamics between different systems. Based on the regression between the PAE scores and σ_d_ metric across **all** systems, we find an *R* of 0.53, which is lower than for pLDDT - Øσ_d,20_. Overall, a σ_d_ increase of 1 Å corresponds to a rise in PAE of 0.7 Å:

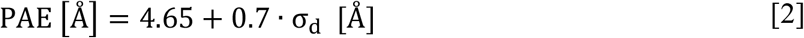

The *R* values for individual systems range from 0.25 to 0.92 (median 0.65), with intercepts within the 0.10 to 6.27 range (median 2.4), and the slopes between 0.38 and 4.83 (median 2.1), suggesting that the comparison of PAE values between the different systems have considerable uncertainties, but rather good correlation to the explicit dynamics within the same system (Table S3). While the overall correlations are not necessarily high enough for a quantitative assessment of protein dynamics, we later show that a threshold-based approach can reliably differentiate between rigid and flexible regions and be used for constructing ENMs (see below).

Overall, the qualitative consistency of the pLDDT and PAE scores across all studied protein systems is striking. We notice that a PAE score of ~8 Å appears to set a good threshold to visibly identify highly flexible regions as well as dynamically separated domains, corresponding to a σ_d_ of around 2.5 Å. For example, for protein systems 1, 2, 3, 21, and 27 (arginine repressor, WW domain-binding protein 4, nuclear factor of activated T-cells cytoplasmic 2, calmodulin, and linear diubiquitin), the PAE threshold correctly predicts the high inter-domain flexibility. AlphaFold also appropriately assesses the increased flexibility of N-terminal (system 7 and 16, sorcin and 30S ribosomal) and C-terminal tails (e.g., system 17, 20, 24, and 26 (cell division protein ZapA, riboflavin synthase, ubiquitin, and green fluorescent protein)), as well as dynamic linker regions (*e*.*g*., systems 2, 11 and 28 (WW domain-binding protein 4, chloride intracellular channel protein 1, and NTD-MD construct of Hsp90)). Generally, residues with lower pLDDT scores show higher PAE scores for residue pairs, suggesting that the PAE matrix also encodes the local dynamics, *e*.*g*., of flexible loop regions. This close link between the scores might explain some of the observed discrepancies between PAE scores and the σ_d_ metric in systems 9, 22, and 23 (elongation factor Ts, peptidyl-prolyl cis-trans isomerase FKBP42, and DNA polymerase beta). The differences could be attributable to the over- or underestimation of the conformational dynamics in a small linker section between two domains, with errors that could propagate and lead to significant changes in the PAE scores.

Based on a closer look at our largest and most complicated system within the dataset, the NTD-MD construct of Hsp90 (protein 28, Fig. S28), we observe an interplay of the abovementioned effects. This system is composed of an N-terminal domain (NTD) and a middle domain (MD), which are connected by a highly charged disordered linker (CL) region ^44–48^. In a previous study, we showed that conformational changes between compact and extended states are enabled by the dissociation of the NTD from the MD ^42^. Moreover, additional conformational dynamics is enabled by the flexible lid region in the NTD, which has well-established open and closed states, and a potential unfolding of a β-sheet comprising β-strand 8 from the CL and β-strand 7 from the NTD ^44, 48^, with partial unfolding leading to highly extended conformations ^42^. Thus, the Hsp90 NTD-MD model comprises a mixture of rigid domains, IDRs, and conditionally folded states. We indeed observe that the pLDDT scores and the Øσ_d,20_ metrics for the Hsp90 construct within the individual regions are consistent with the flexibility derived from the aMD simulations. The exception is the β8-strand, where we notice a pLDDT score of >90 for one of the highly dynamic regions. This finding is consistent with the observation that AlphaFold generally predicts high pLDDT values for residues in conditionally folded areas ^14^. The PAE and σd scores between the NTD and MD domains are indeed significantly higher than those within each domain. For residues in the CL, both scores are also elevated, except for β-strand 8.

Thus, our analysis suggests that the pLDDT scores can be closely linked to local protein dynamics, whereas the PAE scores capture large-scale conformational flexibility, with both scores being closely related to each other. We also find that relatively small errors in the pLDDT scores easily propagate large mismatches between the PAE matrix and global protein dynamics.

### Using AlphaFold scores to construct elastic network models

To utilize the AlphaFold scores in explicit molecular dynamics simulations, we propose a framework to convert the statistical pLDDT and PAE scores into an elastic network model (ENM) ^23–25^. An ENM can be constructed by adding harmonic potentials between, *e*.*g*., C_α_ atoms of the protein backbone of residues *i* and *j* according to,

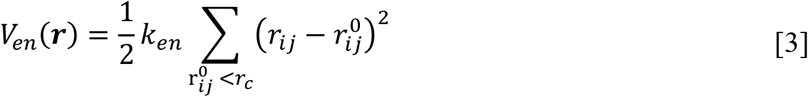

where *V*(*en*) is the total potential energy of the elastic network (*en*), *r*_C_ is the cutoff radius, and *k*_en_ is the force constant for the harmonic bond. The ENM additionally relies on reference distances 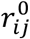 inferred from an initial reference structure. In the simplest formulation, a generic constant is used for all *k*_en_s, although this approach becomes inaccurate for proteins with multiple prominent conformational states. ENMs can also be generated based on short-timescale aMD simulations (in the order of tens of ns) in which both the number as well as the strength of bonds are optimized ^49–51^. However, many processes, such as the dissociation events and large-scale conformational changes, lie outside the short sampling timescales, making this approach less well suited for highly dynamical systems ^49^. Here we propose an alternative strategy to adapt bonds and force constant based on AlphaFold scores.

For our AlphaFold optimized elastic network model (AF-ENM), we suggest scaling the force constants based on the PAE scores. To this end, we assume that the distance distribution between the C_α_ atoms corresponds to a normal distribution, with the predicted aligned error as the standard deviation. The distance distribution P for a given distance between residues *i* and *j* can therefore be defined as,

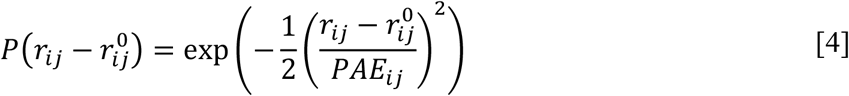

Since the PAE matrix is generally not symmetrical, we use the average value between residue PAEij and PAEji. The relative free energy ΔG(r) can be calculated from the distance distribution,

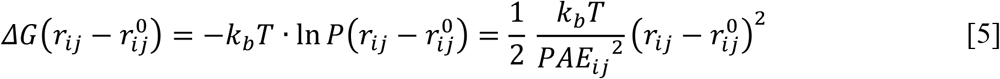

By combining equations [3] and [5], the force constant ken for the bond between residues *i* and *j* can be defined as,

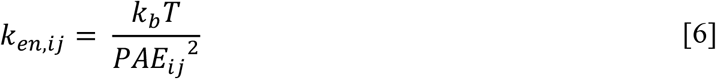

where *k*_*b*_ is the Boltzmann constant and *T* is the temperature.

NMs are typically derived based on a single structure, but this can result in a significant underestimation of the global conformational dynamics. If multiple structures are available, it often remains unclear which of them (if any) would be suited to construct the ENM. For example, linear diubiquitin (Ub2, Fig. S27) has different potential initial structures (Fig. 3a). Ub_2_ is a highly flexible two-domain protein ^38, 52^, in which the C-terminus of a proximal ubiquitin is linked to the N-terminus of a distal ubiquitin subunit. Different experimental structures have resolved Ub_2_ in both compact (PDB ID: 3AXC ^53^ and 4ZQS ^52^) and open (PDB ID: 2W9N ^54^) states, with Fig. 3a also showing Ub_2_ as predicted by AlphaFold2 ^1^. Depending on the initial conformations, the ENMs between the individual Ub-domains are vastly different, with the compact state (PDB ID: 4ZQS) having a high degree of bonds between the subunits, whereas the open model (PDB ID: 2W9N) shows no contacts.

**Figure 3.**
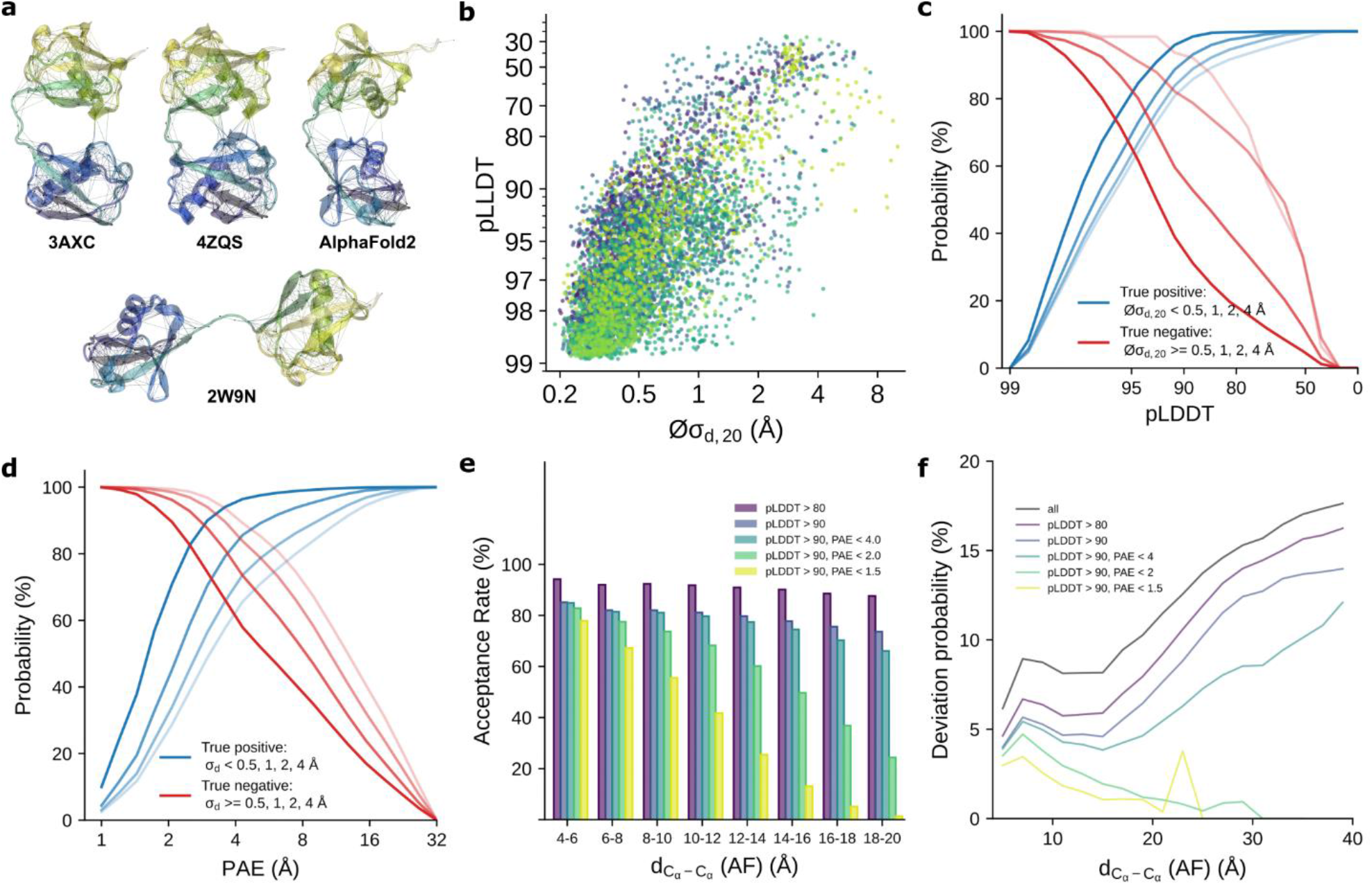
Construction of an elastic network model (ENM) based on AlphaFold for **a**) diubiquitin (Ub_2_) ENMs with different starting structures. The protein is colored according to the residue number (from blue to yellow). The bonds and ENM (cutoff distance of 0.9 nm) between all C_α_ atoms are shown. **b**) Øσ_d,20_ vs. pLDDT scatter plots combined across all systems shown in Fig. 1. **c**) Sensitivity and specificity for the pLDDT/ Øσ_d,20_ relations. The blue curves show the probability that a Øσ_d,20_ value is *below* a cutoff value (represented by different transparency levels) when the corresponding pLDDT score is above a threshold value (x-axis). The red curves show the probability that Øσ_d,20_ is *above* a cutoff value for residues with a pLDDT score *below* a threshold value. **d**) Sensitivity and specificity for the σ_d_/PAE relationship. The blue curves show the probability that a PAE score (x-axis) is *below* a threshold value and is associated with a σ_d_ *below* a threshold value (represented by different transparency levels). The red curves show the probability that a PAE and associated σ_d_ are *above* threshold values (true negative test). **e**) Probability that an AF predicted distance deviates by more than 20% from the average distance estimated by aMD under different selection criteria. **f**) Ratio of aMD-derived distances that differ by more than 20% from the AlphaFold predicted structure.

**Figure 4.**
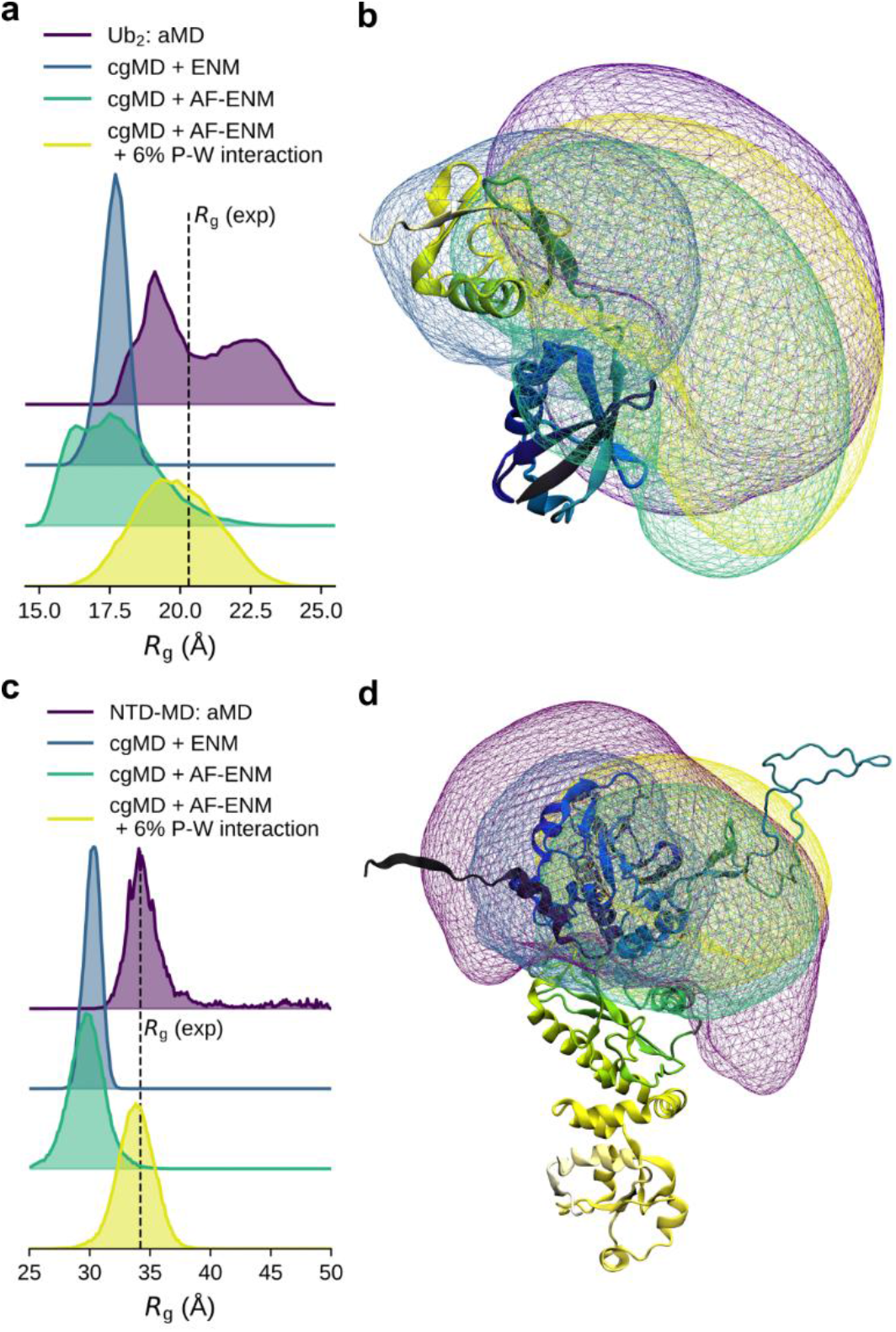
Comparison between aMD and cgMD simulations for linear Ub_2_ (protein 27) and the NTD-MD construct of Hsp90 (protein 28) a,c) Radius of gyration (*R*_*g*_) distributions obtained by aMD and cgMD simulations for (a) linear Ub_2_ and (c) the Hsp90 construct. b,d) Comparison of the conformational space occupied by (b) linear Ub_2_ and (d) the Hsp90 construct. The purple areas show the occupied space in the aMD simulations by the proximal ubiquitin relative to (b) the distal one and (d) the N-domain relative to the M domain of Hsp90. The blue areas show the occupied space in the cgMD simulations with ENF, the green areas correspond to the cgMD simulations with AF-ENM, while the yellow areas relate to the cgMD simulations with increased protein-water interactions.

To account for this bias, we propose to use the AlphaFold scores to remove network bonds with high expected fluctuation (Fig. 3). To this end, we used the pLDDT scores to identify residues of the rigid protein cores (low Øσ_d,20_) and flexible/disordered regions (high Øσ_d,20_) that we further used to construct an optimized AF-ENM. We identify a threshold that balances the probability of finding a pLDDT score above a specific value, corresponding to a low Øσ_d,20_ (sensitivity or true positive rate) with the likelihood that the pLDDT score is below the threshold associated with a high Øσ_d,20_ value (specificity or true negative rate). The specificity and sensitivity show an inverse relationship (Fig. 3c). We observe an excellent balance between them for a pLDDT threshold of 90, demonstrating a true positive rate of 86.5% and a true negative rate of 80.1% at a Øσd,20 threshold of 2 Å, as compared to 63.7% and 92.2% for a pLDDT threshold of 95, and 95.2% and 66.6% for a pLDDT threshold of 80.

While the pLDDT scores correlate with the local dynamics, the PAE scores are also good indicators of the global dynamics. We therefore use the PAEs to avoid over-constraining the conformational dynamics between two rigid regions in our AF-ENM. Similar to the dependency between pLDDT scores and Øσ_d,20_, we obtain a good balance between the sensitivity and specificity of the PAE/σ_d_ relationship (Fig. 3d), as indicated by the likelihood that a low PAE score corresponds to a low σ_d_ (sensitivity or true positive test) relative to the probability that a high PAE value corresponds to a high σ_d_ (specificity or true negative test). As for the pLDDT Øσ_d,20_ relationship, we observe that the true positive and negative rates have an inverse relationship. Since the addition of a bond that spuriously restricts the dynamics is far more problematic than the absence of bonds in a rigid structure, we suggest that high sensitivity is more important than high specificity within the AF-ENM model. In this regard, we find that a PAE threshold of 2 Å provides a specificity of 98.8% and a sensitivity of 30.8%, compared to 99.8% and 14.0% for a PAE threshold of 1.5 Å, and 86.6% and 70.1% for a PAE threshold of 4 Å.

Finally, we also probed the effect of different *r*_*c*_ on our AF-ENMs. We find that increasing the distance cutoffs correlate with increasing PAE values (Fig. S29), with the likelihood of including a given bond in the AF-ENM strongly depending on the C_α_-C_α_ distances (Fig. 3e). By only using a pLDDT threshold of 90, the acceptance rate is around 82% for C_α_-C_α_ distances between 8-10 Å, and 73% for C_α_-C_α_ distances between 18-20 Å (Fig. 3e). In contrast, when a PAE threshold of 2.0 Å is added, these values decrease to 74% and 24%, while the PAE threshold of 1.5 Å leads to 56% and 1.4%. To assess the extent to which the accepted bonds are representative of the structural ensemble, we examined the probability of the discrepancy of C_α_-C_α_ distances from AlphaFold and aMD simulations by more than 20% (Fig. 3e). We observe that the deviation probability for a cutoff radius of 10 Å changes from 8.1% to 3.2% by including only distances with a pLDDT > 90 and PAE < 2 Å.

Based on these results, we used the following framework to construct AF-ENMs for MARTINI3 cgMD simulations: starting from an initial structure, we only considered residues with pLDDT scores > 90 that are separated by at least two residues since the angle and bonded interactions account for their dynamics. Moreover, to balance sensitivity and selectivity, we used a cutoff radius of 9 Å, with force constants determined by the PAE scores based on equation [6]. For the AF-ENM cgMD simulations, we only consider bonds with *k* > 0.75 kJ/(mol · Å^2^), corresponding to a PAE threshold of 1.85 Å.

### Testing the AF-ENM together with Martini3

To test the AF-ENM approach, we compared MARTINI3-cgMD simulations with a standard ENM using a generic *k*_*en*_ (5 kJ/(mol · Å^2^) and the AF-ENM for three single domain (protein 24-26, ubiquitin, lysozyme C, green fluorescent protein) and two proteins with two domains (protein 27 and 28, linear diubiquitin, NTD-MD construct of Hsp90) against aMD simulations. To estimate the difference in global dynamics, we performed principal component analysis (PCA) on the aMD trajectories (C_α_ only) to extract the two components with the highest variance ^55, 56^. These components represent the proteins’ modes of motion with the most extensive conformational changes. We then project the backbone beads of the cgMD trajectories onto these modes and compare the resulting distributions with the multivariate Kullback-Leibler (KL) divergence ^57, 58^ (Fig. S30-S34, Table S4). A lower KL divergence corresponds to a higher similarity of two distributions and, therefore, a higher agreement between the global dynamics described by aMD and cgMD simulations.

Overall, we find that using AF-ENM instead of standard ENM leads to lower KL divergence (Table S4). The difference is particularly pronounced for systems with higher conformational dynamics. For small rigid ubiquitin (Fig. S30), we observe a KL divergence of 1.1 for the cgMD simulation with ENM compared to 1.8 with AF-ENM. In the case of lysozyme C (Fig. S31), which has more flexible loop regions, we observe a reduction in KL divergence from 4.5 to 1.4. GFP with the flexible N-terminal helix (Fig. S32) shows a KL divergence of 11.0 and 4.8 for ENM and AF-ENM, a similar reduction as for the larger and flexible NTD and MD domains (Fig. S34), where we notice a decrease from 11.9 to 1.2 and from 9.0 to 4.8. This is also consistent if comparing two domains, where we observe a KL divergence of 9.4 and 5.9 for linear diubiquitin (Fig. S33) and 10.5 and 4.4 for the combined NTD-MD construct (Fig. S34). This trend appears to be consistent for variations in the elastic network model, as we observe similar KL divergences if we double the force constants for our AF-ENM simulations (Table S4).

The advantage of using the AF-ENM with MARTINI3 cgMD simulations is particularly evident when combined with increased protein-water (P-W) interactions 35-39. In Figure 5, we compare the radius of gyration (*R*_*g*_) distributions of our aMD ensembles with the cgMD simulations for linear Ub2 and the Hsp90 NTD-MD construct. Using a standard elastic network leads to an overly constrained and compact conformation, which fails to reproduce the experimental *R*_*g*_ value obtained from small-angle X-ray scattering (SAXS) experiments or the *R*_*g*_ distribution from aMD simulations. For Ub2, we observe an average *R*_*g*_ of 17.6 Å in our ENM simulation compared to 20.3 Å experimentally ^38^. At the same time, for the Hsp90 construct, the corresponding values are 30.2 Å and 34.5 Å ^42^. Using our AF-ENM increases conformational freedom and leads to broader *R*_*g*_ distributions centered around similar values compared to the ENM simulations (17.7 Å for Ub_2_ and 29.7 Å for NTD-MD construct). However, the increase in P-W interaction in combination with the AF-ENM simulations resulted in higher ratios of open conformations, which are in better agreement with aMD ensembles and experimental data, with average *R*_*g*_ values of 19.9 Å and 33.8 Å for Ub_2_ and the Hsp90 construct, respectively. Overall, combining the AF-ENM framework can improve the quality of the protein discerption in cgMD simulations relative to standard ENM.

**Figure 5.**
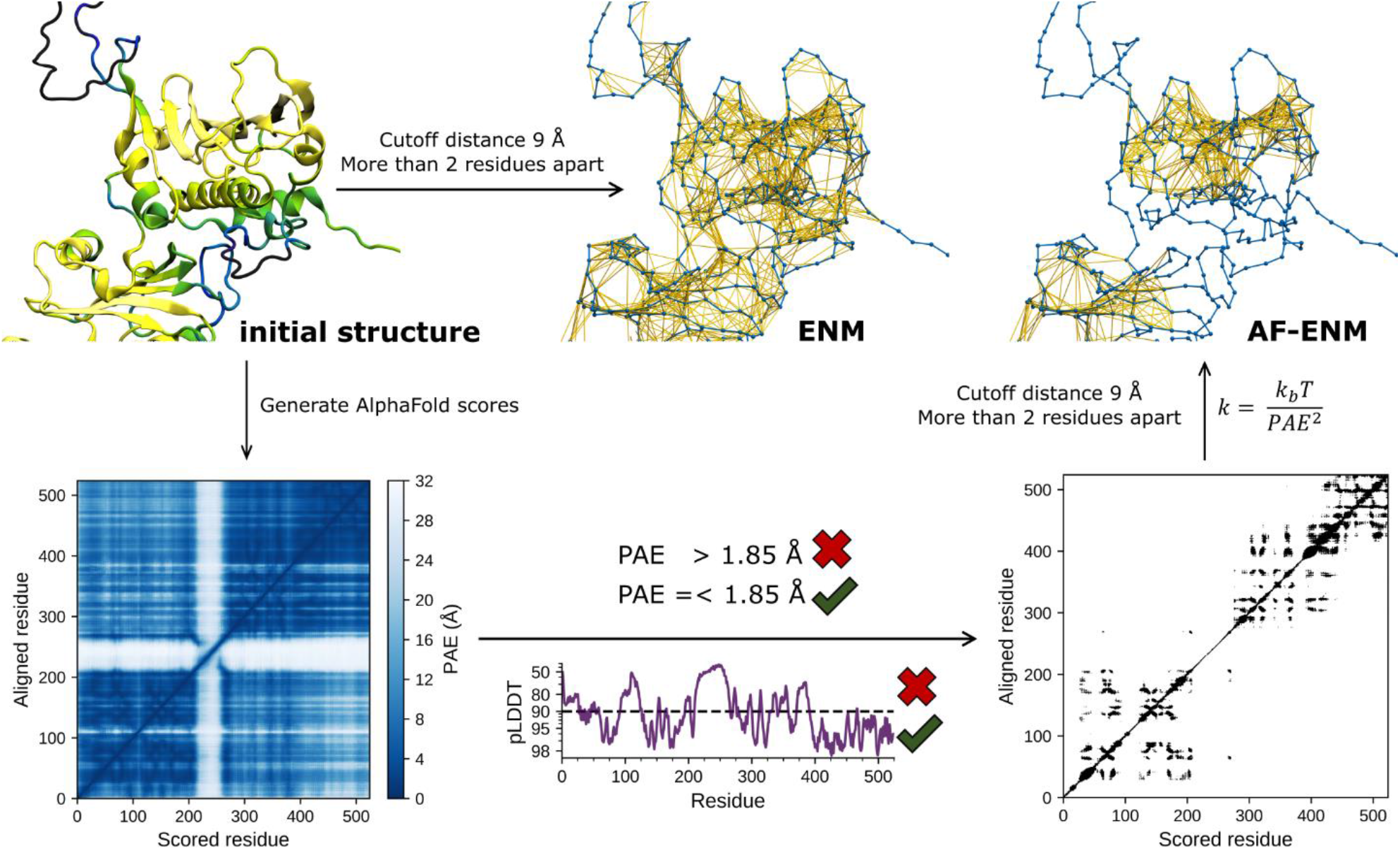
Schematic summary of the AF-ENM generation.

## Discussion

While AlphaFold is primarily used to predict protein structures, it is becoming increasingly clear that it can also be a valuable tool for inferring protein dynamics as shown here (*cf*. also Refs. ^11, 14, 15, 18, 19^). Here we performed a comprehensive comparison of 28 different systems to link AlphaFold confidence scores to metrics obtained by atomistic molecular dynamics simulations. We show that the pLDDT scores correlate with the local residual dynamics, while the PAE scores also provide insight into large-scale conformational dynamics. Although these scores are suited to unambiguously identify highly flexible and rigid protein regions and inter-domain interactions, accurately describing conditionally folded states remains challenging.

Our quantitative analysis allowed us to construct elastic networks based on these statistical scores (Fig. 6), that can be effectively combined with cgMD simulations. For this, we take advantage of AlphaFold’s ability to identify flexible regions between rigid domains. We find that a pLDDT threshold of 90 is sufficient to reliably specify well-defined structural elements, which should be conserved by the network. A PAE threshold between 1.5 to 2.0 Å appears to be well suited to avoid overly restrictive bonds between rigid sections. Since the magnitude of the C_α_-C_α_ distance fluctuation correlates with the PAE score, the PAE matrix can be used to scale the force constants of the individual bonds. Our approach allows us to optimize an ENM with easily accessible or available data without relying on, *e*.*g*., additional atomistic simulations ^49–51^.

While we have here tested AF-ENM in combination with the MARTIN3 coarse-grained force field 29, our approach could also increase the accuracy of other ENM-dependent cgMD models. Furthermore, our AF-ENM method could be used with small modifications to analytically determine effective protein dynamics via normal mode analysis ^59^ by primarily relying on the PAE scores to scale the force constants of the harmonic potentials. Although this approach would not be applicable for highly dynamic systems, it could nevertheless provide an improved description over conventional ENMs for rigid proteins. Similar to the AF-ENM framework, the AlphaFold scores could possibly be applied to derive Go-models ^40, 41^ instead of elastic networks. Hereby, the harmonic potentials would be replaced by van-der-Waals interactions, whose strength and occurrence would be controlled based on pLDDT and PAE scores. This approach could overcome the observed limitations in conditionally folded structural elements, such as the β8-strand of Hsp90^42^. The AF-ENM framework could thus open up new avenues to incorporate structural information into molecular dynamics simulations.

## Material and Methods

### MD simulations

Atomistic molecular dynamics (aMD) simulations were performed for 28 proteins (Figure 1), summarized in Table S1. The aMD simulations were performed with Gromacs ^60^ using the a99SB-disp force field ^43^ at T= 310 K, with a 2 fs timestep, and with the protein models embedded in a water-ion environment with 100 mM NaCl. The temperature and pressure were controlled with the velocity rescaling thermostat ^61^ and Parrinello-Rahman barostat ^62^. The two-domain proteins (1-23 and 27) were simulated for 2 μs, whereas the single-domain constructs (24-26) were simulated for 1 μs. For model 28, the NTD-MD construct of heat shock protein-90 (Hsp90), we used a 10.5 μs ensemble from our previous study ^42^, which was combined from 26 individual a99SB-disp simulations and reweighted with SAXS data based on a Bayesian/maximum entropy approach ^63^ (see reference ^42^ for further details).

Coarse-grained MD (cgMD) simulations (Table S4) were created based on the atomistic models using the MARTINI3 coarse-grained force field ^29^. The cgMD models were embedded in a 100 mM NaCl solution. The simulations were performed at T=310 K in an NPT ensemble with the velocity rescaling thermostat ^61^ and Parrinello-Rahman barostat ^62^ using Gromacs ^60^ and 20 fs timesteps. The cgMD simulation was performed with different ENMs: As reference ENMs, we used a cutoff distance of 0.9 nm with 500 kJ / (mol · nm^2^). For our newly introduced AF-ENM approach, we used force constants calculated with equation [6] or with twice as large ken values (AF-ENMx2). Additional simulations were performed with AF-ENM, and an extra 6% increased protein-water (P-W) interaction (AF-ENM+6%). For the single-domain systems, cgMD simulations were performed with the standard ENM, AF-ENM, AF-ENMx2 for 10 μs each. For linear Ub_2_ and the NTD-MD construct, we run 20 μs simulations with the standard ENM, AF-ENM, and AF-ENM+6%.

## ASSOCIATED CONTENT

### Supporting Information

Detailed comparison between the AlphaFold statistical scores and molecular dynamics simulations of the individual system; Comparisons between aMD and cgMD simulations (PDF).

## AUTHOR INFORMATION

### Notes

The simulation trajectories can be found in Zenodo, 10.5281/zenodo.7212856.

## ACKNOWLEDGMENT

This project was supported by the Knut and Allice Wallenberg Foundation (2019.0043 and 2019.0251) and Cancerfonden (pj200968). We are thankful for the computing time provided by SuperMuc at the Leibniz Rechenzentrum (project: pn98ha), and Swedish National Infrastructure for Computing (SNIC 2022/1-29, LUMI project id: 465000179).

## REFERENCES

(1) Jumper, J.; Evans, R.; Pritzel, A.; Green, T.; Figurnov, M.; Ronneberger, O.; Tunyasuvunakool, K.; Bates, R.; Zidek, A.; Potapenko, A.; et al. Highly accurate protein structure prediction with AlphaFold. Nature 2021, 596 (7873), 583–589. DOI: 10.1038/s41586-021-03819-2.

(2) Baek, M.; DiMaio, F.; Anishchenko, I.; Dauparas, J.; Ovchinnikov, S.; Lee, G. R.; Wang, J.; Cong, Q.; Kinch, L. N.; Schaeffer, R. D.; et al. Accurate prediction of protein structures and interactions using a three-track neural network. Science 2021, 373 (6557), 871–876. DOI: 10.1126/science.abj8754.

(3) Varadi, M.; Anyango, S.; Deshpande, M.; Nair, S.; Natassia, C.; Yordanova, G.; Yuan, D.; Stroe, O.; Wood, G.; Laydon, A.; et al. AlphaFold Protein Structure Database: massively expanding the structural coverage of protein-sequence space with high-accuracy models. Nucleic Acids Res 2022, 50 (D1), D439–D444. DOI: 10.1093/nar/gkab1061.

(4) Callaway, E. ‘The entire protein universe’: AI predicts shape of nearly every known protein. Nature 2022, 608 (7921), 15–16. DOI: 10.1038/d41586-022-02083-2.

(5) Mirdita, M.; Schütze, K.; Moriwaki, Y.; Heo, L.; Ovchinnikov, S.; Steinegger, M. ColabFold - Making protein folding accessible to all. bioRxiv 2022, 2021.2008.2015.456425. DOI: 10.1101/2021.08.15.456425.

(6) Steinegger, M.; Soding, J. MMseqs2 enables sensitive protein sequence searching for the analysis of massive data sets. Nat Biotechnol 2017, 35 (11), 1026–1028. DOI: 10.1038/nbt.3988.

(7) Evans, R.; O’Neill, M.; Pritzel, A.; Antropova, N.; Senior, A.; Green, T.; Žídek, A.; Bates, R.; Blackwell, S.; Yim, J.; et al. Protein complex prediction with AlphaFold-Multimer. bioRxiv 2022, 2021.2010.2004.463034. DOI: 10.1101/2021.10.04.463034.

(8) Bryant, P.; Pozzati, G.; Zhu, W.; Shenoy, A.; Kundrotas, P.; Elofsson, A. Predicting the structure of large protein complexes using AlphaFold and Monte Carlo tree search. Nat Commun 2022, 13 (1), 6028. DOI: 10.1038/s41467-022-33729-4.

(9) Mariani, V.; Biasini, M.; Barbato, A.; Schwede, T. lDDT: a local superposition-free score for comparing protein structures and models using distance difference tests. Bioinformatics 2013, 29 (21), 2722–2728. DOI: 10.1093/bioinformatics/btt473.

(10) Tunyasuvunakool, K.; Adler, J.; Wu, Z.; Green, T.; Zielinski, M.; Zidek, A.; Bridgland, A.; Cowie, A.; Meyer, C.; Laydon, A.; et al. Highly accurate protein structure prediction for the human proteome. Nature 2021, 596 (7873), 590–596. DOI: 10.1038/s41586-021-03828-1.

(11) Akdel, M.; Pires, D. E. V.; Porta Pardo, E.; Jänes, J.; Zalevsky, A. O.; Mészáros, B.; Bryant, P.; Good, L. L.; Laskowski, R. A.; Pozzati, G.; et al. A structural biology community assessment of AlphaFold 2 applications. bioRxiv 2021, 2021.2009.2026.461876. DOI: 10.1101/2021.09.26.461876.

(12) Ruff, K. M.; Pappu, R. V. AlphaFold and Implications for Intrinsically Disordered Proteins. J Mol Biol 2021, 433 (20), 167208. DOI: 10.1016/j.jmb.2021.167208.

(13) Binder, J. L.; Berendzen, J.; Stevens, A. O.; He, Y.; Wang, J.; Dokholyan, N. V.; Oprea, T. I. AlphaFold illuminates half of the dark human proteins. Curr Opin Struct Biol 2022, 74, 102372. DOI: 10.1016/j.sbi.2022.102372.

(14) Alderson, T. R.; Pritišanac, I.; Moses, A. M.; Forman-Kay, J. D. Systematic identification of conditionally folded intrinsically disordered regions by AlphaFold2. bioRxiv 2022, 2022.2002.2018.481080. DOI: 10.1101/2022.02.18.481080.

(15) Piovesan, D.; Monzon, A. M.; Tosatto, S. C. E. Intrinsic Protein Disorder, Conditional Folding and AlphaFold2. bioRxiv 2022, 2022.2003.2003.482768. DOI: 10.1101/2022.03.03.482768.

(16) Zhang, Y.; Skolnick, J. Scoring function for automated assessment of protein structure template quality. Proteins 2004, 57 (4), 702–710. DOI: 10.1002/prot.20264.

(17) Zemla, A. LGA: A method for finding 3D similarities in protein structures. Nucleic Acids Res 2003, 31 (13), 3370–3374. DOI: 10.1093/nar/gkg571.

(18) Jendrusch, M.; Korbel, J. O.; Sadiq, S. K. AlphaDesign: A de novo protein design framework based on AlphaFold. bioRxiv 2021, 2021.2010.2011.463937. DOI: 10.1101/2021.10.11.463937.

(19) Fowler, N. J.; Williamson, M. P. The accuracy of protein structures in solution determined by AlphaFold and NMR. Structure 2022, 30 (7), 925–933 e922. DOI: 10.1016/j.str.2022.04.005.

(20) Del Alamo, D.; Sala, D.; McHaourab, H. S.; Meiler, J. Sampling alternative conformational states of transporters and receptors with AlphaFold2. Elife 2022, 11. DOI: 10.7554/eLife.75751.

(21) Zuckerman, D. M. Equilibrium sampling in biomolecular simulations. Annu Rev Biophys 2011, 40, 41-62. DOI: 10.1146/annurev-biophys-042910-155255.

(22) Camilloni, C.; Pietrucci, F. Advanced simulation techniques for the thermodynamic and kinetic characterization of biological systems. Advances in Physics: X 2018, 3 (1). DOI: 10.1080/23746149.2018.1477531.

(23) Periole, X.; Cavalli, M.; Marrink, S. J.; Ceruso, M. A. Combining an Elastic Network With a Coarse-Grained Molecular Force Field: Structure, Dynamics, and Intermolecular Recognition. J Chem Theory Comput 2009, 5 (9), 2531–2543. DOI: 10.1021/ct9002114.

(24) Monticelli, L.; Kandasamy, S. K.; Periole, X.; Larson, R. G.; Tieleman, D. P.; Marrink, S. J. The MARTINI Coarse-Grained Force Field: Extension to Proteins. J Chem Theory Comput 2008, 4 (5), 819–834. DOI: 10.1021/ct700324x.

(25) Tirion, M. M. Large Amplitude Elastic Motions in Proteins from a Single-Parameter, Atomic Analysis. Phys Rev Lett 1996, 77 (9), 1905–1908. DOI: 10.1103/PhysRevLett.77.1905.

(26) Kmiecik, S.; Gront, D.; Kolinski, M.; Wieteska, L.; Dawid, A. E.; Kolinski, A. Coarse-Grained Protein Models and Their Applications. Chem Rev 2016, 116 (14), 7898–7936. DOI: 10.1021/acs.chemrev.6b00163.

(27) Marrink, S. J.; Risselada, H. J.; Yefimov, S.; Tieleman, D. P.; de Vries, A. H. The MARTINI force field: coarse grained model for biomolecular simulations. J Phys Chem B 2007, 111 (27), 7812–7824. DOI: 10.1021/jp071097f.

(28) de Jong, D. H.; Singh, G.; Bennett, W. F.; Arnarez, C.; Wassenaar, T. A.; Schafer, L. V.; Periole, X.; Tieleman, D. P.; Marrink, S. J. Improved Parameters for the Martini Coarse-Grained Protein Force Field. J Chem Theory Comput 2013, 9 (1), 687–697. DOI: 10.1021/ct300646g.

(29) Souza, P. C. T.; Alessandri, R.; Barnoud, J.; Thallmair, S.; Faustino, I.; Grunewald, F.; Patmanidis, I.; Abdizadeh, H.; Bruininks, B. M. H.; Wassenaar, T. A.; et al. Martini 3: a general purpose force field for coarse-grained molecular dynamics. Nat Methods 2021, 18 (4), 382–388. DOI: 10.1038/s41592-021-01098-3.

(30) Marrink, S. J.; Tieleman, D. P. Perspective on the Martini model. Chem Soc Rev 2013, 42 (16), 6801–6822. DOI: 10.1039/c3cs60093a.

(31) Uusitalo, J. J.; Ingolfsson, H. I.; Akhshi, P.; Tieleman, D. P.; Marrink, S. J. Martini Coarse-Grained Force Field: Extension to DNA. J Chem Theory Comput 2015, 11 (8), 3932–3945. DOI: 10.1021/acs.jctc.5b00286.

(32) Uusitalo, J. J.; Ingolfsson, H. I.; Marrink, S. J.; Faustino, I. Martini Coarse-Grained Force Field: Extension to RNA. Biophys J 2017, 113 (2), 246–256. DOI: 10.1016/j.bpj.2017.05.043.

(33) Lopez, C. A.; Rzepiela, A. J.; de Vries, A. H.; Dijkhuizen, L.; Hunenberger, P. H.; Marrink, S. J. Martini Coarse-Grained Force Field: Extension to Carbohydrates. J Chem Theory Comput 2009, 5 (12), 3195–3210. DOI: 10.1021/ct900313w.

(34) Alessandri, R.; Barnoud, J.; Gertsen, A.; Patmanidis, I.; Vries, A.; Telles de Souza, P. C.; Marrink, S. Martini 3 Coarse Grained Force Field: Small Molecules. Advanced Theory and Simulations 2021, 5, 2100391. DOI: 10.1002/adts.202100391.

(35) Stark, A. C.; Andrews, C. T.; Elcock, A. H. Toward optimized potential functions for protein-protein interactions in aqueous solutions: osmotic second virial coefficient calculations using the MARTINI coarse-grained force field. J Chem Theory Comput 2013, 9 (9). DOI: 10.1021/ct400008p.

(36) Javanainen, M.; Martinez-Seara, H.; Vattulainen, I. Excessive aggregation of membrane proteins in the Martini model. PLoS One 2017, 12 (11), e0187936. DOI: 10.1371/journal.pone.0187936.

(37) Larsen, A. H.; Wang, Y.; Bottaro, S.; Grudinin, S.; Arleth, L.; Lindorff-Larsen, K. Combining molecular dynamics simulations with small-angle X-ray and neutron scattering data to study multi-domain proteins in solution. PLoS Comput Biol 2020, 16 (4), e1007870. DOI: 10.1371/journal.pcbi.1007870.

(38) Jussupow, A.; Messias, A. C.; Stehle, R.; Geerlof, A.; Solbak, S. M. O.; Paissoni, C.; Bach, A.; Sattler, M.; Camilloni, C. The dynamics of linear polyubiquitin. Sci Adv 2020, 6 (42), eabc3786. DOI: 10.1126/sciadv.abc3786.

(39) Thomasen, F. E.; Pesce, F.; Roesgaard, M. A.; Tesei, G.; Lindorff-Larsen, K. Improving Martini 3 for Disordered and Multidomain Proteins. J Chem Theory Comput 2022, 2021.2010.2001.462803. DOI: 10.1021/acs.jctc.1c01042.

(40) Sulkowska, J. I.; Cieplak, M. Selection of optimal variants of Go-like models of proteins through studies of stretching. Biophys J 2008, 95 (7), 3174–3191. DOI: 10.1529/biophysj.107.127233.

(41) Poma, A. B.; Cieplak, M.; Theodorakis, P. E. Combining the MARTINI and Structure-Based Coarse-Grained Approaches for the Molecular Dynamics Studies of Conformational Transitions in Proteins. J Chem Theory Comput 2017, 13 (3), 1366–1374. DOI: 10.1021/acs.jctc.6b00986.

(42) Jussupow, A.; Lopez, A.; Baumgart, M.; Mader, S. L.; Sattler, M.; Kaila, V. R. I. Extended conformational states dominate the Hsp90 chaperone dynamics. J Biol Chem 2022, 102101. DOI: 10.1016/j.jbc.2022.102101.

(43) Robustelli, P.; Piana, S.; Shaw, D. E. Developing a molecular dynamics force field for both folded and disordered protein states. Proc Natl Acad Sci U S A 2018, 115 (21), E4758–E4766. DOI: 10.1073/pnas.1800690115.

(44) Tsutsumi, S.; Mollapour, M.; Graf, C.; Lee, C. T.; Scroggins, B. T.; Xu, W.; Haslerova, L.; Hessling, M.; Konstantinova, A. A.; Trepel, J. B.; et al. Hsp90 charged-linker truncation reverses the functional consequences of weakened hydrophobic contacts in the N domain. Nat Struct Mol Biol 2009, 16 (11), 1141–1147. DOI: 10.1038/nsmb.1682.

(45) Tsutsumi, S.; Mollapour, M.; Prodromou, C.; Lee, C. T.; Panaretou, B.; Yoshida, S.; Mayer, M. P.; Neckers, L. M. Charged linker sequence modulates eukaryotic heat shock protein 90 (Hsp90) chaperone activity. Proc Natl Acad Sci U S A 2012, 109 (8), 2937–2942. DOI: 10.1073/pnas.1114414109.

(46) Lorenz, O. R.; Freiburger, L.; Rutz, D. A.; Krause, M.; Zierer, B. K.; Alvira, S.; Cuellar, J.; Valpuesta, J. M.; Madl, T.; Sattler, M.; et al. Modulation of the Hsp90 chaperone cycle by a stringent client protein. Mol Cell 2014, 53 (6), 941–953. DOI: 10.1016/j.molcel.2014.02.003.

(47) Mader, S. L.; Lopez, A.; Lawatscheck, J.; Luo, Q.; Rutz, D. A.; Gamiz-Hernandez, A. P.; Sattler, M.; Buchner, J.; Kaila, V. R. I. Conformational dynamics modulate the catalytic activity of the molecular chaperone Hsp90. Nat Commun 2020, 11 (1), 1410. DOI: 10.1038/s41467-020-15050-0.

(48) López, A.; Elimelech, A. R.; Klimm, K.; Sattler, M. The Charged Linker Modulates the Conformations and Molecular Interactions of Hsp90. ChemBioChem 2021, 22 (6), 1084–1092 DOI: 10.1002/cbic.202000699.

(49) Kanada, R.; Terayama, K.; Tokuhisa, A.; Matsumoto, S.; Okuno, Y. Enhanced Conformational Sampling with an Adaptive Coarse-Grained Elastic Network Model Using Short-Time All-Atom Molecular Dynamics. J Chem Theory Comput 2022. DOI: 10.1021/acs.jctc.1c01074.

(50) Romo, T. D.; Grossfield, A. Validating and improving elastic network models with molecular dynamics simulations. Proteins 2011, 79 (1), 23–34. DOI: 10.1002/prot.22855.

(51) Ahmed, A.; Villinger, S.; Gohlke, H. Large-scale comparison of protein essential dynamics from molecular dynamics simulations and coarse-grained normal mode analyses. Proteins 2010, 78 (16), 3341–3352. DOI: 10.1002/prot.22841.

(52) Thach, T. T.; Shin, D.; Han, S.; Lee, S. New conformations of linear polyubiquitin chains from crystallographic and solution-scattering studies expand the conformational space of polyubiquitin. Acta Crystallogr D Struct Biol 2016, 72 (Pt 4), 524–535. DOI: 10.1107/S2059798316001510.

(53) Rohaim, A.; Kawasaki, M.; Kato, R.; Dikic, I.; Wakatsuki, S. Structure of a compact conformation of linear diubiquitin. Acta Crystallogr D Biol Crystallogr 2012, 68 (Pt 2), 102–108. DOI: 10.1107/S0907444911051195.

(54) Komander, D.; Reyes-Turcu, F.; Licchesi, J. D.; Odenwaelder, P.; Wilkinson, K. D.; Barford, D. Molecular discrimination of structurally equivalent Lys 63-linked and linear polyubiquitin chains. EMBO Rep 2009, 10 (5), 466–473. DOI: 10.1038/embor.2009.55.

(55) David, C. C.; Jacobs, D. J. Principal component analysis: a method for determining the essential dynamics of proteins. Methods Mol Biol 2014, 1084, 193–226. DOI: 10.1007/978-1-62703-658-0_11.

(56) Pearson, K. L III. On lines and planes of closest fit to systems of points in space. The London, Edinburgh, and Dublin Philosophical Magazine and Journal of Science 1901, 2 (11), 559–572. DOI: 10.1080/14786440109462720.

(57) Kullback, S.; Leibler, R. A. On Information and Sufficiency. The Annals of Mathematical Statistics 1951, 22 (1), 79–86. DOI: 10.1214/aoms/1177729694.

(58) Perez-Cruz, F. Kullback-Leibler Divergence Estimation of Continuous Distributions. Ieee Int Symp Info 2008, 1666–1670. DOI: Doi 10.1109/Isit.2008.4595271.

(59) Bahar, I.; Atilgan, A. R.; Erman, B. Direct evaluation of thermal fluctuations in proteins using a single-parameter harmonic potential. Folding and Design 1997, 2 (3), 173–181. DOI: 10.1016/s1359-0278(97)00024-2.

(60) Abraham, M. J.; Murtola, T.; Schulz, R.; Páll, S.; Smith, J. C.; Hess, B.; Lindahl, E. GROMACS: High performance molecular simulations through multi-level parallelism from laptops to supercomputers. SoftwareX 2015, 1-2, 19–25. DOI: 10.1016/j.softx.2015.06.001.

(61) Bussi, G.; Donadio, D.; Parrinello, M. Canonical sampling through velocity rescaling. J Chem Phys 2007, 126 (1), 014101. DOI: 10.1063/1.2408420.

(62) Parrinello, M.; Rahman, A. Polymorphic transitions in single crystals: A new molecular dynamics method. Journal of Applied Physics 1981, 52 (12), 7182–7190. DOI: 10.1063/1.328693.

(63) Bottaro, S.; Bengtsen, T.; Lindorff-Larsen, K. Integrating Molecular Simulation and Experimental Data: A Bayesian/Maximum Entropy Reweighting Approach. Methods Mol Biol 2020, 2112, 219–240. DOI: 10.1007/978-1-0716-0270-6_15.

