## Supplementary material for "Effective Molecular Dynamics from Neural-Network Based Structure Prediction Models": SI Appendix

### Effective Molecular Dynamics from AlphaFold

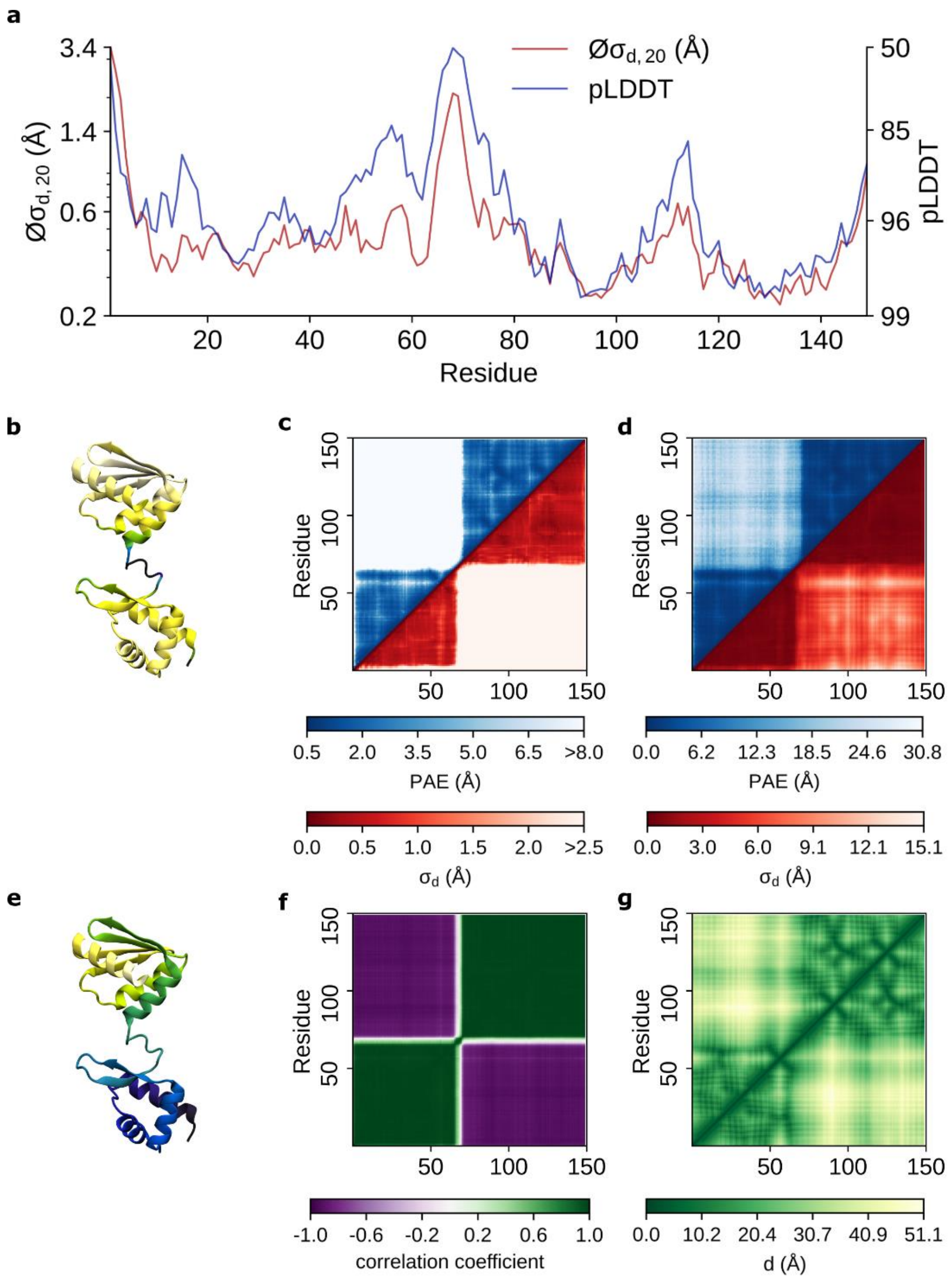


**Figure S1:** AlphaFold structure prediction vs. aMD of *arginine repressor* (protein 1) **a**) pLDDT scores vs. $Ø\sigma_{d,20}$ values **b**) The protein structure colored based on its pLDDT scores. The dark blue colors correspond to residues with a pLDDT score <= 60, while the white color corresponds to residues with a pLDDT score close to 100. **c**) Comparison between (symmetrized) PAE matrices (blue) against the standard deviation of all $C_\alpha$ distances $\sigma_d$ (red). The PAE scores range between 0.5 to 8.0 while the $\sigma_d$ are limited to <2.5 Å. **d**) Comparison between (symmetrized) PAE matrices (blue) against the standard deviation of all $C_\alpha$ distances $\sigma_d$ (red) for the maximum range. **e**) The protein structure colored based on its residue number. Dark blue colors correspond to low values, and yellows correspond to high values. **f**) Distance correlation matrix obtained from aMD simulations. **g**) Distance matrix obtained from aMD simulation.

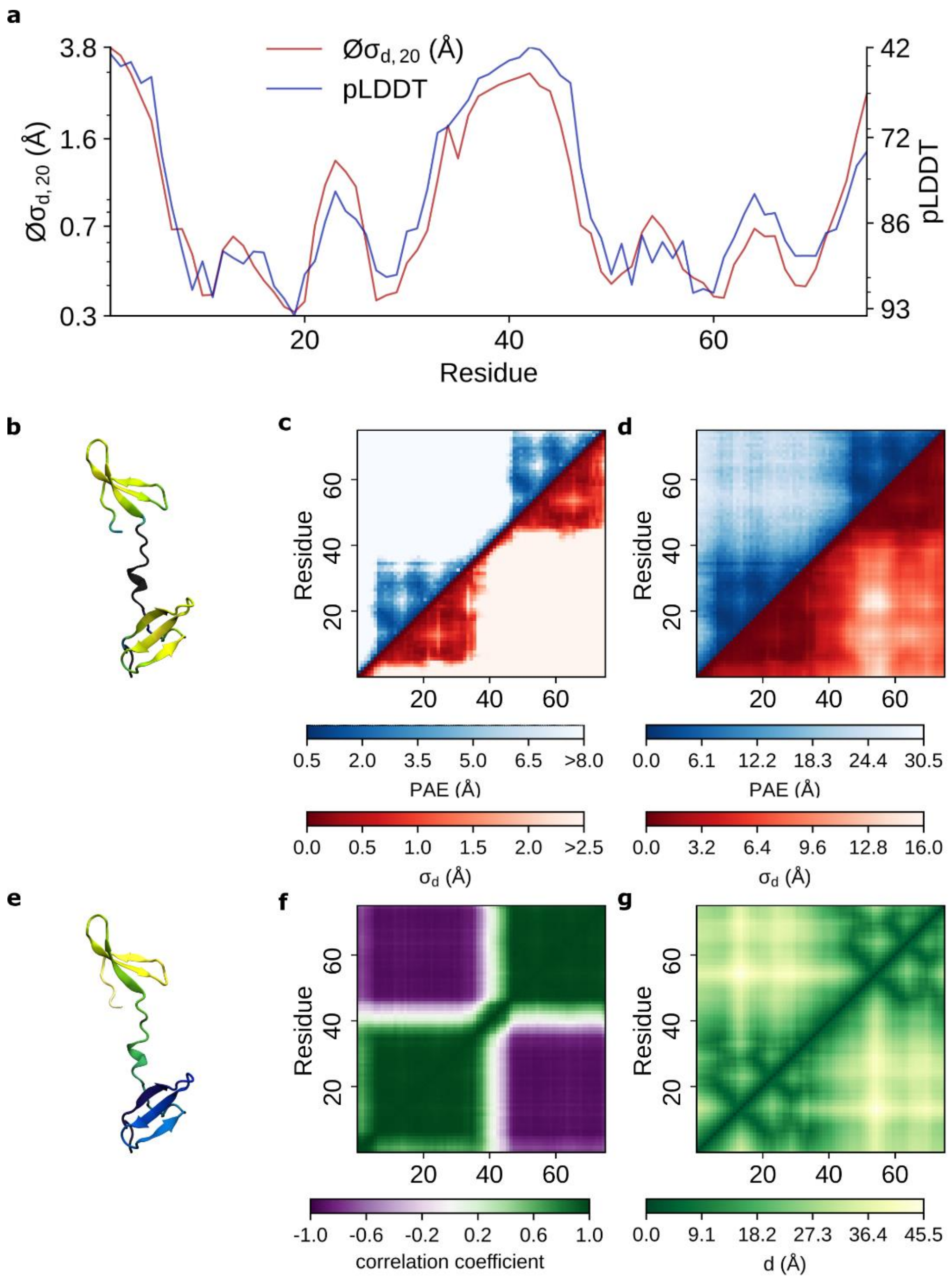


**Figure S2:** AlphaFold vs. aMD for *WW domain-binding protein 4* (protein 2) **a**) pLDDT scores vs. $Ø\sigma_{d,20}$ values **b**) The protein structure colored based on its pLDDT scores. The dark blue colors correspond to residues with a pLDDT score <= 60, while the white color corresponds to residues with a pLDDT score close to 100. **c**) Comparison between (symmetrized) PAE matrices (blue) against the standard deviation of all $C_\alpha$ distances $\sigma_d$ (red). The PAE scores range between 0.5 to 8.0 while the $\sigma_d$ are limited to <2.5 Å. **d**) Comparison between (symmetrized) PAE matrices (blue) against the standard deviation of all $C_\alpha$ distances $\sigma_d$ (red) for the maximum range. **e**) The protein structure colored based on its residue number. Dark blue colors correspond to low values, and yellows correspond to high values. **f**) Distance correlation matrix obtained from aMD simulations. **g**) Distance matrix obtained from aMD simulation.

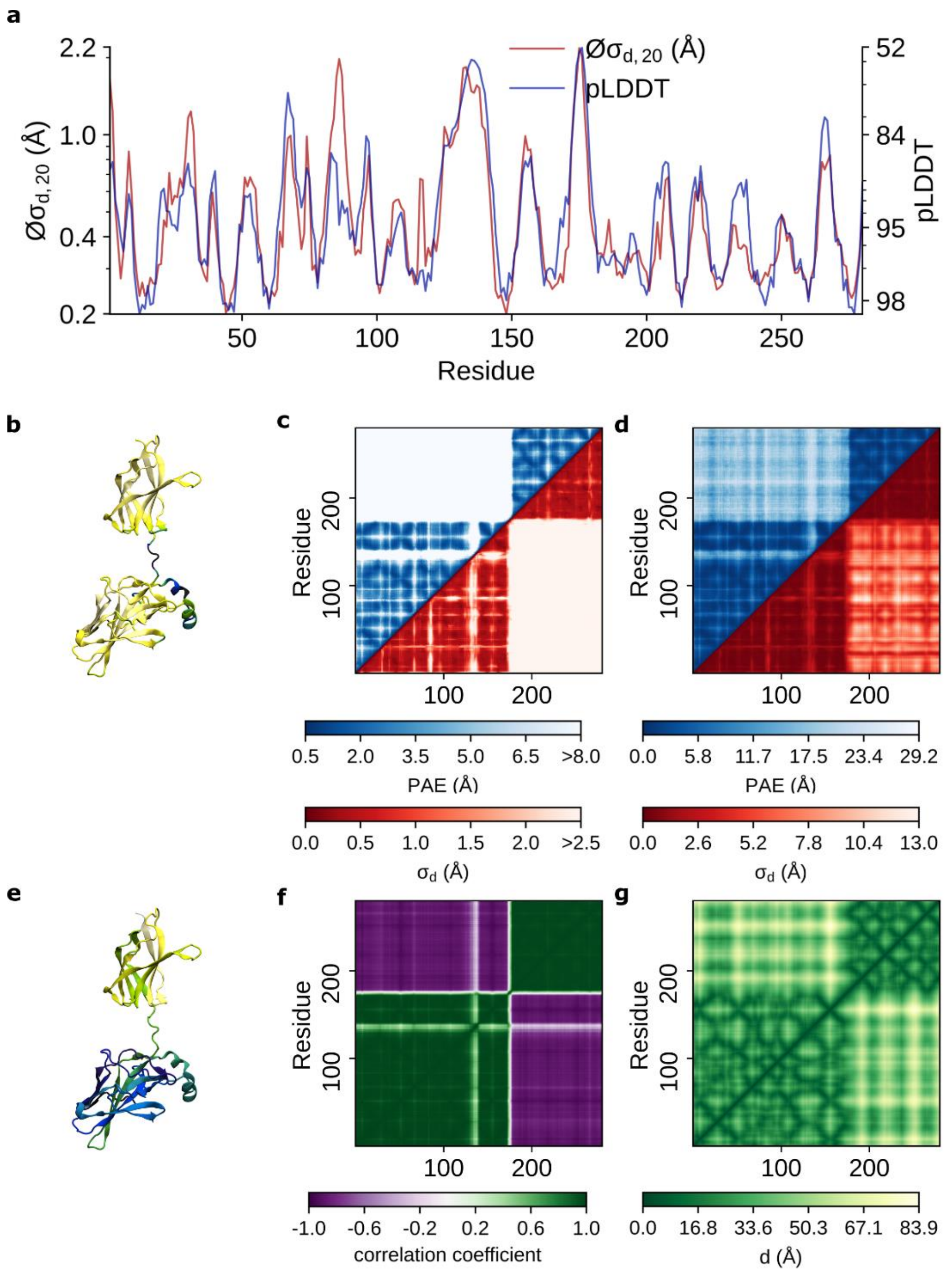


**Figure S3:** AlphaFold vs. aMD for *nuclear factor of activated T-cells, cytoplasmic 2* (protein 3) **a**) pLDDT scores vs. $Ø\sigma_{d,20}$ values **b**) The protein structure colored based on its pLDDT scores. The dark blue colors correspond to residues with a pLDDT score <= 60, while the white color corresponds to residues with a pLDDT score close to 100. **c**) Comparison between (symmetrized) PAE matrices (blue) against the standard deviation of all $C_\alpha$ distances $\sigma_d$ (red). The PAE scores range between 0.5 to 8.0 while the $\sigma_d$ are limited to <2.5 Å. **d**) Comparison between (symmetrized) PAE matrices (blue) against the standard deviation of all $C_\alpha$ distances $\sigma_d$ (red) for the maximum range. **e**) The protein structure colored based on its residue number. Dark blue colors correspond to low values, and yellows correspond to high values. **f**) Distance correlation matrix obtained from aMD simulations. **g**) Distance matrix obtained from aMD simulation.

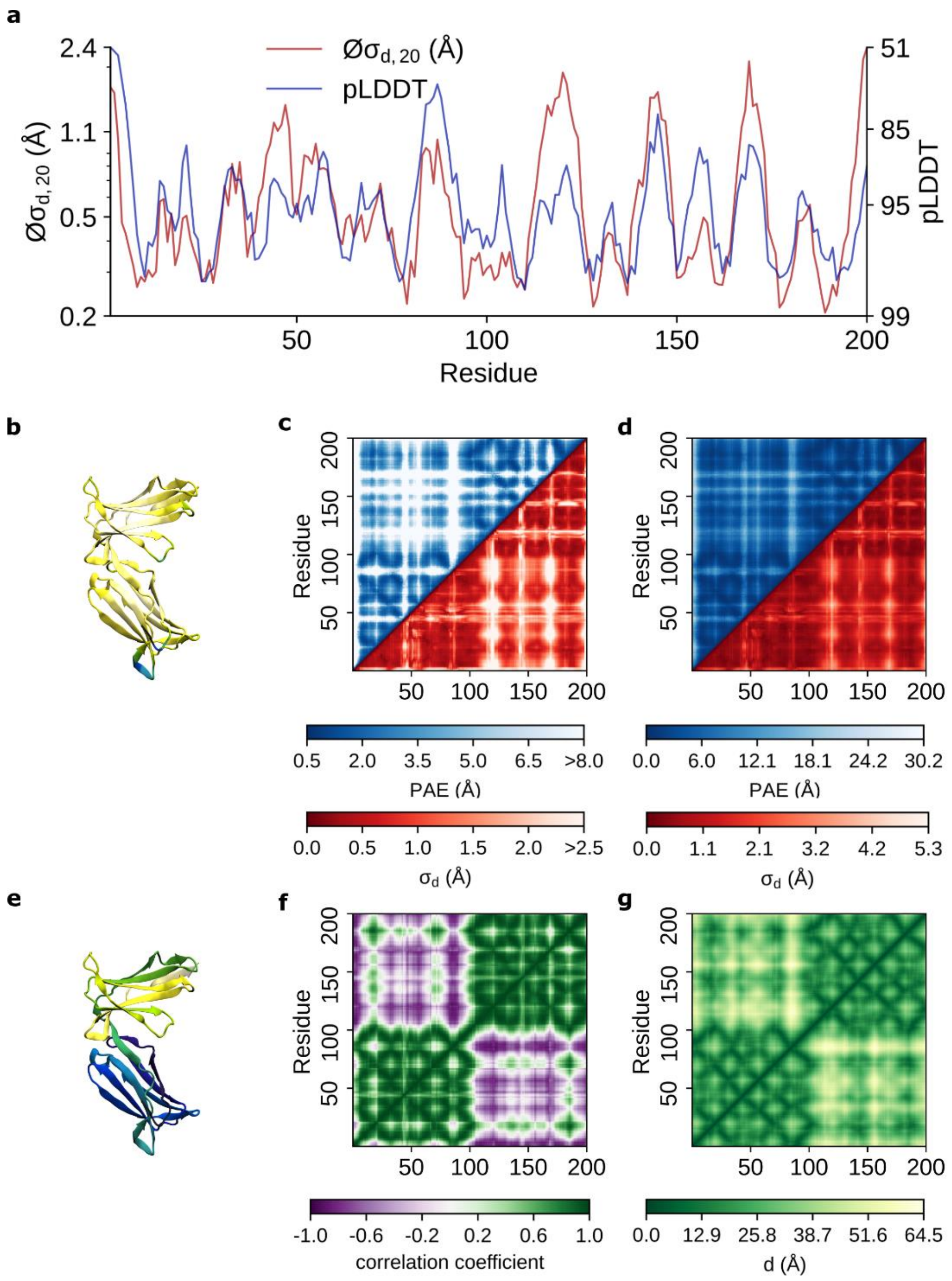


**Figure S4:** AlphaFold vs. aMD for *killer cell immunoglobulin-like receptor 2DL1* (protein 4) **a**) pLDDT scores vs. $Ø\sigma_{d,20}$ values **b**) The protein structure colored based on its pLDDT scores. The dark blue colors correspond to residues with a pLDDT score <= 60, while the white color corresponds to residues with a pLDDT score close to 100. **c**) Comparison between (symmetrized) PAE matrices (blue) against the standard deviation of all $C_\alpha$ distances $\sigma_d$ (red). The PAE scores range between 0.5 to 8.0 while the $\sigma_d$ are limited to <2.5 Å. **d**) Comparison between (symmetrized) PAE matrices (blue) against the standard deviation of all $C_\alpha$ distances $\sigma_d$ (red) for the maximum range. **e**) The protein structure colored based on its residue number. Dark blue colors correspond to low values, and yellows correspond to high values. **f**) Distance correlation matrix obtained from aMD simulations. **g**) Distance matrix obtained from aMD simulation.

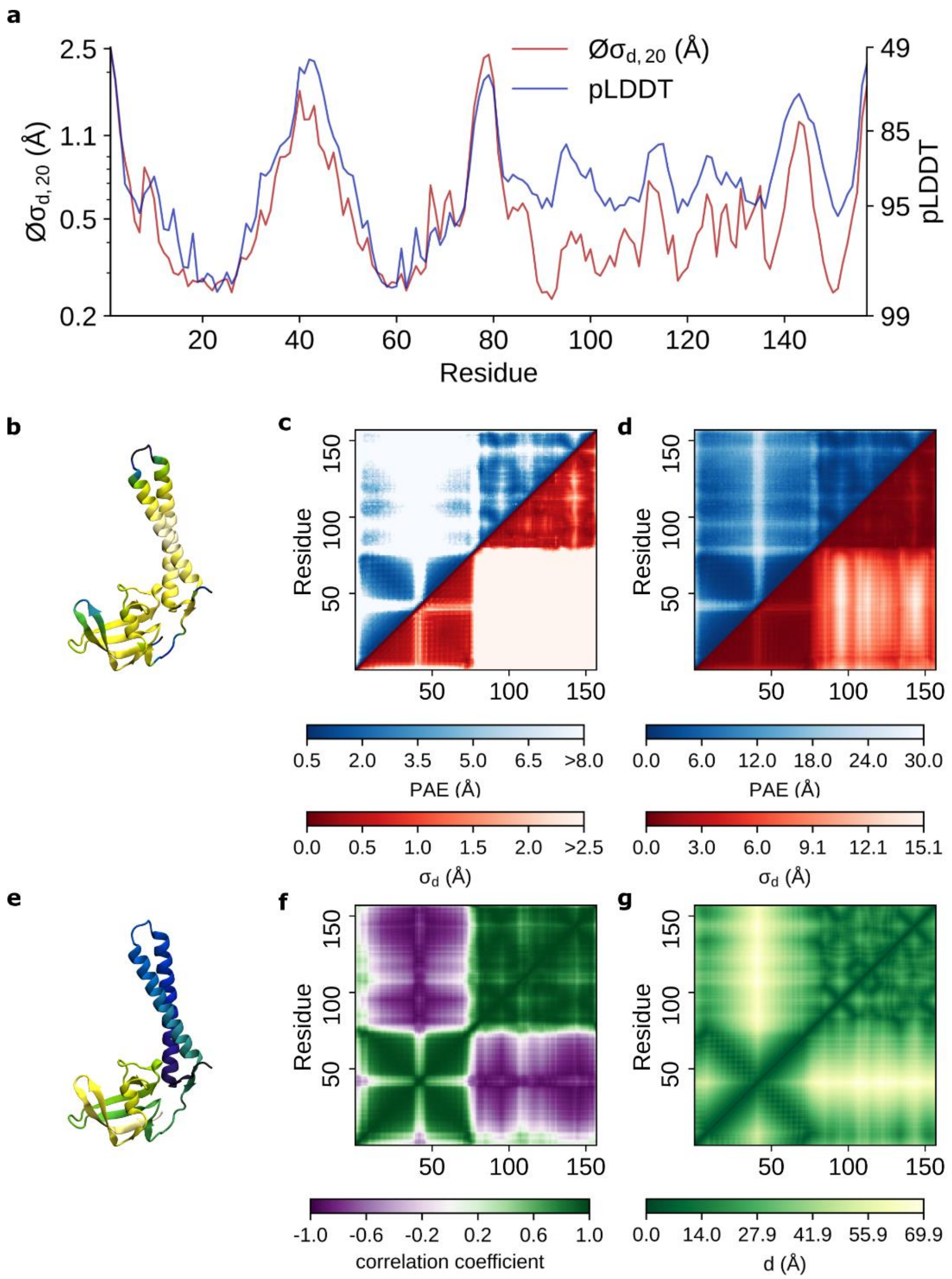


**Figure S5:** AlphaFold vs. aMD for *transcription inhibitor protein Gfh1* (protein 5) **a**) pLDDT scores vs. $Ø\sigma_{d,20}$ values **b**) The protein structure colored based on its pLDDT scores. The dark blue colors correspond to residues with a pLDDT score <= 60, while the white color corresponds to residues with a pLDDT score close to 100. **c**) Comparison between (symmetrized) PAE matrices (blue) against the standard deviation of all $C_\alpha$ distances $\sigma_d$ (red). The PAE scores range between 0.5 to 8.0 while the $\sigma_d$ are limited to <2.5 Å. **d**) Comparison between (symmetrized) PAE matrices (blue) against the standard deviation of all $C_\alpha$ distances $\sigma_d$ (red) for the maximum range. **e**) The protein structure colored based on its residue number. Dark blue colors correspond to low values, and yellows correspond to high values. **f**) Distance correlation matrix obtained from aMD simulations. **g**) Distance matrix obtained from aMD simulation.

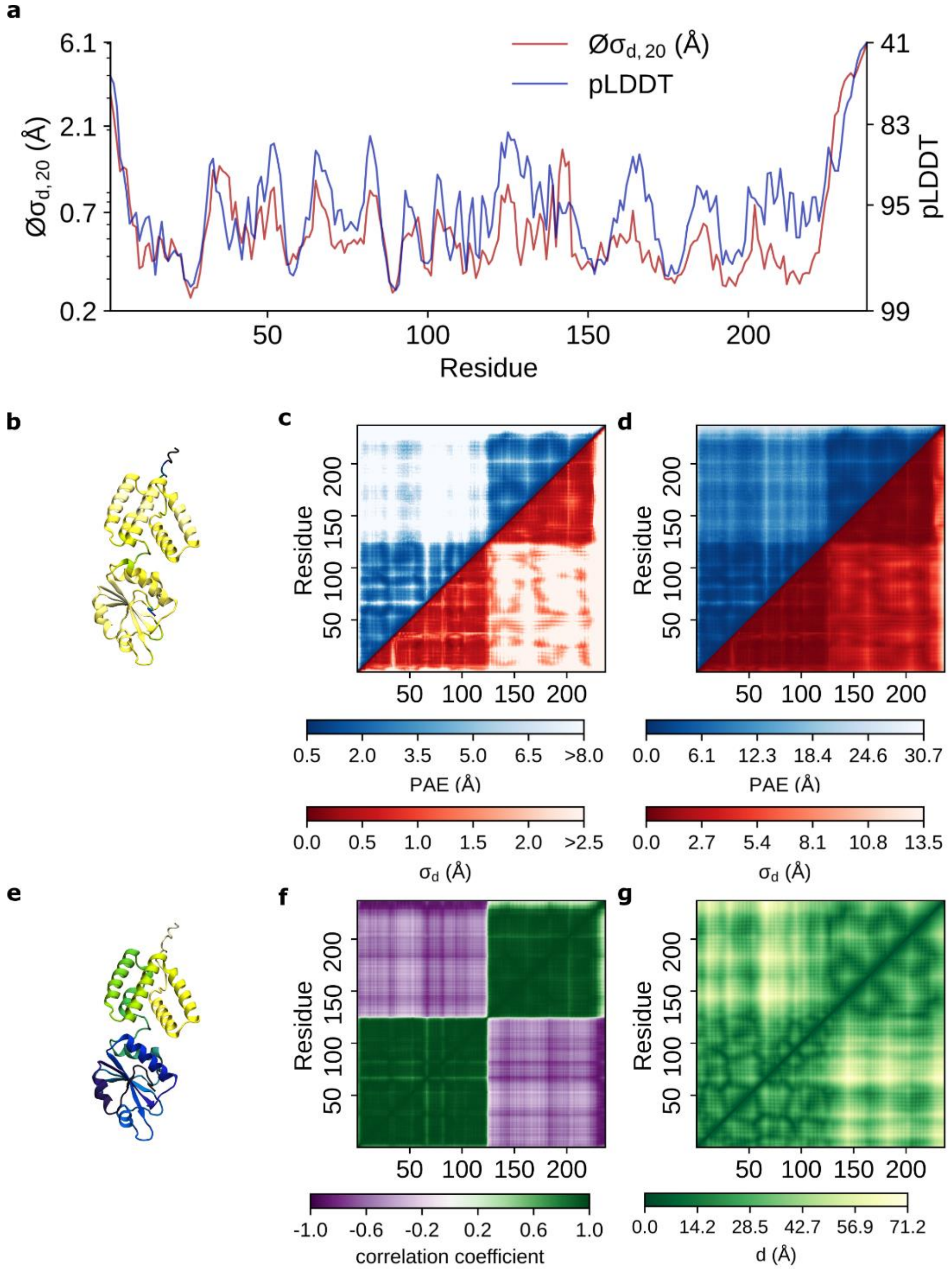


**Figure S6:** AlphaFold vs. aMD for *protein windbeutel* (protein 6) **a**) pLDDT scores vs. $Øσ_{d,20}$ values **b**) The protein structure colored based on its pLDDT scores. The dark blue colors correspond to residues with a pLDDT score <= 60, while the white color corresponds to residues with a pLDDT score close to 100. **c**) Comparison between (symmetrized) PAE matrices (blue) against the standard deviation of all $C_α$ distances $σ_d$ (red). The PAE scores range between 0.5 to 8.0 while the $σ_d$ are limited to <2.5 Å. **d**) Comparison between (symmetrized) PAE matrices (blue) against the standard deviation of all $C_α$ distances $σ_d$ (red) for the maximum range. **e**) The protein structure colored based on its residue number. Dark blue colors correspond to low values, and yellows correspond to high values. **f**) Distance correlation matrix obtained from aMD simulations. **g**) Distance matrix obtained from aMD simulation.

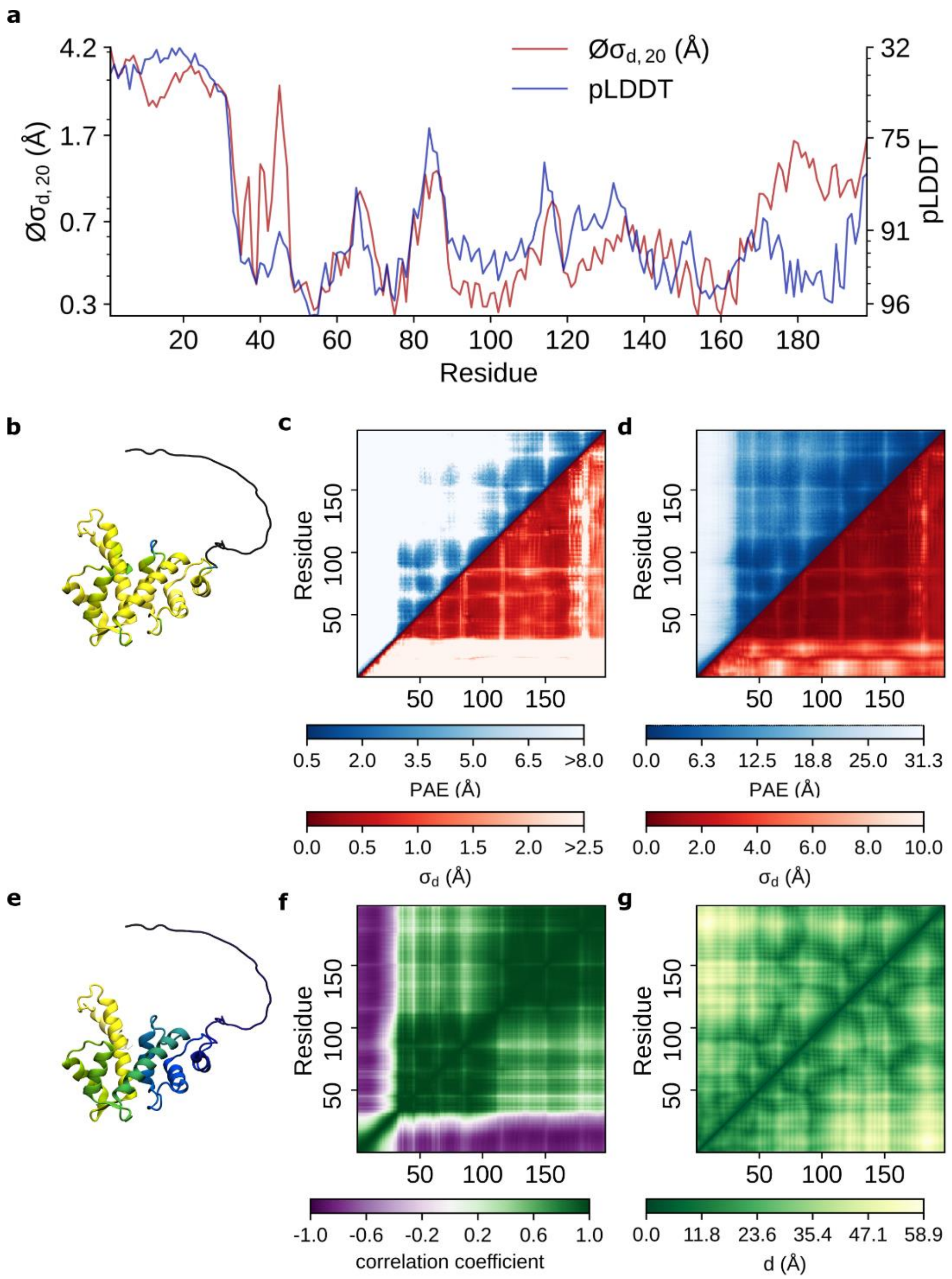


**Figure S7:** AlphaFold vs. *aMD for sorcin* (protein 7) **a**) pLDDT scores vs. $Ø\sigma_{d,20}$ values **b**) The protein structure colored based on its pLDDT scores. The dark blue colors correspond to residues with a pLDDT score <= 60, while the white color corresponds to residues with a pLDDT score close to 100. **c**) Comparison between (symmetrized) PAE matrices (blue) against the standard deviation of all $C_\alpha$ distances $\sigma_d$ (red). The PAE scores range between 0.5 to 8.0 while the $\sigma_d$ are limited to <2.5 Å. **d**) Comparison between (symmetrized) PAE matrices (blue) against the standard deviation of all $C_\alpha$ distances $\sigma_d$ (red) for the maximum range. **e**) The protein structure colored based on its residue number. Dark blue colors correspond to low values, and yellows correspond to high values. **f**) Distance correlation matrix obtained from aMD simulations. **g**) Distance matrix obtained from aMD simulation.

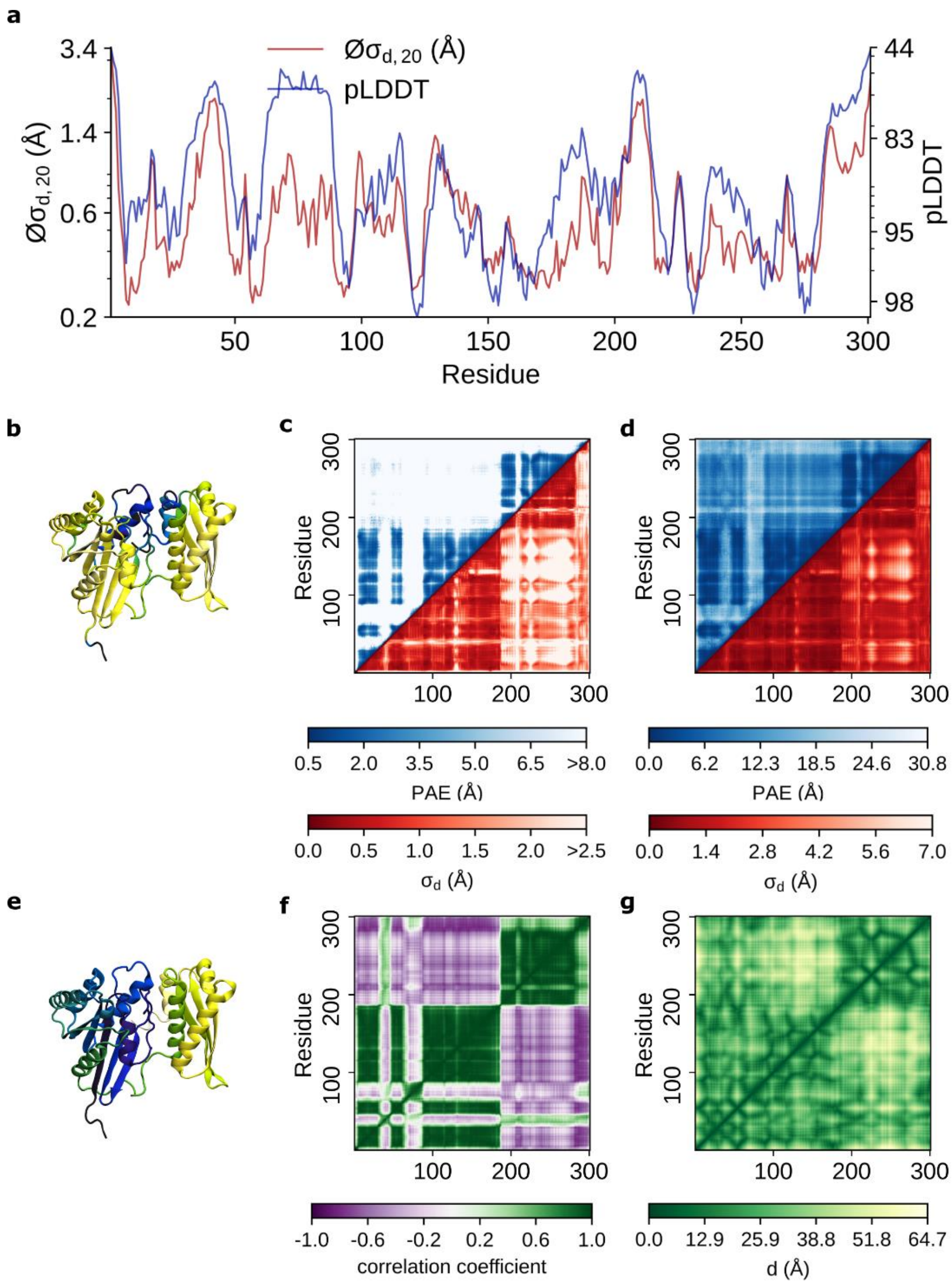


**Figure S8:** AlphaFold vs. aMD for *GTPase Era* (protein 8) **a**) pLDDT scores vs. $Ø\sigma_{d,20}$ values **b**) The protein structure colored based on its pLDDT scores. The dark blue colors correspond to residues with a pLDDT score <= 60, while the white color corresponds to residues with a pLDDT score close to 100. **c**) Comparison between (symmetrized) PAE matrices (blue) against the standard deviation of all $C_\alpha$ distances $\sigma_d$ (red). The PAE scores range between 0.5 to 8.0 while the $\sigma_d$ are limited to <2.5 Å. **d**) Comparison between (symmetrized) PAE matrices (blue) against the standard deviation of all $C_\alpha$ distances $\sigma_d$ (red) for the maximum range. **e**) The protein structure colored based on its residue number. Dark blue colors correspond to low values, and yellows correspond to high values. **f**) Distance correlation matrix obtained from aMD simulations. **g**) Distance matrix obtained from aMD simulation.

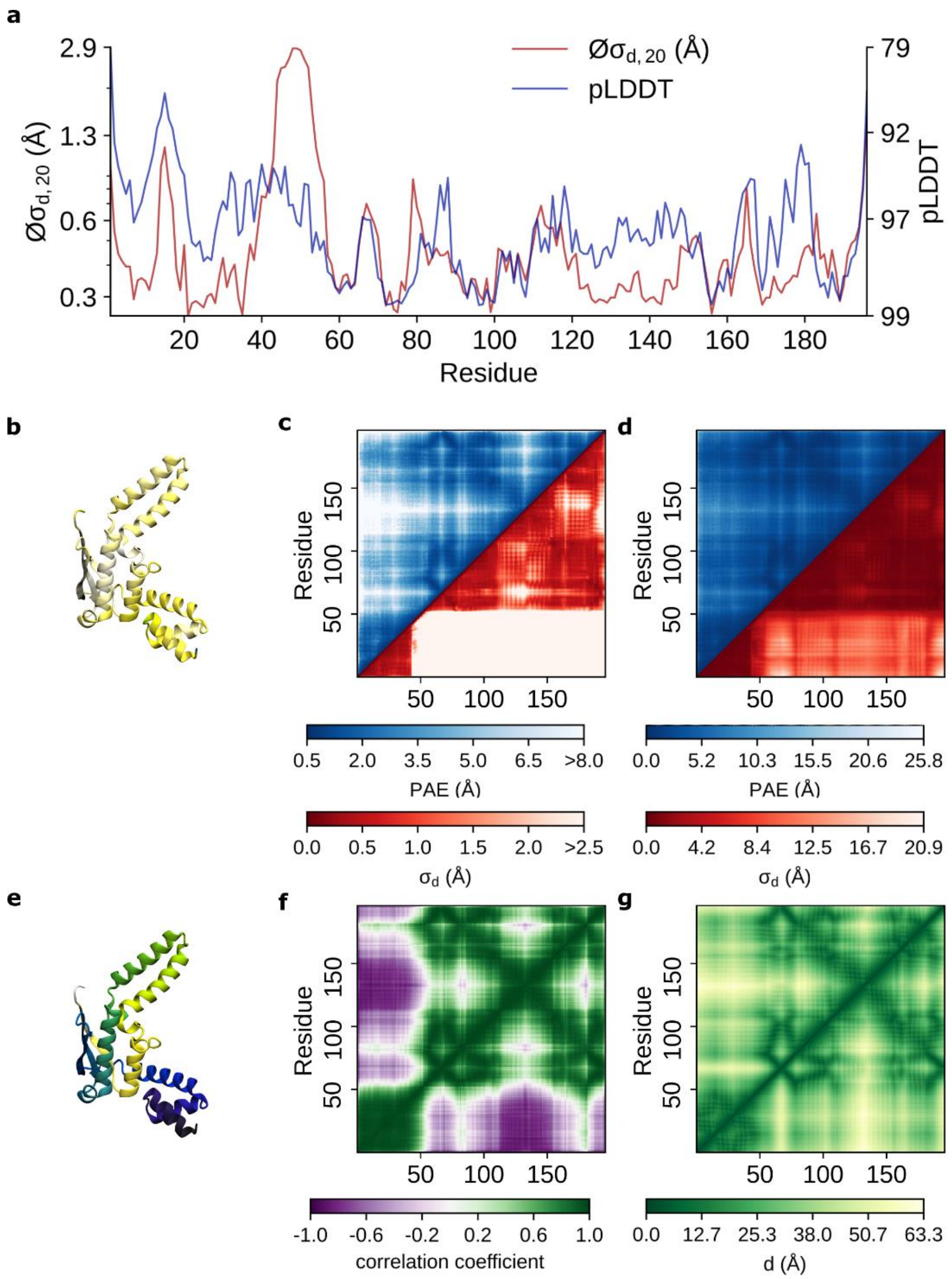


**Figure S9:** AlphaFold vs. aMD for *elongation factor T* (protein 9) **a**) pLDDT scores vs. $Ø\sigma_{d,20}$ values **b**) The protein structure colored based on its pLDDT scores. The dark blue colors correspond to residues with a pLDDT score <= 60, while the white color corresponds to residues with a pLDDT score close to 100. **c**) Comparison between (symmetrized) PAE matrices (blue) against the standard deviation of all $C_\alpha$ distances $\sigma_d$ (red). The PAE scores range between 0.5 to 8.0 while the $\sigma_d$ are limited to <2.5 Å. **d**) Comparison between (symmetrized) PAE matrices (blue) against the standard deviation of all $C_\alpha$ distances $\sigma_d$ (red) for the maximum range. **e**) The protein structure colored based on its residue number. Dark blue colors correspond to low values, and yellows correspond to high values. **f**) Distance correlation matrix obtained from aMD simulations. **g**) Distance matrix obtained from aMD simulation.

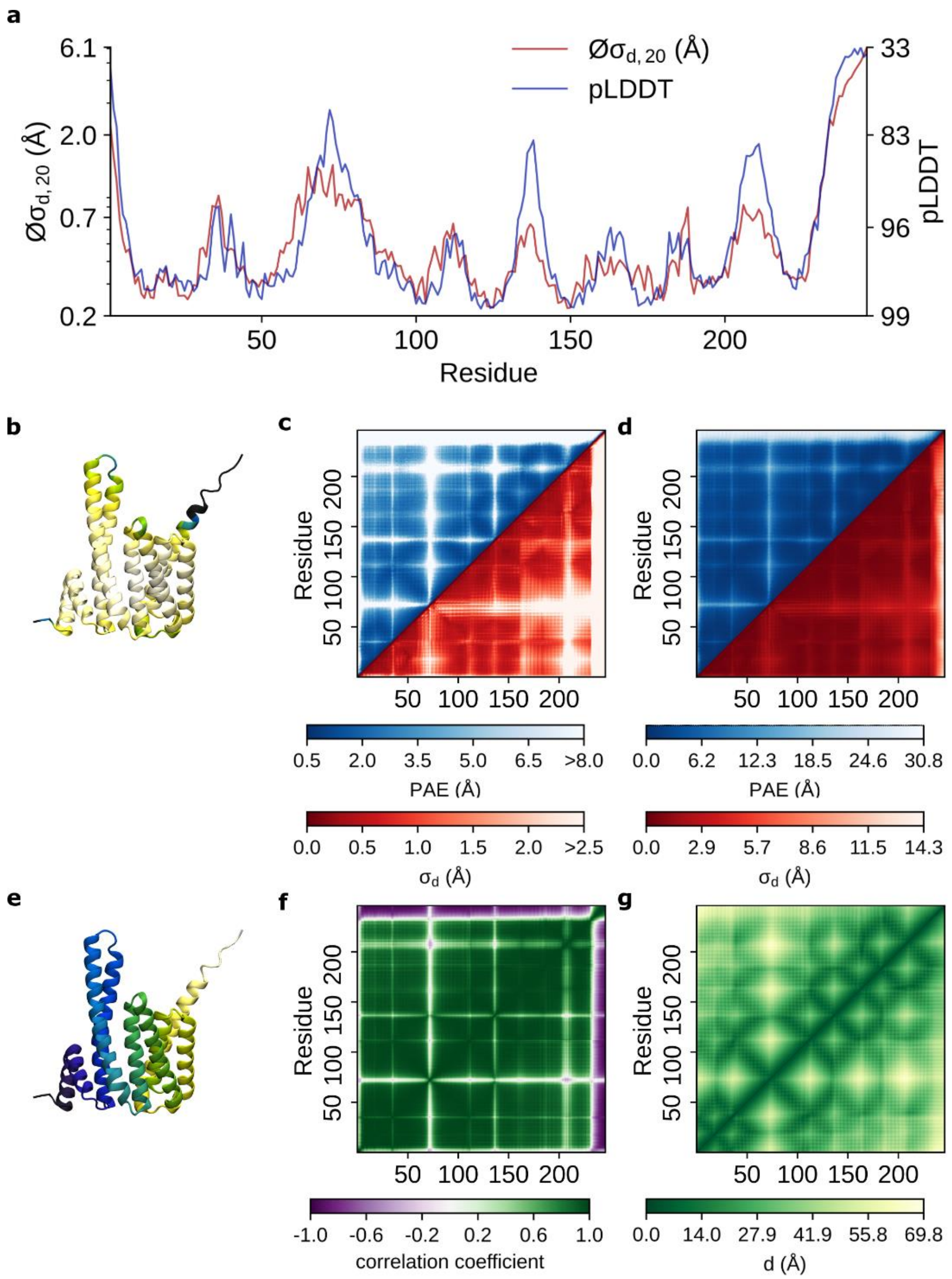


**Figure S10:** AlphaFold vs. aMD for *14-3-3 protein beta/alpha* (protein 10) **a**) pLDDT scores vs. $Ø\sigma_{d,20}$ values **b**) The protein structure colored based on its pLDDT scores. The dark blue colors correspond to residues with a pLDDT score <= 60, while the white color corresponds to residues with a pLDDT score close to 100. **c**) Comparison between (symmetrized) PAE matrices (blue) against the standard deviation of all $C_\alpha$ distances $\sigma_d$ (red). The PAE scores range between 0.5 to 8.0 while the $\sigma_d$ are limited to <2.5 Å. **d**) Comparison between (symmetrized) PAE matrices (blue) against the standard deviation of all $C_\alpha$ distances $\sigma_d$ (red) for the maximum range. **e**) The protein structure colored based on its residue number. Dark blue colors correspond to low values, and yellows correspond to high values. **f**) Distance correlation matrix obtained from aMD simulations. **g**) Distance matrix obtained from aMD simulation.

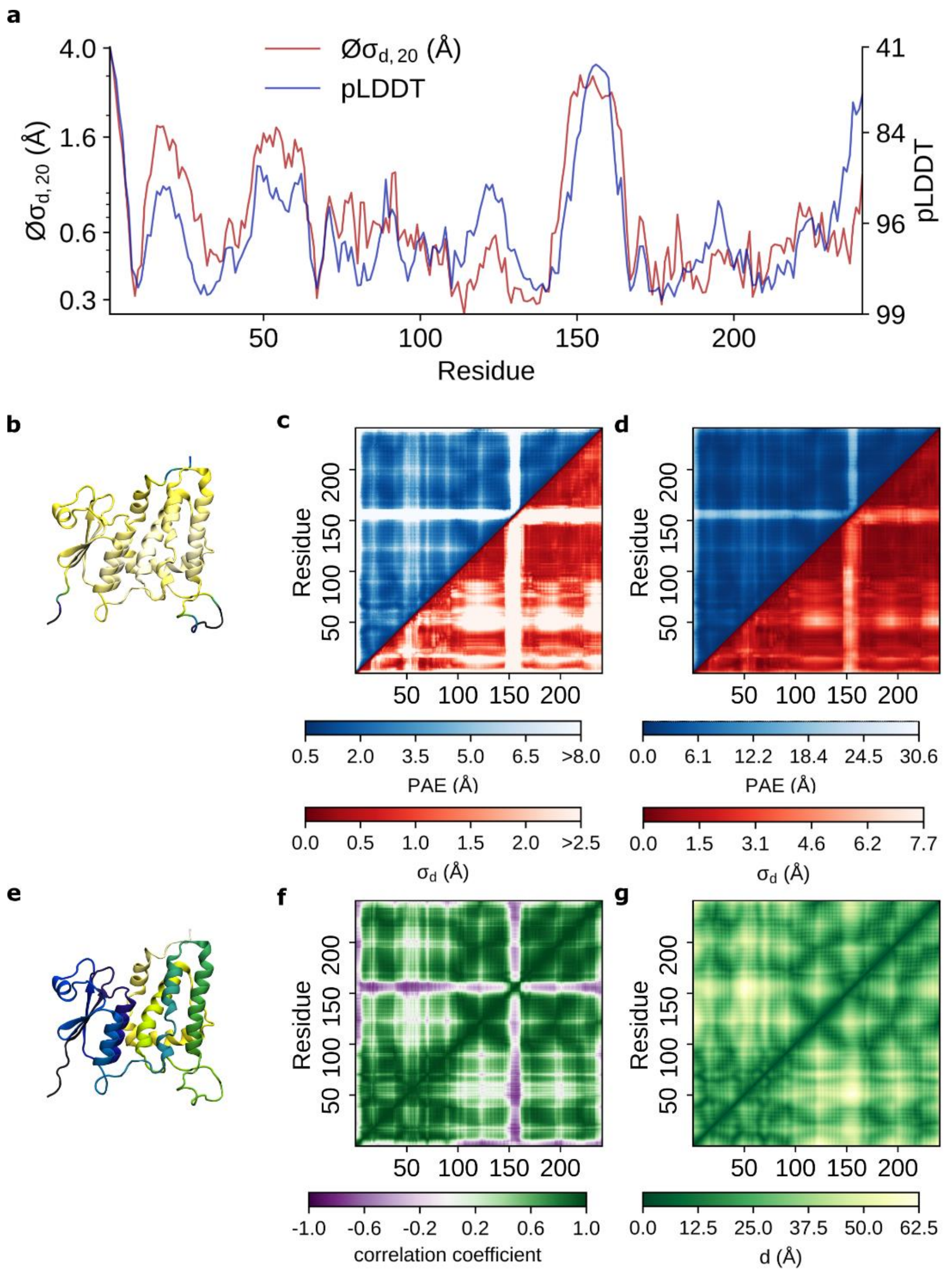


**Figure S11:** AlphaFold vs. aMD for *chloride intracellular channel protein 1* (protein 11) **a**) pLDDT scores vs. $Ø\sigma_{d,20}$ values **b**) The protein structure colored based on its pLDDT scores. The dark blue colors correspond to residues with a pLDDT score <= 60, while the white color corresponds to residues with a pLDDT score close to 100. **c**) Comparison between (symmetrized) PAE matrices (blue) against the standard deviation of all $C_\alpha$ distances $\sigma_d$ (red). The PAE scores range between 0.5 to 8.0 while the $\sigma_d$ are limited to <2.5 Å. **d**) Comparison between (symmetrized) PAE matrices (blue) against the standard deviation of all $C_\alpha$ distances $\sigma_d$ (red) for the maximum range. **e**) The protein structure colored based on its residue number. Dark blue colors correspond to low values, and yellows correspond to high values. **f**) Distance correlation matrix obtained from aMD simulations. **g**) Distance matrix obtained from aMD simulation.

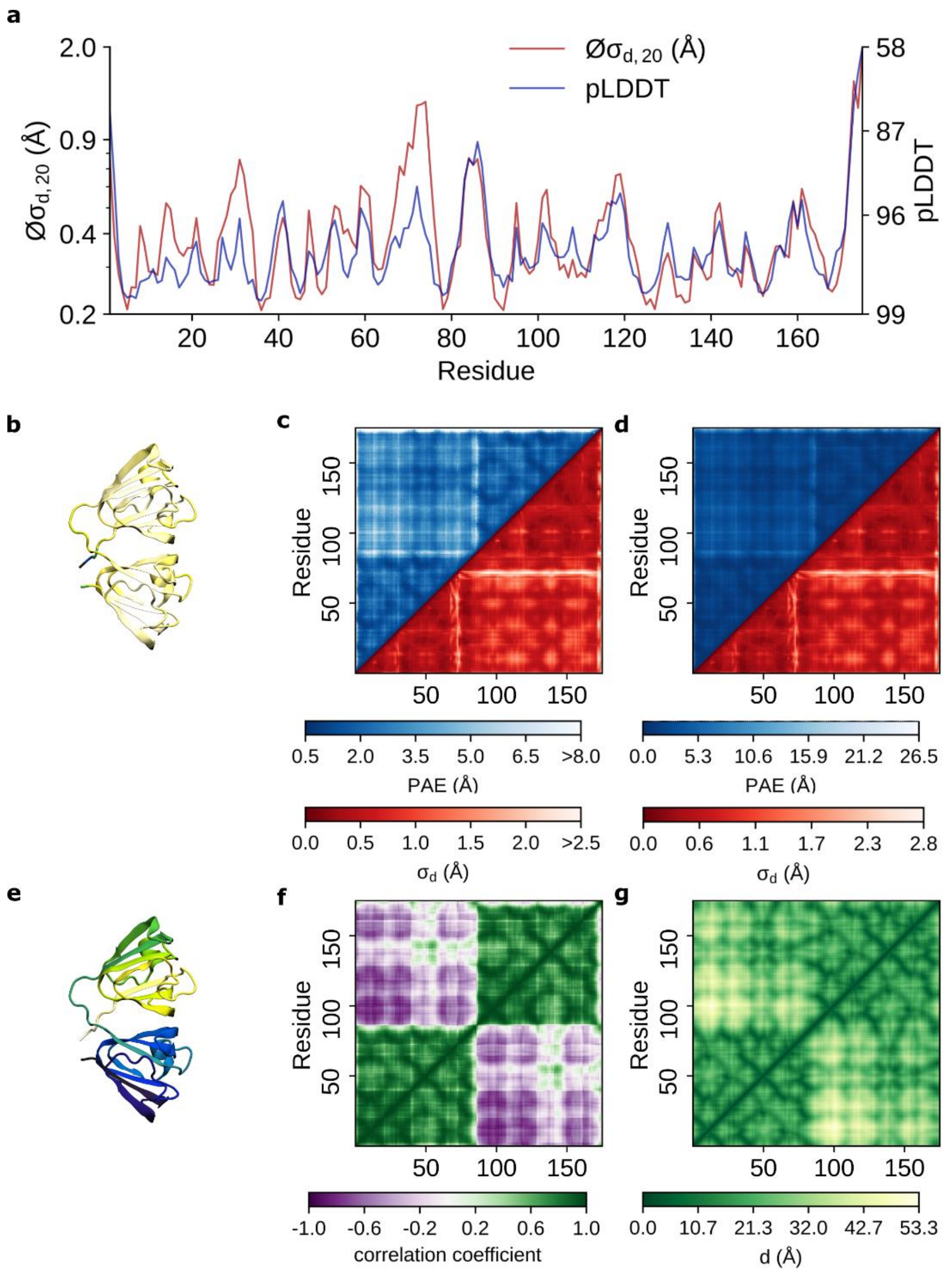


**Figure S12:** AlphaFold vs. aMD for *gamma-crystallin B* (protein 12) **a**) pLDDT scores vs. $\text{Ø}\sigma_{d,20}$ values **b**) The protein structure colored based on its pLDDT scores. The dark blue colors correspond to residues with a pLDDT score <= 60, while the white color corresponds to residues with a pLDDT score close to 100. **c**) Comparison between (symmetrized) PAE matrices (blue) against the standard deviation of all $C_\alpha$ distances $\sigma_d$ (red). The PAE scores range between 0.5 to 8.0 while the $\sigma_d$ are limited to <2.5 Å. **d**) Comparison between (symmetrized) PAE matrices (blue) against the standard deviation of all $C_\alpha$ distances $\sigma_d$ (red) for the maximum range. **e**) The protein structure colored based on its residue number. Dark blue colors correspond to low values, and yellows correspond to high values. **f**) Distance correlation matrix obtained from aMD simulations. **g**) Distance matrix obtained from aMD simulation.

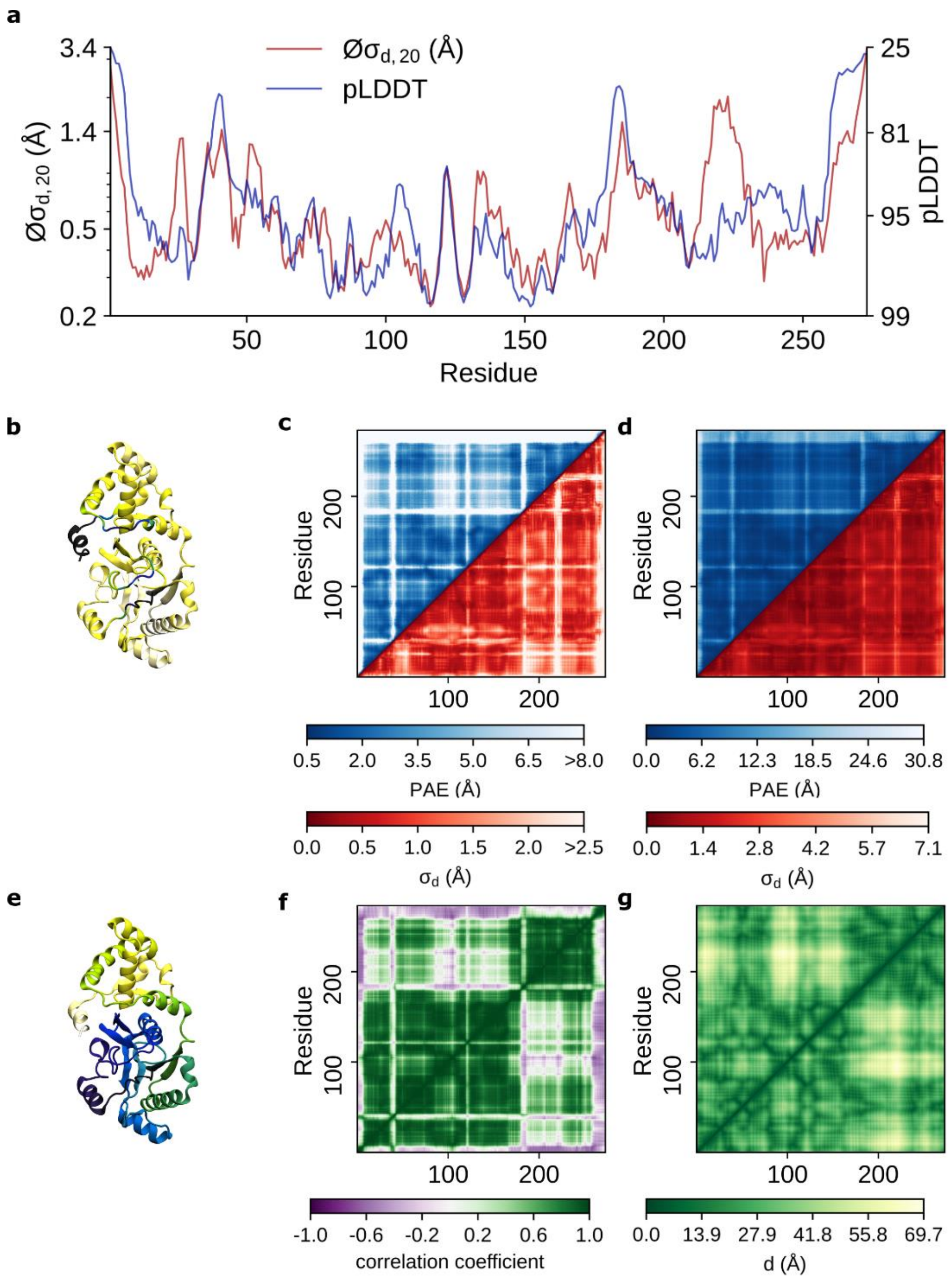


**Figure S13:** AlphaFold vs. aMD for *transposon Tn7 transposition protein TnsA* (protein 13) **a**) pLDDT scores vs. $Ø\sigma_{d,20}$ values **b**) The protein structure colored based on its pLDDT scores. The dark blue colors correspond to residues with a pLDDT score <= 60, while the white color corresponds to residues with a pLDDT score close to 100. **c**) Comparison between (symmetrized) PAE matrices (blue) against the standard deviation of all $C_\alpha$ distances $\sigma_d$ (red). The PAE scores range between 0.5 to 8.0 while the $\sigma_d$ are limited to <2.5 Å. **d**) Comparison between (symmetrized) PAE matrices (blue) against the standard deviation of all $C_\alpha$ distances $\sigma_d$ (red) for the maximum range. **e**) The protein structure colored based on its residue number. Dark blue colors correspond to low values, and yellows correspond to high values. **f**) Distance correlation matrix obtained from aMD simulations. **g**) Distance matrix obtained from aMD simulation.

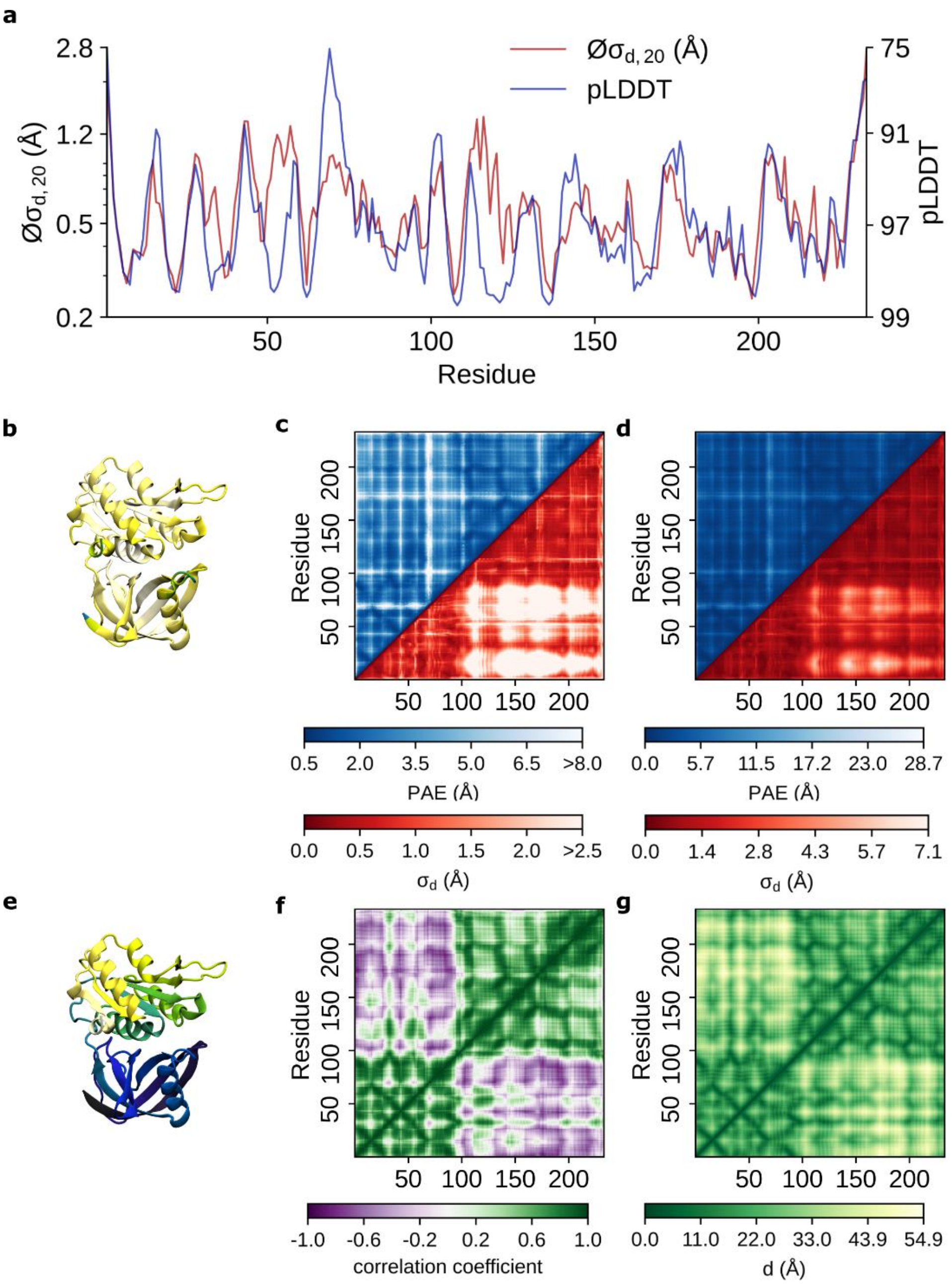


**Figure S14:** AlphaFold vs. aMD for *NAD(P)H-flavin reductase* (protein 14) **a**) pLDDT scores vs. $\text{Ø}\sigma_{d,20}$ values **b**) The protein structure colored based on its pLDDT scores. The dark blue colors correspond to residues with a pLDDT score <= 60, while the white color corresponds to residues with a pLDDT score close to 100. **c**) Comparison between (symmetrized) PAE matrices (blue) against the standard deviation of all $C_\alpha$ distances $\sigma_d$ (red). The PAE scores range between 0.5 to 8.0 while the $\sigma_d$ are limited to <2.5 Å. **d**) Comparison between (symmetrized) PAE matrices (blue) against the standard deviation of all $C_\alpha$ distances $\sigma_d$ (red) for the maximum range. **e**) The protein structure colored based on its residue number. Dark blue colors correspond to low values, and yellows correspond to high values. **f**) Distance correlation matrix obtained from aMD simulations. **g**) Distance matrix obtained from aMD simulation.

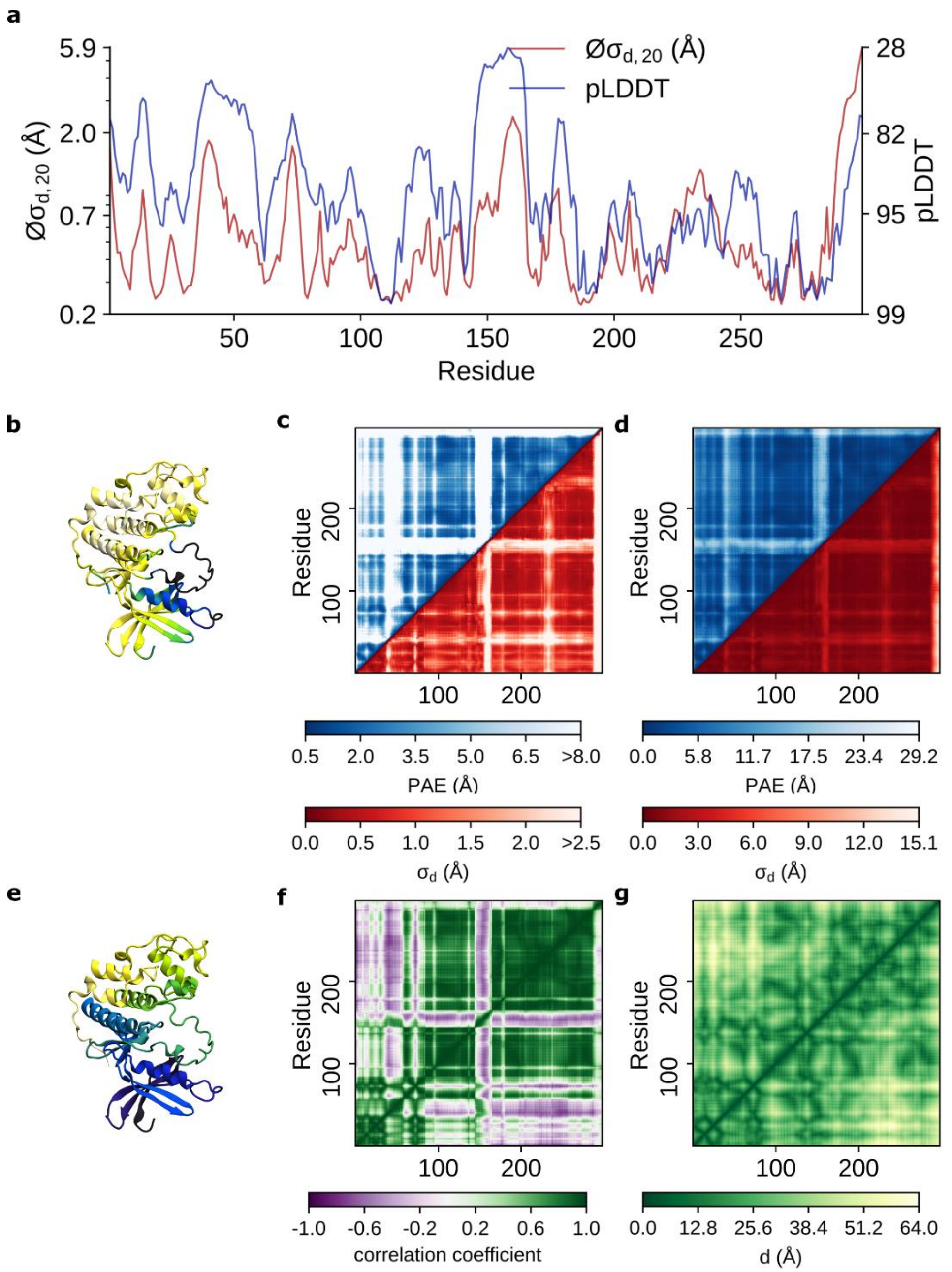


**Figure S15:** AlphaFold vs. aMD for *cyclin-dependent kinase 2* (protein 15) **a**) pLDDT scores vs. $Øσ_{d,20}$ values **b**) The protein structure colored based on its pLDDT scores. The dark blue colors correspond to residues with a pLDDT score <= 60, while the white color corresponds to residues with a pLDDT score close to 100. **c**) Comparison between (symmetrized) PAE matrices (blue) against the standard deviation of all $C_α$ distances $σ_d$ (red). The PAE scores range between 0.5 to 8.0 while the $σ_d$ are limited to <2.5 Å. **d**) Comparison between (symmetrized) PAE matrices (blue) against the standard deviation of all $C_α$ distances $σ_d$ (red) for the maximum range. **e**) The protein structure colored based on its residue number. Dark blue colors correspond to low values, and yellows correspond to high values. **f**) Distance correlation matrix obtained from aMD simulations. **g**) Distance matrix obtained from aMD simulation.

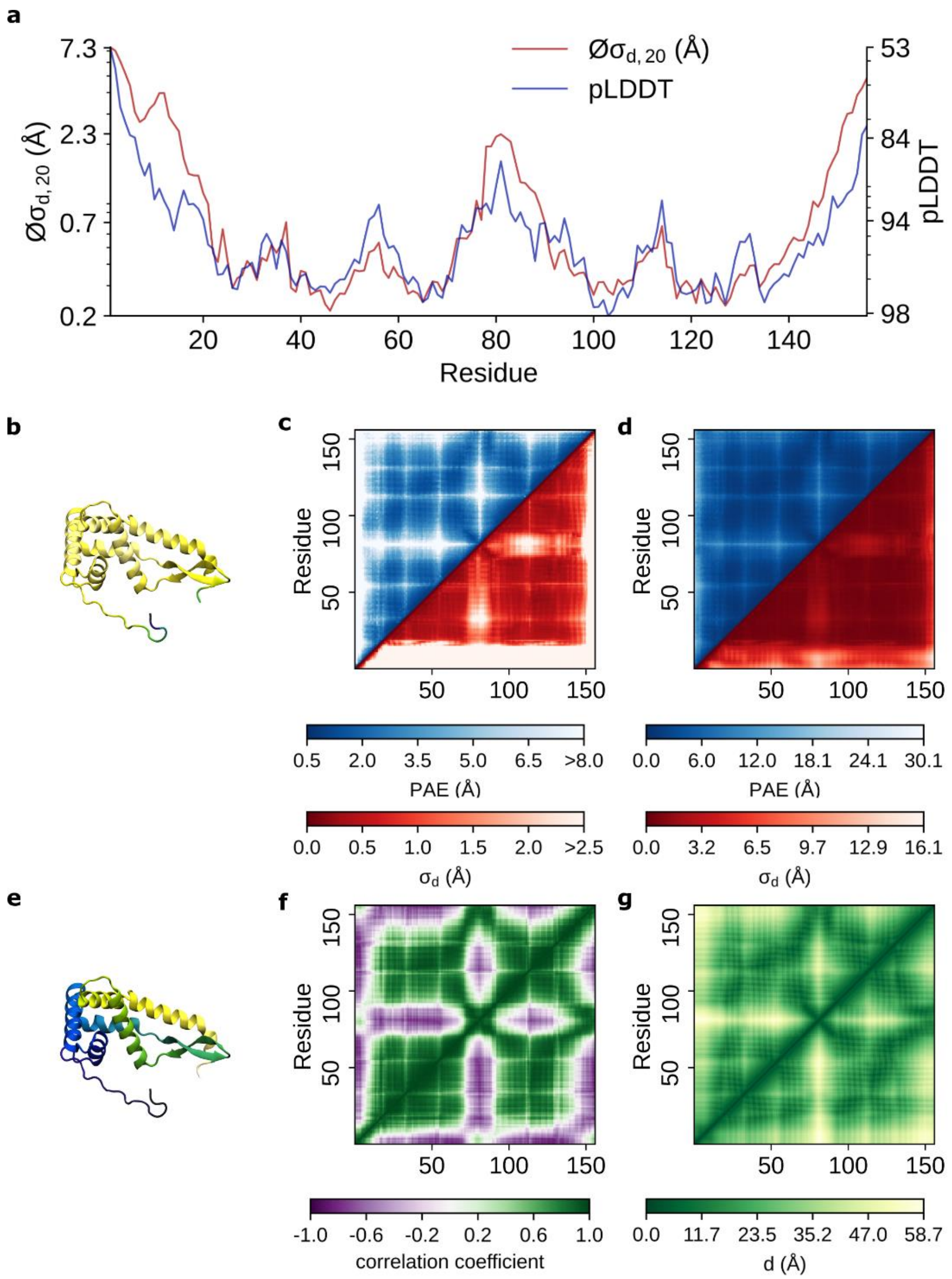


**Figure S16:** AlphaFold vs. aMD for *30S ribosomal protein S7* (protein 16) **a**) pLDDT scores vs. $Ø\sigma_{d,20}$ values **b**) The protein structure colored based on its pLDDT scores. The dark blue colors correspond to residues with a pLDDT score <= 60, while the white color corresponds to residues with a pLDDT score close to 100. **c**) Comparison between (symmetrized) PAE matrices (blue) against the standard deviation of all $C_\alpha$ distances $\sigma_d$ (red). The PAE scores range between 0.5 to 8.0 while the $\sigma_d$ are limited to <2.5 Å. **d**) Comparison between (symmetrized) PAE matrices (blue) against the standard deviation of all $C_\alpha$ distances $\sigma_d$ (red) for the maximum range. **e**) The protein structure colored based on its residue number. Dark blue colors correspond to low values, and yellows correspond to high values. **f**) Distance correlation matrix obtained from aMD simulations. **g**) Distance matrix obtained from aMD simulation.

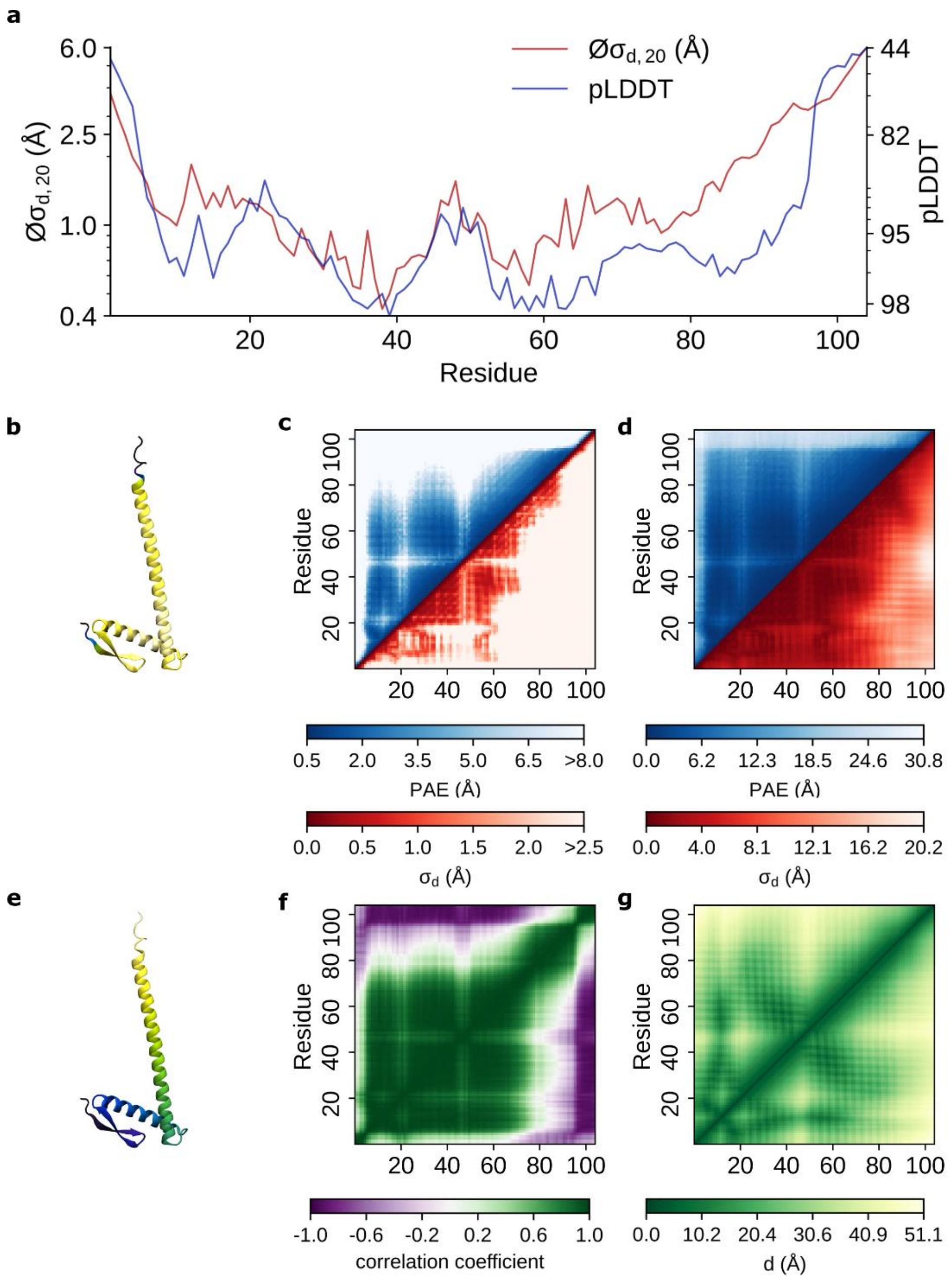


**Figure S17:** AlphaFold vs. aMD for *cell division protein ZapA* (protein 17) **a**) pLDDT scores vs. $Ø\sigma_{d,20}$ values **b**) The protein structure colored based on its pLDDT scores. The dark blue colors correspond to residues with a pLDDT score <= 60, while the white color corresponds to residues with a pLDDT score close to 100. **c**) Comparison between (symmetrized) PAE matrices (blue) against the standard deviation of all $C_\alpha$ distances $\sigma_d$ (red). The PAE scores range between 0.5 to 8.0 while the $\sigma_d$ are limited to <2.5 Å. **d**) Comparison between (symmetrized) PAE matrices (blue) against the standard deviation of all $C_\alpha$ distances $\sigma_d$ (red) for the maximum range. **e**) The protein structure colored based on its residue number. Dark blue colors correspond to low values, and yellows correspond to high values. **f**) Distance correlation matrix obtained from aMD simulations. **g**) Distance matrix obtained from aMD simulation.

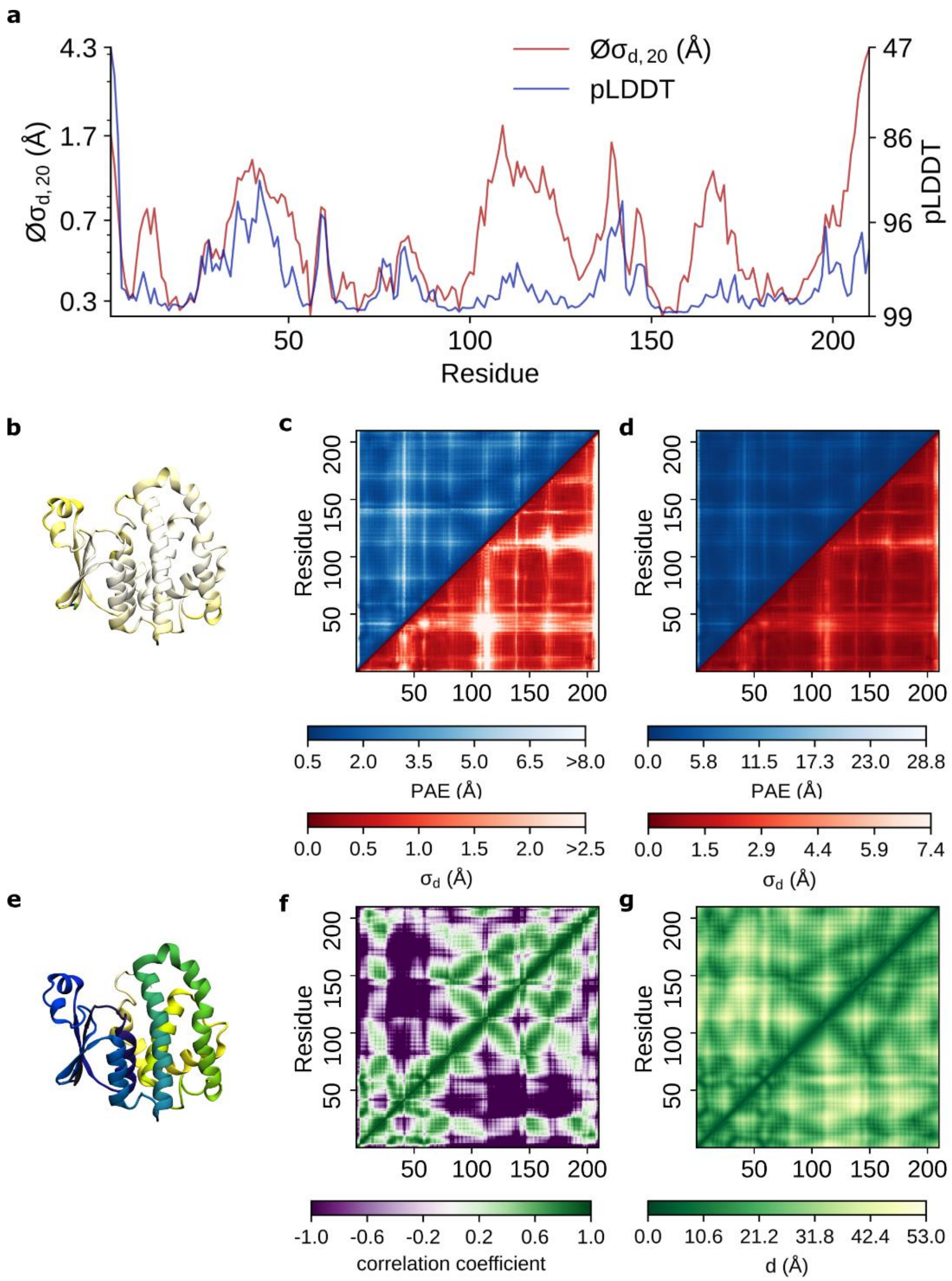


**Figure S18:** AlphaFold vs. aMD for *dephospho-CoA kinase* (protein 18) **a**) pLDDT scores vs. $Ø\sigma_{d,20}$ values **b**) The protein structure colored based on its pLDDT scores. The dark blue colors correspond to residues with a pLDDT score <= 60, while the white color corresponds to residues with a pLDDT score close to 100. **c**) Comparison between (symmetrized) PAE matrices (blue) against the standard deviation of all $C_\alpha$ distances $\sigma_d$ (red). The PAE scores range between 0.5 to 8.0 while the $\sigma_d$ are limited to <2.5 Å. **d**) Comparison between (symmetrized) PAE matrices (blue) against the standard deviation of all $C_\alpha$ distances $\sigma_d$ (red) for the maximum range. **e**) The protein structure colored based on its residue number. Dark blue colors correspond to low values, and yellows correspond to high values. **f**) Distance correlation matrix obtained from aMD simulations. **g**) Distance matrix obtained from aMD simulation.

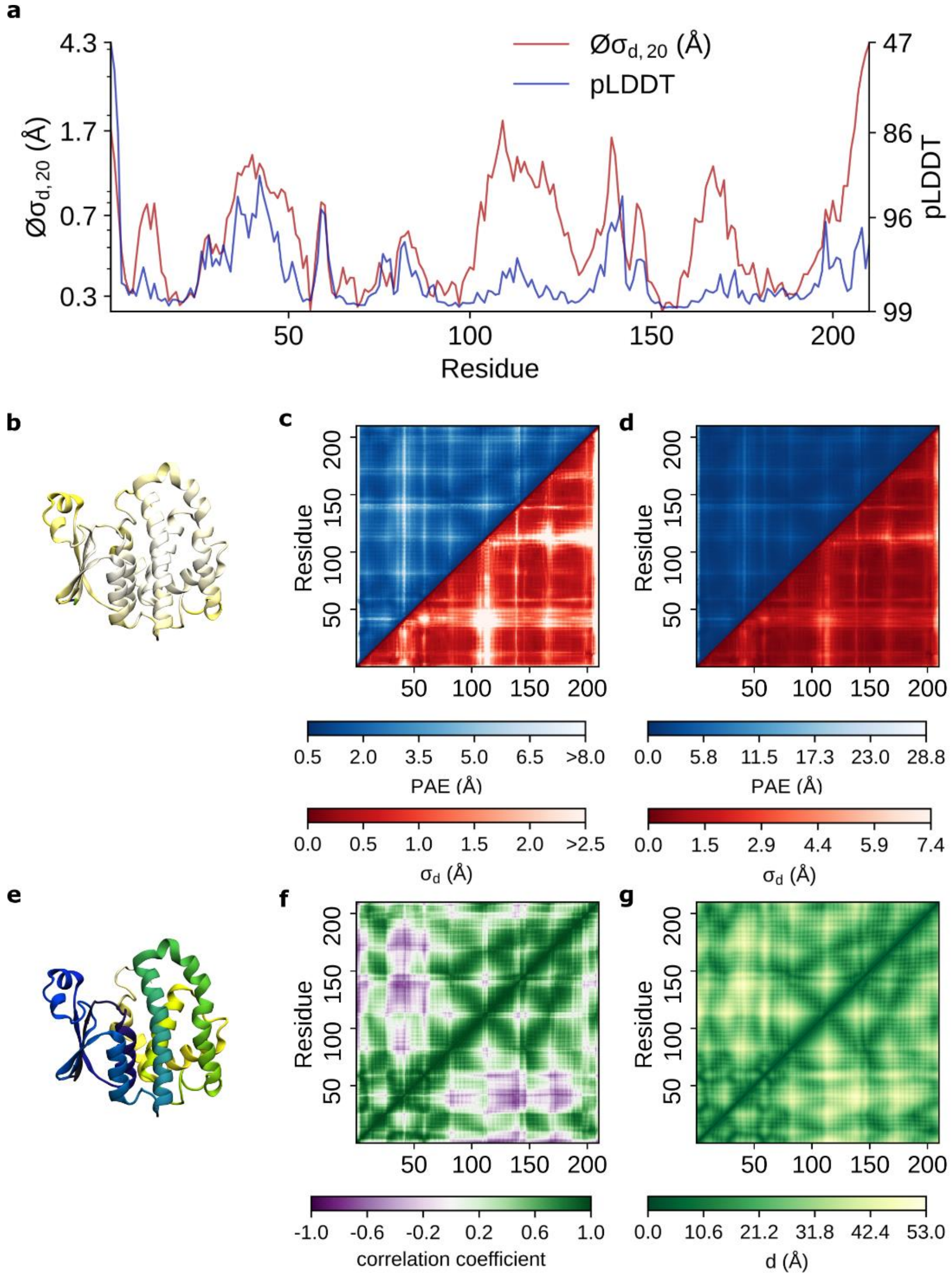


**Figure S19:** AlphaFold vs. aMD for *glutathione S-transferase P 1* (protein 19) **a**) pLDDT scores vs. $Ø\sigma_{d,20}$ values **b**) The protein structure colored based on its pLDDT scores. The dark blue colors correspond to residues with a pLDDT score <= 60, while the white color corresponds to residues with a pLDDT score close to 100. **c**) Comparison between (symmetrized) PAE matrices (blue) against the standard deviation of all $C_\alpha$ distances $\sigma_d$ (red). The PAE scores range between 0.5 to 8.0 while the $\sigma_d$ are limited to <2.5 Å. **d**) Comparison between (symmetrized) PAE matrices (blue) against the standard deviation of all $C_\alpha$ distances $\sigma_d$ (red) for the maximum range. **e**) The protein structure colored based on its residue number. Dark blue colors correspond to low values, and yellows correspond to high values. **f**) Distance correlation matrix obtained from aMD simulations. **g**) Distance matrix obtained from aMD simulation.

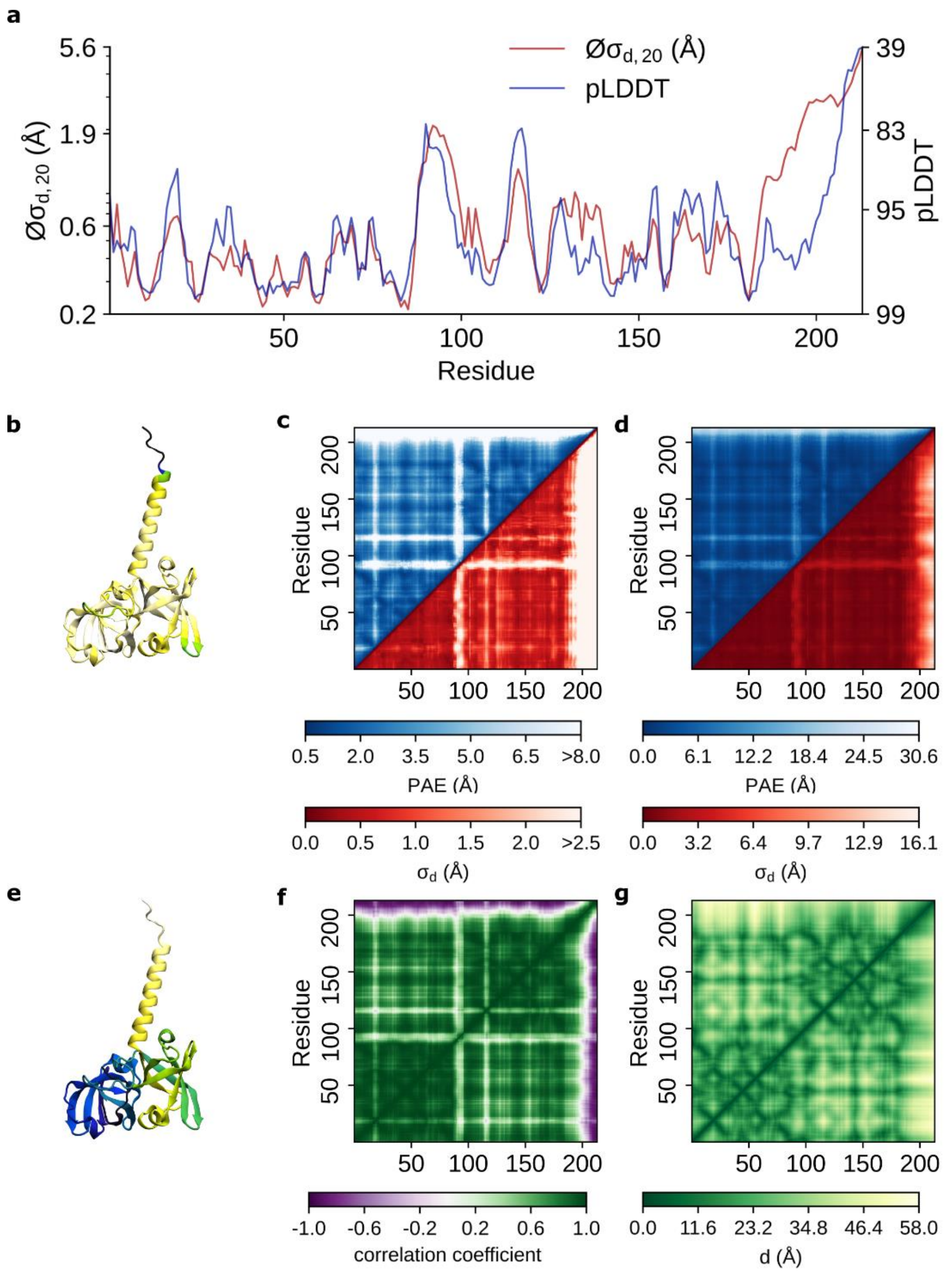


**Figure S20:** AlphaFold vs. aMD for *riboflavin synthase* (protein 20) **a**) pLDDT scores vs. $Ø\sigma_{d,20}$ values **b**) The protein structure colored based on its pLDDT scores. The dark blue colors correspond to residues with a pLDDT score <= 60, while the white color corresponds to residues with a pLDDT score close to 100. **c**) Comparison between (symmetrized) PAE matrices (blue) against the standard deviation of all $C_\alpha$ distances $\sigma_d$ (red). The PAE scores range between 0.5 to 8.0 while the $\sigma_d$ are limited to <2.5 Å. **d**) Comparison between (symmetrized) PAE matrices (blue) against the standard deviation of all $C_\alpha$ distances $\sigma_d$ (red) for the maximum range. **e**) The protein structure colored based on its residue number. Dark blue colors correspond to low values, and yellows correspond to high values. **f**) Distance correlation matrix obtained from aMD simulations. **g**) Distance matrix obtained from aMD simulation.

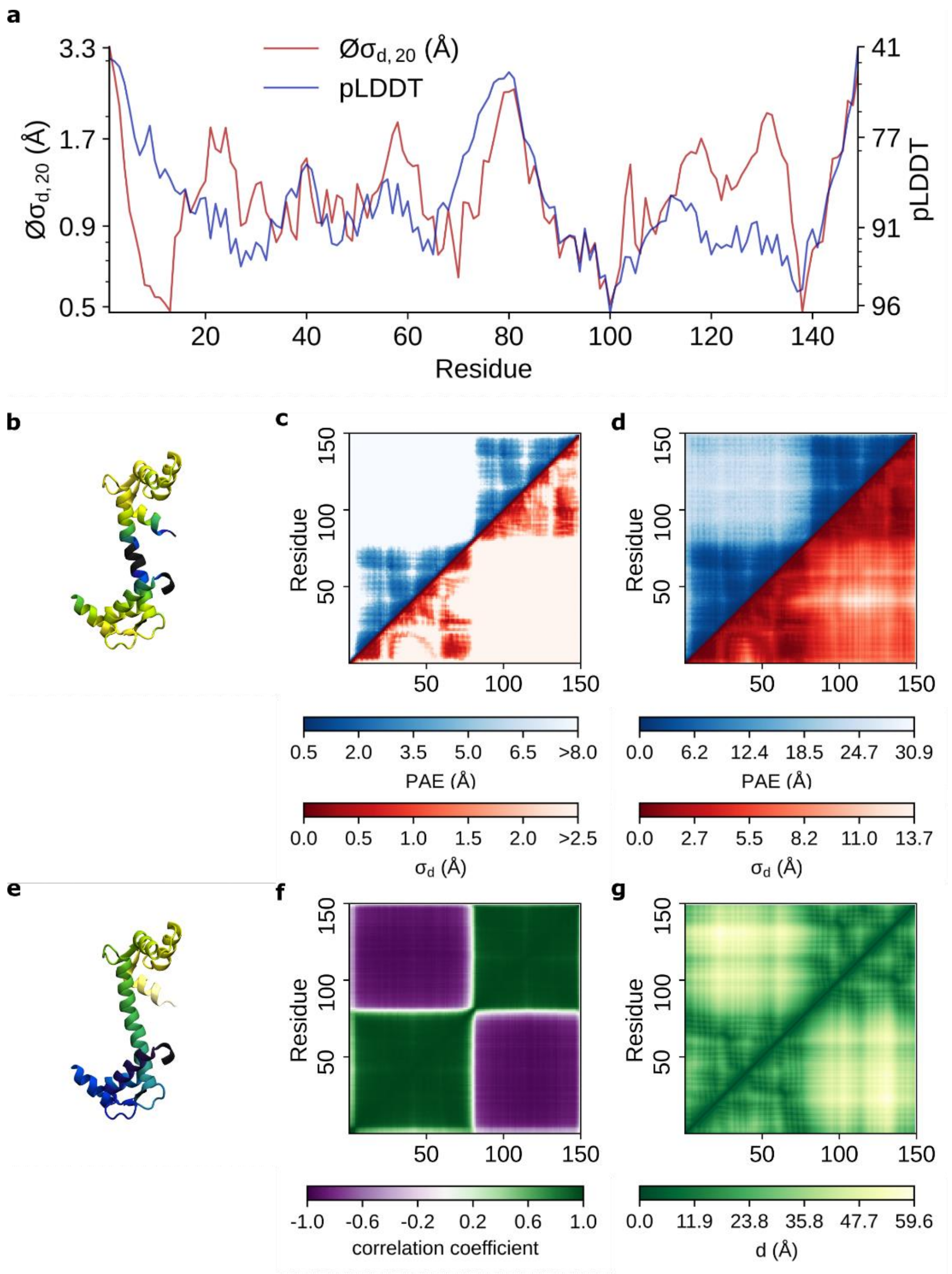


**Figure S21:** AlphaFold vs. aMD for *calmodulin* (protein 21) **a**) pLDDT scores vs. $Øσ_{d,20}$ values **b**) The protein structure colored based on its pLDDT scores. The dark blue colors correspond to residues with a pLDDT score <= 60, while the white color corresponds to residues with a pLDDT score close to 100. **c**) Comparison between (symmetrized) PAE matrices (blue) against the standard deviation of all $C_α$ distances $σ_d$ (red). The PAE scores range between 0.5 to 8.0 while the $σ_d$ are limited to <2.5 Å. **d**) Comparison between (symmetrized) PAE matrices (blue) against the standard deviation of all $C_α$ distances $σ_d$ (red) for the maximum range. **e**) The protein structure colored based on its residue number. Dark blue colors correspond to low values, and yellows correspond to high values. **f**) Distance correlation matrix obtained from aMD simulations. **g**) Distance matrix obtained from aMD simulation.

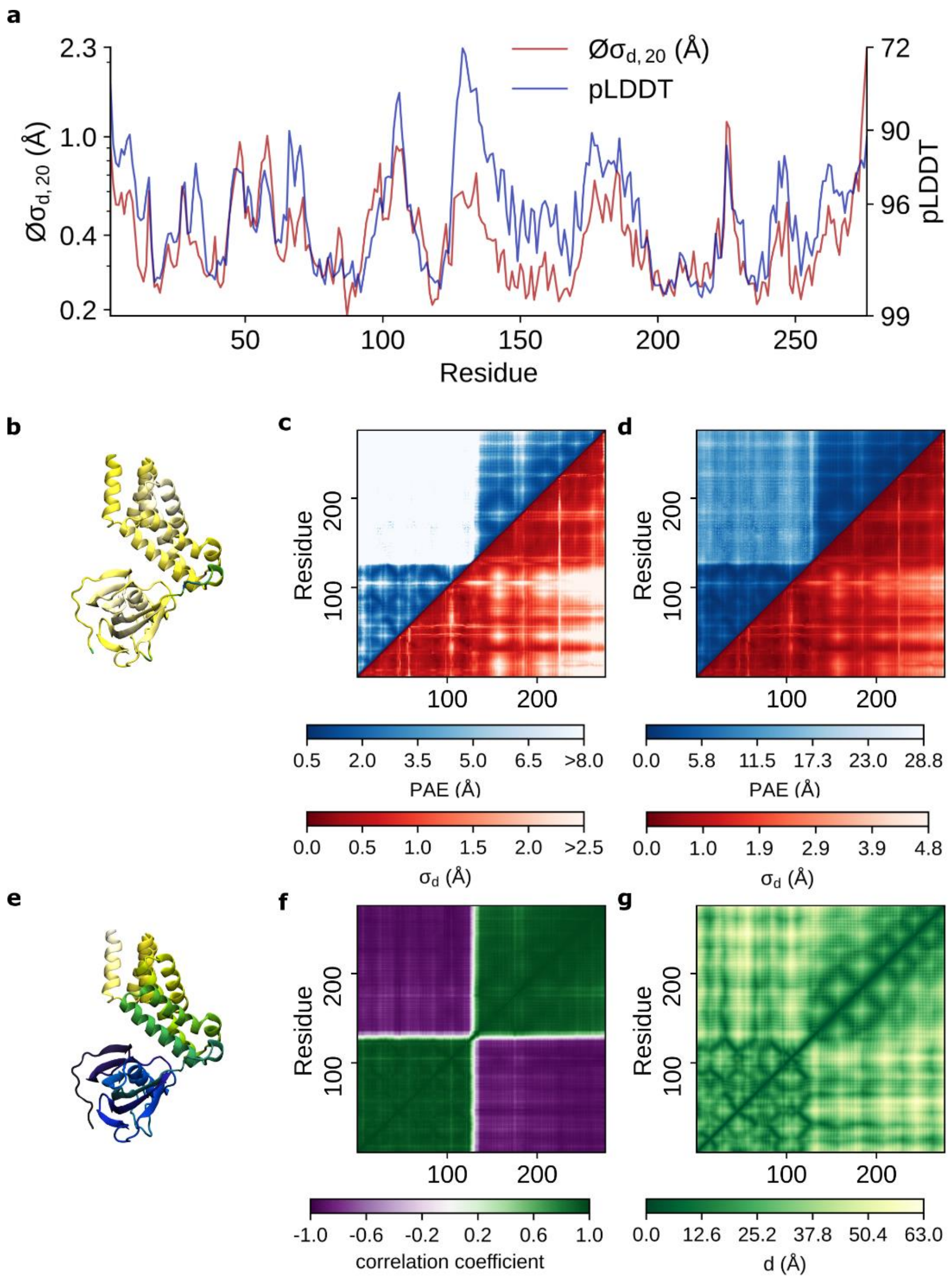


**Figure S22:** AlphaFold vs. aMD for *peptidyl-prolyl cis-trans isomerase FKBP42* (protein 22) **a**) pLDDT scores vs. $Ø\sigma_{d,20}$ values **b**) The protein structure colored based on its pLDDT scores. The dark blue colors correspond to residues with a pLDDT score <= 60, while the white color corresponds to residues with a pLDDT score close to 100. **c**) Comparison between (symmetrized) PAE matrices (blue) against the standard deviation of all $C_\alpha$ distances $\sigma_d$ (red). The PAE scores range between 0.5 to 8.0 while the $\sigma_d$ are limited to <2.5 Å. **d**) Comparison between (symmetrized) PAE matrices (blue) against the standard deviation of all $C_\alpha$ distances $\sigma_d$ (red) for the maximum range. **e**) The protein structure colored based on its residue number. Dark blue colors correspond to low values, and yellows correspond to high values. **f**) Distance correlation matrix obtained from aMD simulations. **g**) Distance matrix obtained from aMD simulation.

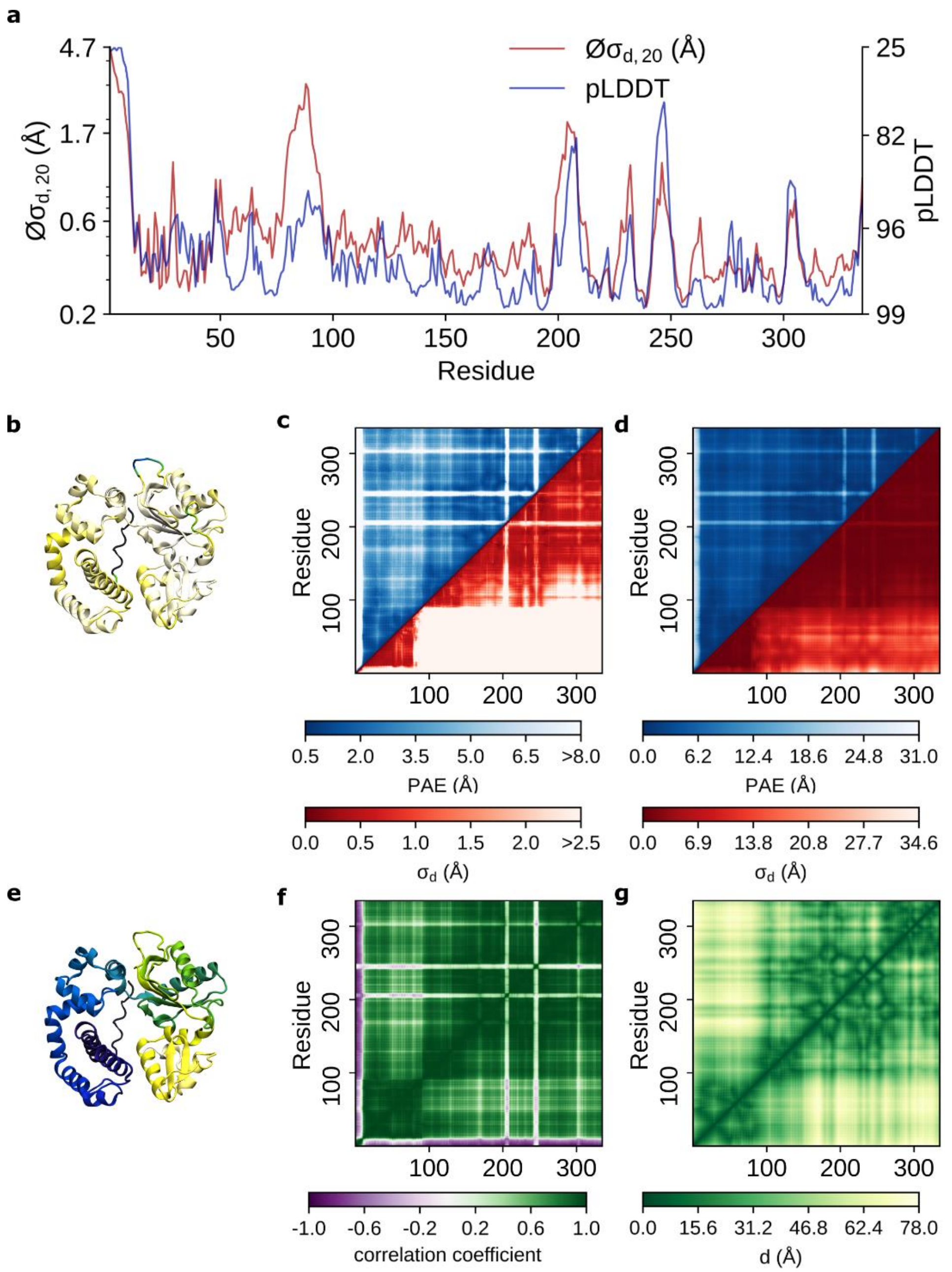


**Figure S23:** AlphaFold vs. aMD for *DNA polymerase beta* (protein 23) **a**) pLDDT scores vs. $\varnothing\sigma_{d,20}$ values **b**) The protein structure colored based on its pLDDT scores. The dark blue colors correspond to residues with a pLDDT score <= 60, while the white color corresponds to residues with a pLDDT score close to 100. **c**) Comparison between (symmetrized) PAE matrices (blue) against the standard deviation of all $C_\alpha$ distances $\sigma_d$ (red). The PAE scores range between 0.5 to 8.0 while the $\sigma_d$ are limited to <2.5 Å. **d**) Comparison between (symmetrized) PAE matrices (blue) against the standard deviation of all $C_\alpha$ distances $\sigma_d$ (red) for the maximum range. **e**) The protein structure colored based on its residue number. Dark blue colors correspond to low values, and yellows correspond to high values. **f**) Distance correlation matrix obtained from aMD simulations. **g**) Distance matrix obtained from aMD simulation.

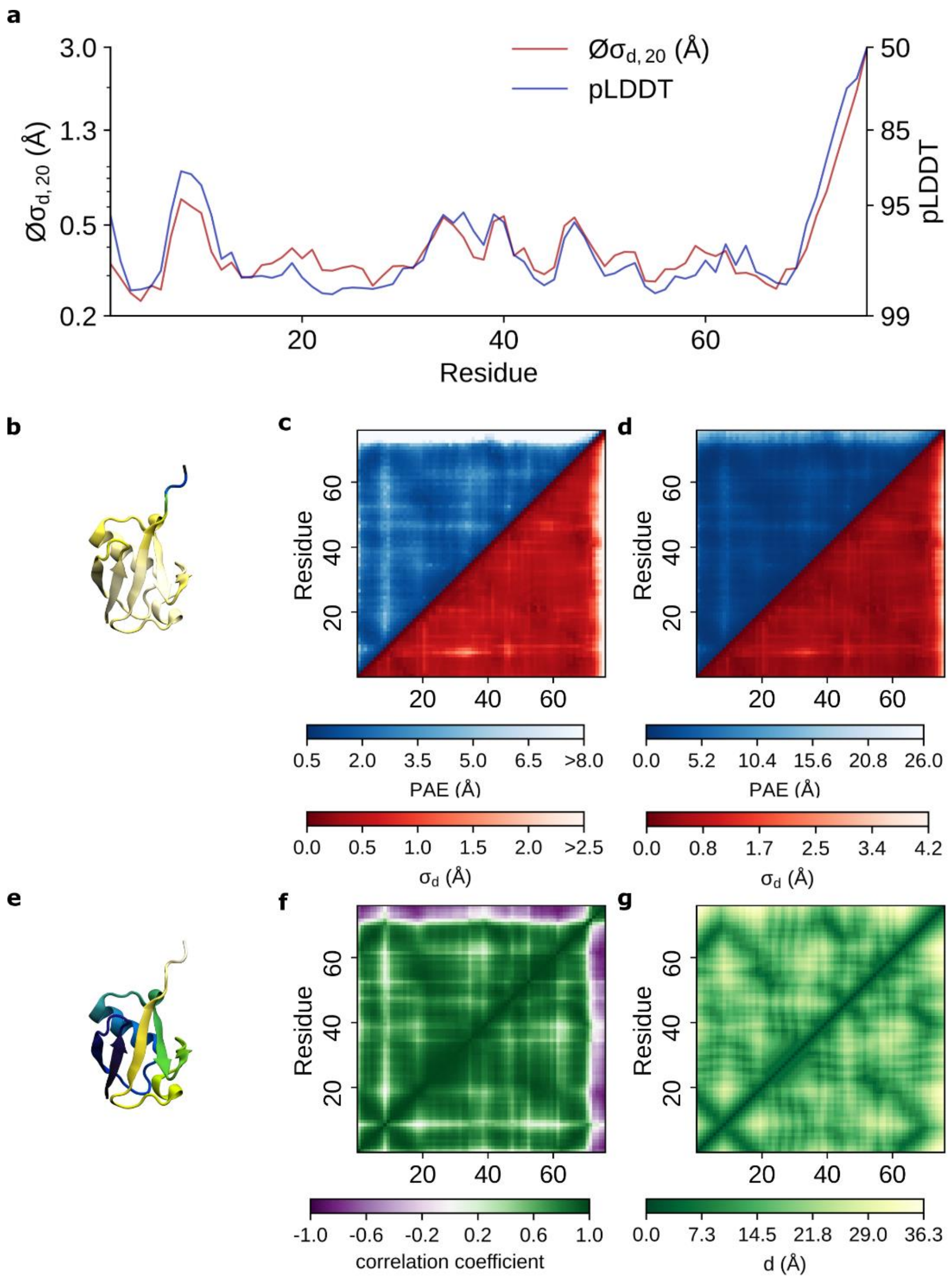


**Figure S24:** AlphaFold vs. aMD for *ubiquitin* (protein 24) **a**) pLDDT scores vs. $Ø\sigma_{d,20}$ values **b**) The protein structure colored based on its pLDDT scores. The dark blue colors correspond to residues with a pLDDT score <= 60, while the white color corresponds to residues with a pLDDT score close to 100. **c**) Comparison between (symmetrized) PAE matrices (blue) against the standard deviation of all $C_\alpha$ distances $\sigma_d$ (red). The PAE scores range between 0.5 to 8.0 while the $\sigma_d$ are limited to <2.5 Å. **d**) Comparison between (symmetrized) PAE matrices (blue) against the standard deviation of all $C_\alpha$ distances $\sigma_d$ (red) for the maximum range. **e**) The protein structure colored based on its residue number. Dark blue colors correspond to low values, and yellows correspond to high values. **f**) Distance correlation matrix obtained from aMD simulations. **g**) Distance matrix obtained from aMD simulation.

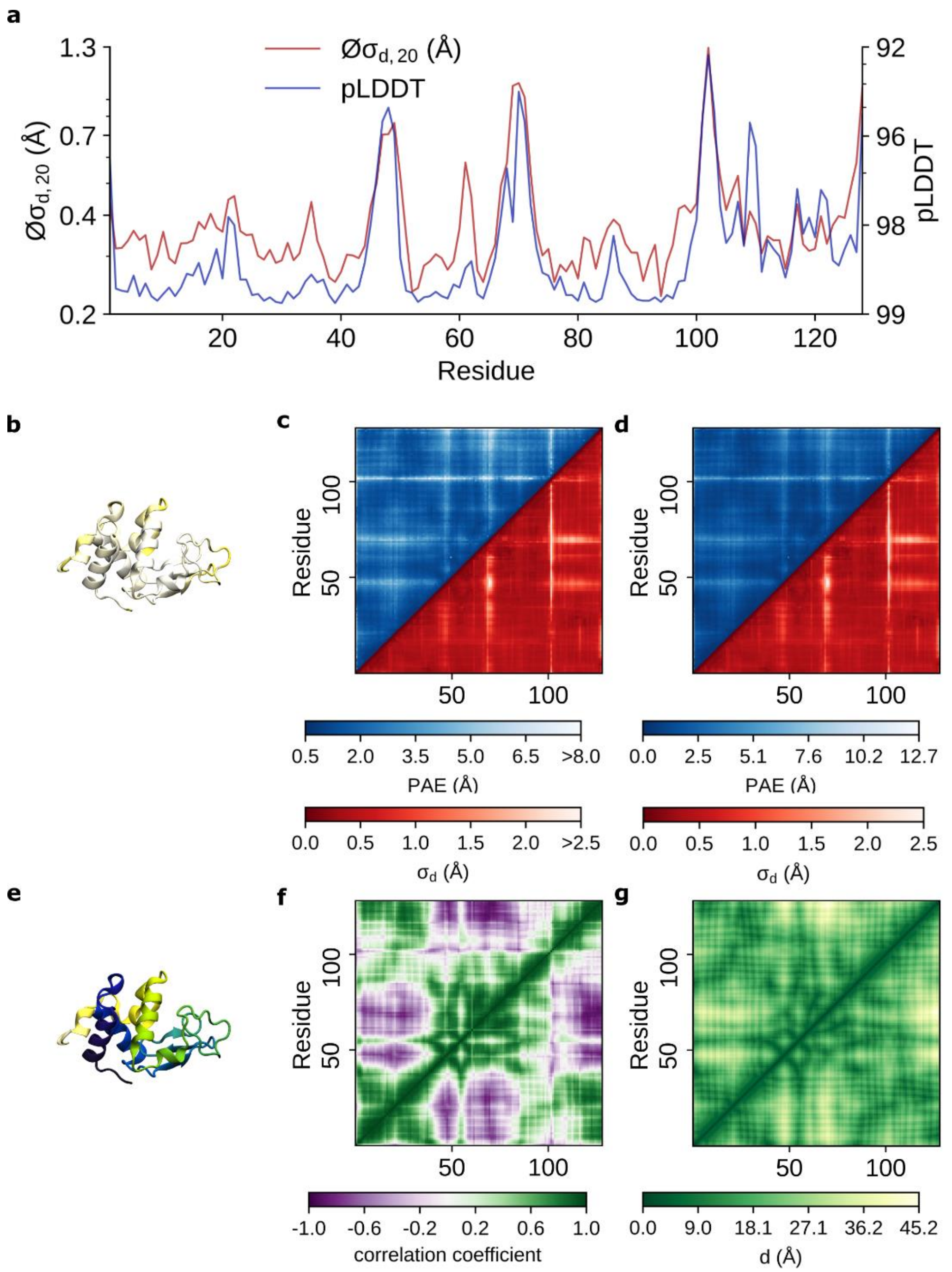


**Figure S25:** AlphaFold vs. aMD for *lysozyme C* (protein 25) **a**) pLDDT scores vs. $Øσ_{d,20}$ values **b**) The protein structure colored based on its pLDDT scores. The dark blue colors correspond to residues with a pLDDT score <= 60, while the white color corresponds to residues with a pLDDT score close to 100. **c**) Comparison between (symmetrized) PAE matrices (blue) against the standard deviation of all $C_α$ distances $σ_d$ (red). The PAE scores range between 0.5 to 8.0 while the $σ_d$ are limited to <2.5 Å. **d**) Comparison between (symmetrized) PAE matrices (blue) against the standard deviation of all $C_α$ distances $σ_d$ (red) for the maximum range. **e**) The protein structure colored based on its residue number. Dark blue colors correspond to low values, and yellows correspond to high values. **f**) Distance correlation matrix obtained from aMD simulations. **g**) Distance matrix obtained from aMD simulation.

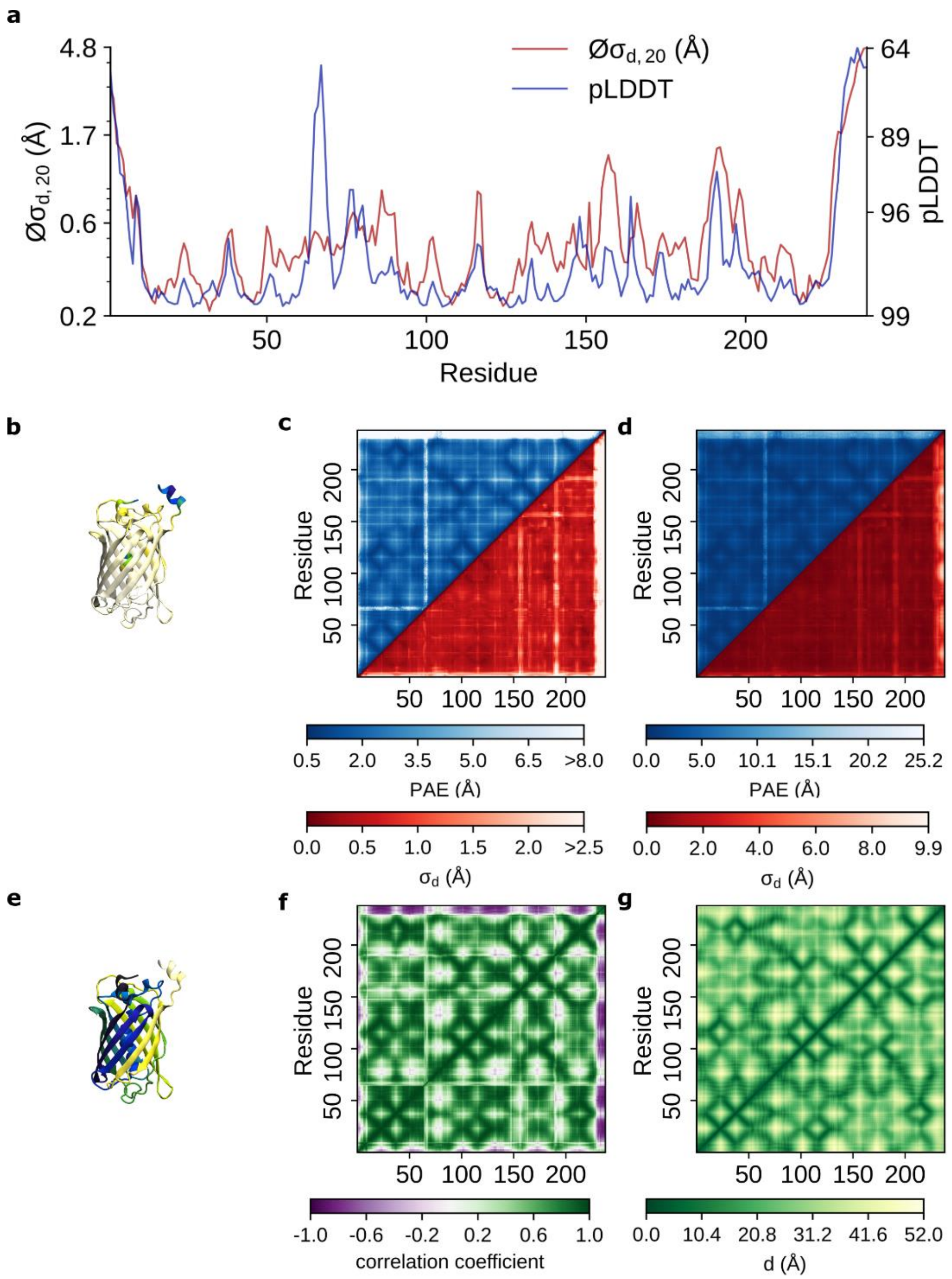


**Figure S26:** AlphaFold vs. aMD for *green fluorescent protein* (protein 26) **a**) pLDDT scores vs. $Øσ_{d,20}$ values **b**) The protein structure colored based on its pLDDT scores. The dark blue colors correspond to residues with a pLDDT score <= 60, while the white color corresponds to residues with a pLDDT score close to 100. **c**) Comparison between (symmetrized) PAE matrices (blue) against the standard deviation of all $C_α$ distances $σ_d$ (red). The PAE scores range between 0.5 to 8.0 while the $σ_d$ are limited to <2.5 Å. **d**) Comparison between (symmetrized) PAE matrices (blue) against the standard deviation of all $C_α$ distances $σ_d$ (red) for the maximum range. **e**) The protein structure colored based on its residue number. Dark blue colors correspond to low values, and yellows correspond to high values. **f**) Distance correlation matrix obtained from aMD simulations. **g**) Distance matrix obtained from aMD simulation.

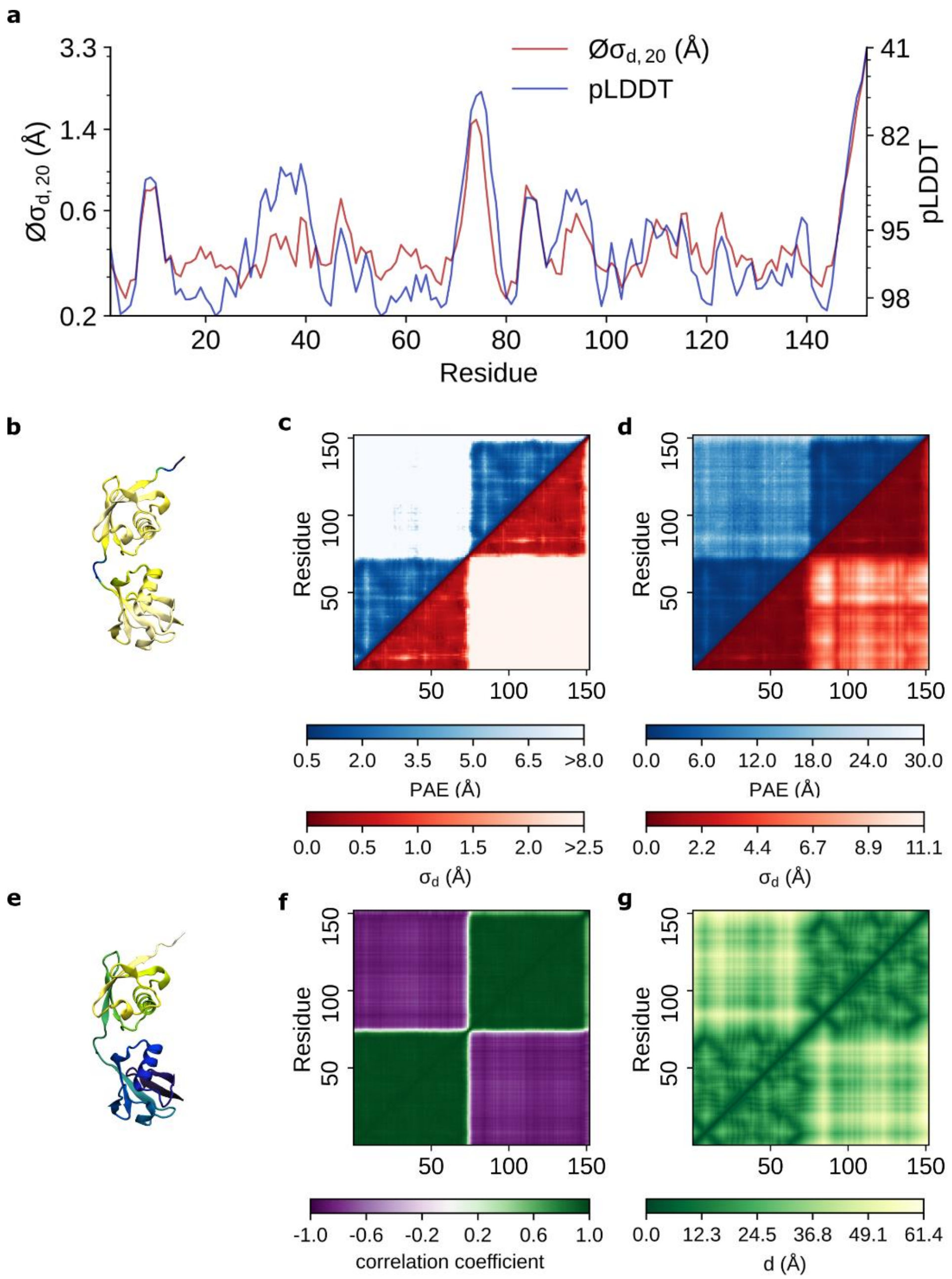


**Figure S27:** AlphaFold vs. aMD for *linear diubiquitin* (protein 27) **a**) pLDDT scores vs. $Øσ_{d,20}$ values **b**) The protein structure colored based on its pLDDT scores. The dark blue colors correspond to residues with a pLDDT score <= 60, while the white color corresponds to residues with a pLDDT score close to 100. **c**) Comparison between (symmetrized) PAE matrices (blue) against the standard deviation of all $C_α$ distances $σ_d$ (red). The PAE scores range between 0.5 to 8.0 while the $σ_d$ are limited to <2.5 Å. **d**) Comparison between (symmetrized) PAE matrices (blue) against the standard deviation of all $C_α$ distances $σ_d$ (red) for the maximum range. **e**) The protein structure colored based on its residue number. Dark blue colors correspond to low values, and yellows correspond to high values. **f**) Distance correlation matrix obtained from aMD simulations. **g**) Distance matrix obtained from aMD simulation.

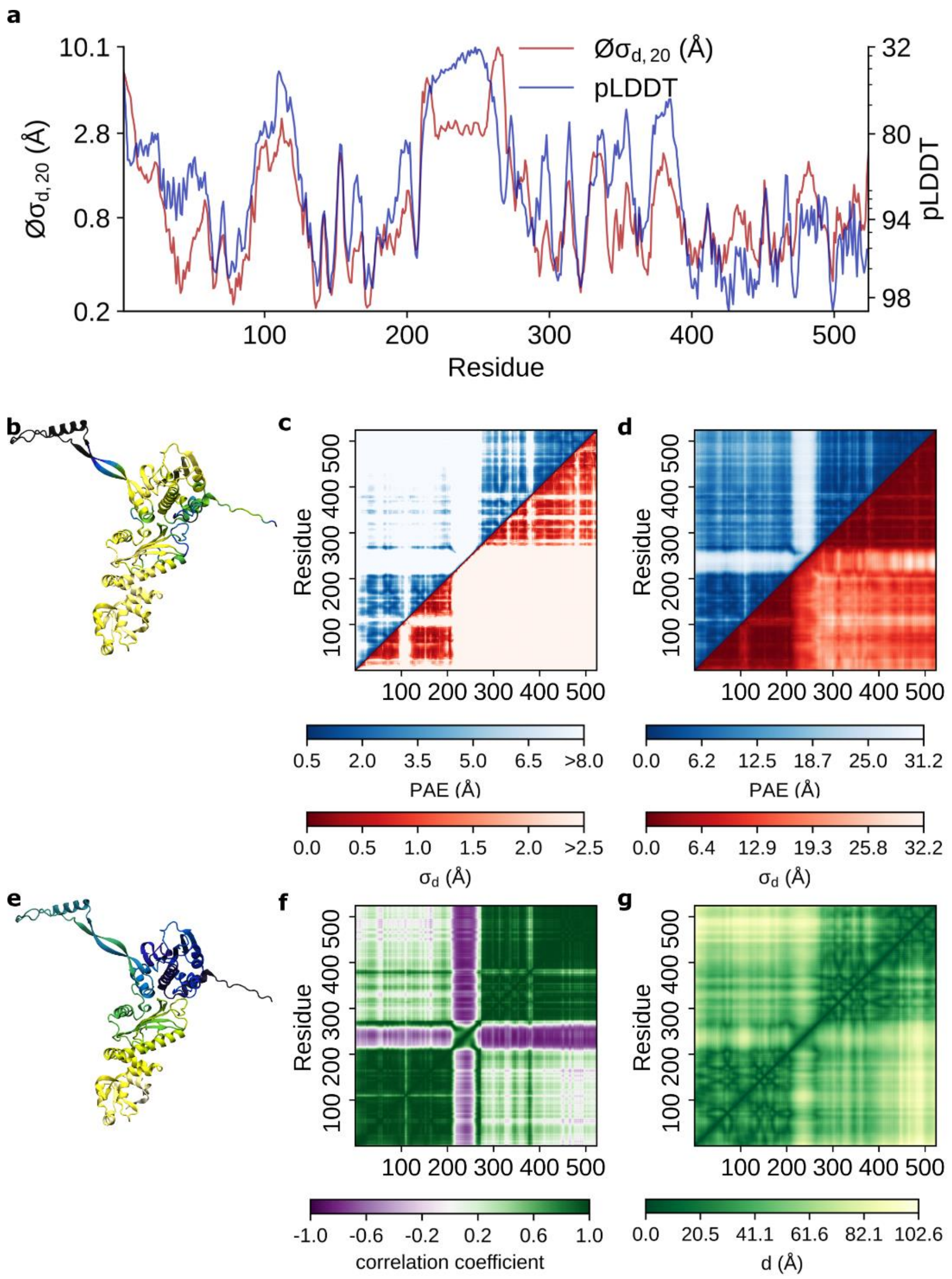


**Figure S28:** AlphaFold vs. aMD for *ATP-dependent molecular chaperon HSP82* (NTD-MD only) (protein 28) **a**) pLDDT scores vs. $Ø\sigma_{d,20}$ values **b**) The protein structure colored based on its pLDDT scores. The dark blue colors correspond to residues with a pLDDT score <= 60, while the white color corresponds to residues with a pLDDT score close to 100. **c**) Comparison between (symmetrized) PAE matrices (blue) against the standard deviation of all $C_\alpha$ distances $\sigma_d$ (red). The PAE scores range between 0.5 to 8.0 while the $\sigma_d$ are limited to <2.5 Å. **d**) Comparison between (symmetrized) PAE matrices (blue) against the standard deviation of all $C_\alpha$ distances $\sigma_d$ (red) for the maximum range. **e**) The protein structure colored based on its residue number. Dark blue colors correspond to low values, and yellows correspond to high values. **f**) Distance correlation matrix obtained from aMD simulations. **g**) Distance matrix obtained from aMD simulation.

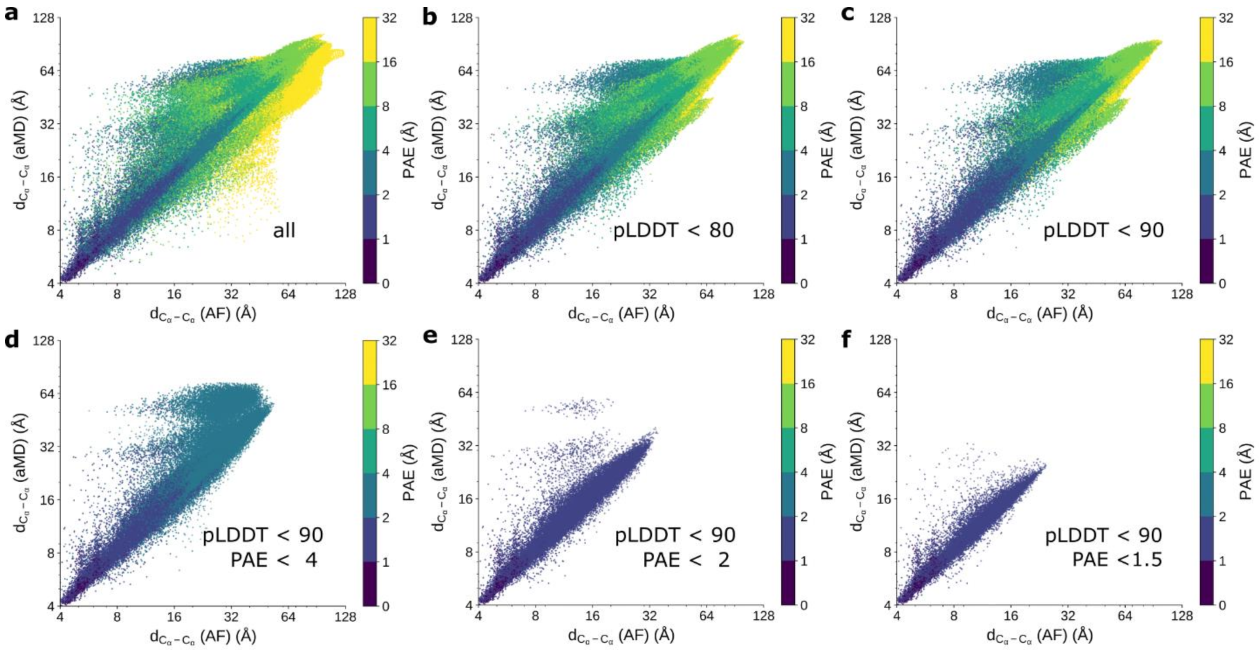


**Figure S29:** Protein distances predicted by AlphaFold (AF) compared to aMD under different constraints

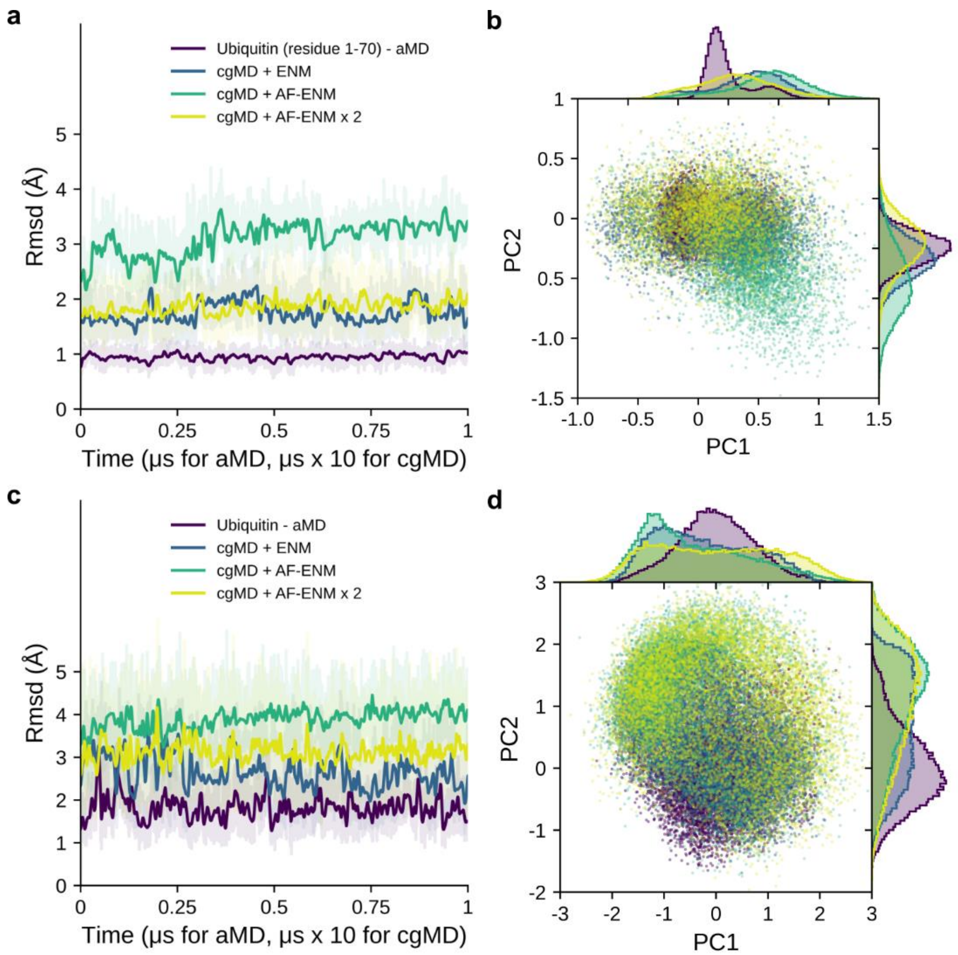


**Figure S30:** Ubiquitin (protein 24) global dynamics. **a,c**) RSMD of the aMD and cgMD simulations for (a) residue 1 - 70 or (c) complete system. **b,d**) Principal component analysis for aMD and cgMD simulations for (b) residue 1 - 70 or (d) complete system. The cgMD trajectories are projected on the aMD-derived first (PC1) and second (PC2) principal components.

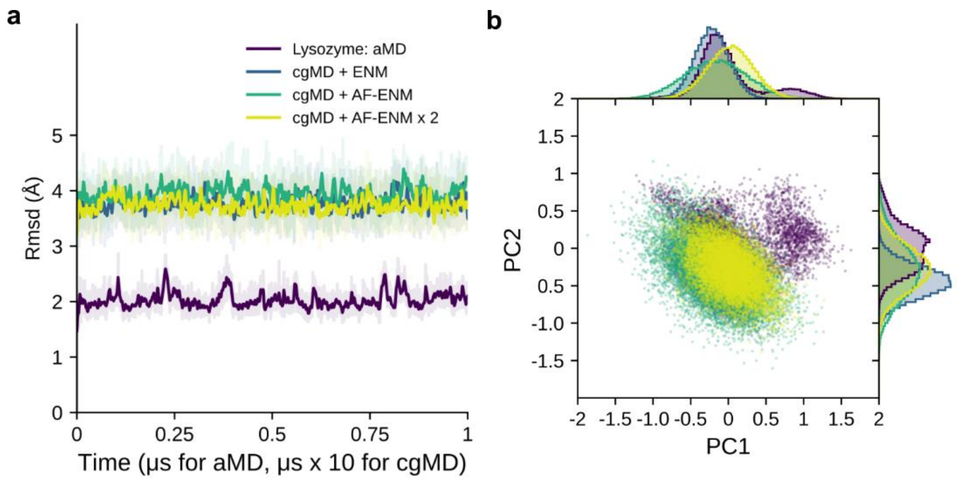


**Figure S31:** Lysozyme C (protein 25) global dynamics. **a**) RSMD of the aMD and cgMD simulations. **b**) Principal component analysis of aMD and cgMD simulations. The cgMD trajectories are projected on the aMD-derived first (PC1) and second (PC2) principal components.

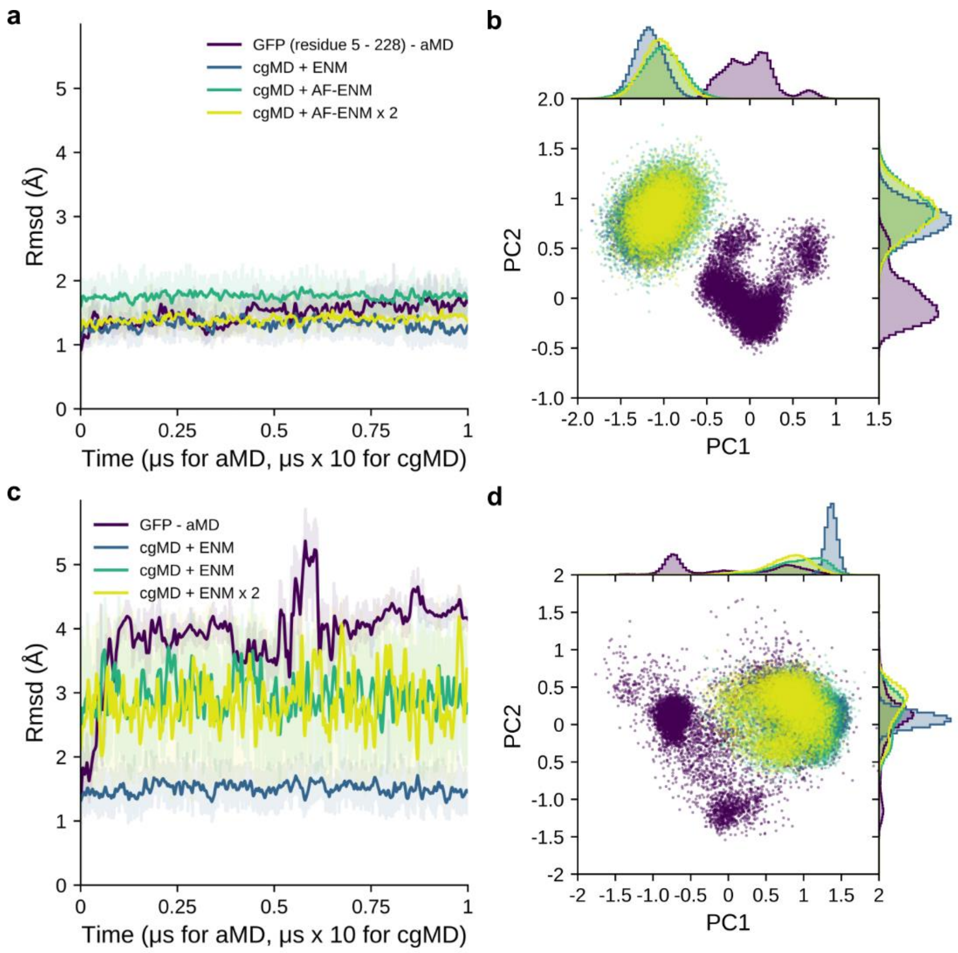


**Figure S32:** Green fluorescent protein (protein 26) global dynamics. **a,c**) RSMD of the aMD and cgMD simulations for (a) residue 5 - 228 or (c) complete system. **b,d**) Principal component analysis for aMD and cgMD simulations for (b) residue5 - 228 or (d) complete system. The cgMD trajectories are projected on the aMD-derived first (PC1) and second (PC2) principal components.

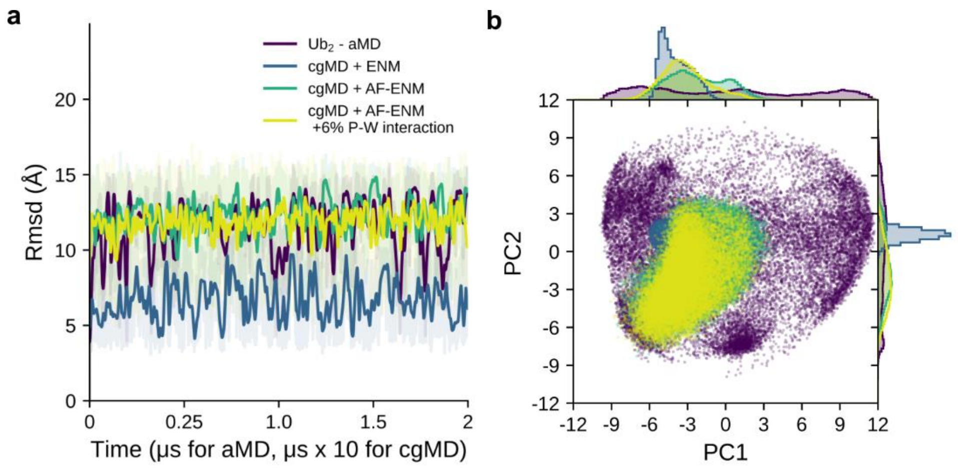


**Figure S33:** Linear Diubiquitin (protein 27) global dynamics. **a**) RSMD of the aMD and cgMD simulations. **b**) Principal component analysis of aMD and cgMD simulations. The cgMD trajectories are projected on the aMD-derived first (PC1) and second (PC2) principal components. The cgMD trajectories are projected on the aMD-derived first (PC1) and second (PC2) principal components.

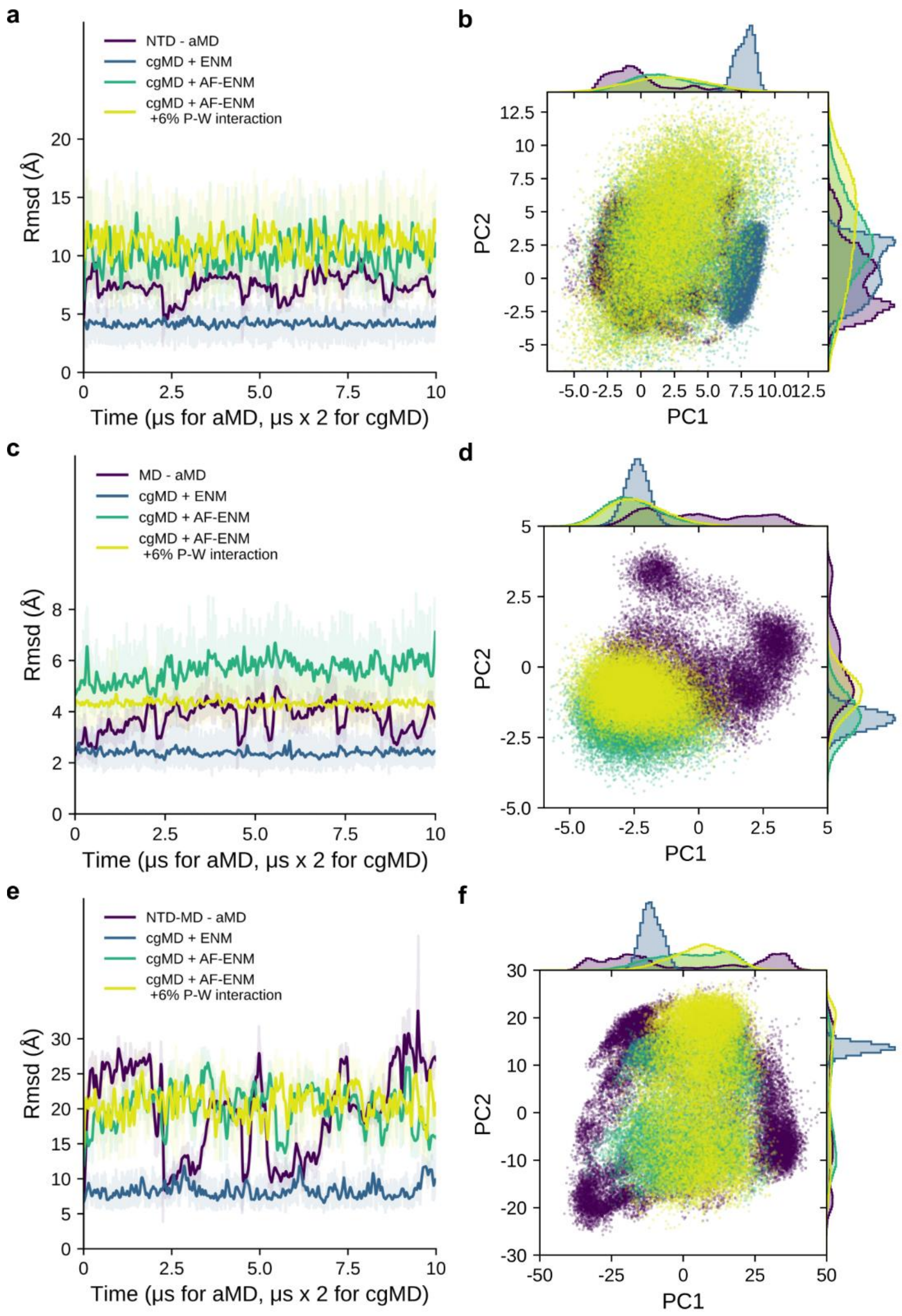


**Figure S34:** NTD-MD domain of Hsp90 (protein 28) global dynamics. **a,c,e**) RSMD of the aMD and cgMD simulations for (a) the N-terminal domain (NTD, residue 1 – 210), (c) middle domain (MD, residue 275 – 524) or (e) complete NTD-MD system (residue 1 - 524). . **b,d,f**) Principal component analysis for aMD and cgMD simulations for (b) NTD, (d) MD or (f) NTD-MD. The cgMD trajectories are projected on the aMD-derived first (PC1) and second (PC2) principal components.

**Table S1**: Summary of all simulated proteins.

| protein | Name | UniProt | Amino acids | Amino acid range | Simulation length [µs] | System size |
|---|---|---|---|---|---|---|
| 1 | Arginine repressor | O31408 | 149 | 1 - 149 | 2 | 87,444 |
| 2 | WW domain-binding protein 4 | O75554 | 75 | 122 - 196 | 2 | 84,175 |
| 3 | Nuclear factor of activated T-cells, cytoplasmic 2 | Q13469 | 280 | 399 - 678 | 2 | 207,648 |
| 4 | Killer cell immunoglobulin-like receptor 2DL1 | P43626 | 200 | 22 - 221 | 2 | 96,392 |
| 5 | Transcription inhibitor protein Gfh1 | Q8VQD7 | 157 | 1 - 157 | 2 | 110,503 |
| 6 | Protein windbeutel | O44342 | 223 | 21 - 257 | 2 | 131,963 |
| 7 | Sorcin | P30626 | 198 | 1 - 198 | 2 | 112,655 |
| 8 | GTPase Era | P06616 | 301 | 1 - 301 | 2 | 96,207 |
| 9 | Elongation factor Ts | P43895 | 196 | 1 - 196 | 2 | 83,255 |
| 10 | 14-3-3 protein beta/alpha | P31946 | 246 | 1 - 246 | 2 | 162,596 |
| 11 | Chloride intracellular channel protein 1 | O00299 | 241 | 1 - 241 | 2 | 95,819 |
| 12 | Gamma-crystallin B | P02526 | 175 | 1 - 175 | 2 | 61,795 |
| 13 | Transposon Tn7 transposition protein TnsA | P13988 | 273 | 1 - 273 | 2 | 100,299 |
| 14 | NAD(P)H-flavin reductase | P0AEN1 | 233 | 1 - 233 | 2 | 64,350 |
| 15 | Cyclin-dependent kinase 2 | P24941 | 298 | 1 - 298 | 2 | 84,322 |
| 16 | 30S ribosomal protein S7 | P17291 | 156 | 1 - 156 | 2 | 68,090 |
| 17 | Cell division protein ZapA | Q9HTW3 | 104 | 1 - 104 | 2 | 161,499 |
| 18 | Dephospho-CoA kinase | P0A6I9 | 206 | 1 - 206 | 2 | 70,273 |
| 19 | Glutathione S-transferase P 1 | P19157 | 210 | 1 - 210 | 2 | 63,788 |
| 20 | Riboflavin synthase | P0AFU8 | 213 | 1 - 213 | 2 | 119,495 |
| 21 | Calmodulin | O16305 | 149 | 1 - 149 | 2 | 91,499 |
| 22 | Peptidyl-prolyl cis-trans isomerase FKBP42 | Q9LDC0 | 276 | 35 - 310 | 2 | 96,889 |
| 23 | DNA polymerase beta | P06766 | 335 | 1 - 335 | 2 | 98,518 |
| 24 | Ubiquitin | P0CG48 | 76 | 1 - 76 | 1 | 38,843 |
| 25 | Lysozyme C | P00698 | 128 | 1 - 128 | 1 | 49,023 |
| 26 | Green fluorescent protein | P42212 | 238 | 1 - 238 | 1 | 68,624 |
| 27 | Linear diubiquitin | P0CG48 | 152 | 1 - 152 | 2 | 92,946 |
| 28 | ATP-dependent molecular chaperon Hsp82 NTD-MD construct | P02829 | 524 | 1 - 524 | 10.5 | 542,474 |

**Table S2**: Comparison of pLDDT scores across all investigated systems.

| protein | *R* | Intercept | Slope | $pLDDT_{min}$ | $pLDDT_{max}$ | $pLDDT_{median}$ |
|---|---|---|---|---|---|---|
| 1 | 0.78 | 101.40 | -15.71 | 50.44 | 98.7 | 96.28 |
| 2 | 0.94 | 96.68 | -16.51 | 41.88 | 93.4 | 86.92 |
| 3 | 0.82 | 101.83 | -17.68 | 52.21 | 98.31 | 95.23 |
| 4 | 0.44 | 98.29 | -7.17 | 50.92 | 98.53 | 95.76 |
| 5 | 0.88 | 101.77 | -19.13 | 49.45 | 98.58 | 93.40 |
| 6 | 0.90 | 99.58 | -8.43 | 40.73 | 98.63 | 95.55 |
| 7 | 0.89 | 102.11 | -16.7 | 32.44 | 96.50 | 92.64 |
| 8 | 0.76 | 100.11 | -20.14 | 44.39 | 98.37 | 92.10 |
| 9 | 0.28 | 97.67 | -1.29 | 79.10 | 98.87 | 97.63 |
| 10 | 0.96 | 102.41 | -13.37 | 32.90 | 98.88 | 97.87 |
| 11 | 0.79 | 103.07 | -10.78 | 41.48 | 98.77 | 97.19 |
| 12 | 0.76 | 102.42 | -13.35 | 58.02 | 98.79 | 97.79 |
| 13 | 0.70 | 105.04 | -21.67 | 25.47 | 98.84 | 95.52 |
| 14 | 0.65 | 100.58 | -7.29 | 75.14 | 98.85 | 97.35 |
| 15 | 0.45 | 94.53 | -9.49 | 27.73 | 98.81 | 93.88 |
| 16 | 0.89 | 98.11 | -3.60 | 53.49 | 98.06 | 95.81 |
| 17 | 0.85 | 106.31 | -9.72 | 43.54 | 98.29 | 95.94 |
| 18 | 0.78 | 101.74 | -9.87 | 49.82 | 98.72 | 95.72 |
| 19 | 0.24 | 99.13 | -2.02 | 47.46 | 98.94 | 98.64 |
| 20 | 0.77 | 100.94 | -7.43 | 38.72 | 98.77 | 97.41 |
| 21 | 0.61 | 102.00 | -13.26 | 41.16 | 96.28 | 89.62 |
| 22 | 0.51 | 99.19 | -8.62 | 72.32 | 98.74 | 96.74 |
| 23 | 0.77 | 104.01 | -14.21 | 25.10 | 98.93 | 98.05 |
| 24 | 0.99 | 104.21 | -18.68 | 50.34 | 98.63 | 97.97 |
| 25 | 0.85 | 100.37 | -5.43 | 92.48 | 98.91 | 98.68 |
| 26 | 0.85 | 101.16 | -7.38 | 63.53 | 98.88 | 98.46 |
| 27 | 0.94 | 103.50 | -19.57 | 40.70 | 98.43 | 96.80 |
| 28 | 0.59 | 94.07 | -5.78 | 31.83 | 98.35 | 92.47 |

**Table S3**: Comparison of PAE scores across all investigated systems.

| protein | ***R*** | **Intercept** | **Slope** | **PAE$_{median}$** |
|---|---|---|---|---|
| 1 | 0.92 | 2.16 | 2.47 | 10.5 |
| 2 | 0.89 | 4.41 | 1.94 | 14.3 |
| 3 | 0.85 | 3.18 | 2.21 | 8.8 |
| 4 | 0.59 | 1.64 | 3.73 | 4.2 |
| 5 | 0.55 | 4.36 | 0.81 | 6.5 |
| 6 | 0.83 | 2.07 | 2.29 | 5.8 |
| 7 | 0.83 | 4.66 | 4.05 | 10.0 |
| 8 | 0.41 | 6.27 | 2.42 | 9.5 |
| 9 | 0.65 | 2.25 | 0.38 | 3.3 |
| 10 | 0.84 | 1.51 | 2.49 | 3.2 |
| 11 | 0.56 | 0.54 | 2.21 | 2.6 |
| 12 | 0.46 | 1.02 | 2.77 | 2.3 |
| 13 | 0.58 | 0.71 | 4.22 | 3.7 |
| 14 | 0.35 | 2.16 | 0.72 | 2.6 |
| 15 | 0.51 | 4.25 | 1.61 | 4.2 |
| 16 | 0.65 | 2.45 | 0.94 | 3.2 |
| 17 | 0.76 | 1.49 | 1.15 | 5.1 |
| 18 | 0.45 | 2.4 | 0.94 | 3.2 |
| 19 | 0.25 | 1.68 | 0.68 | 1.9 |
| 20 | 0.78 | 1.78 | 1.31 | 2.7 |
| 21 | 0.88 | 1.57 | 2.65 | 13.2 |
| 22 | 0.66 | 2.59 | 4.83 | 7.6 |
| 23 | 0.46 | 2.51 | 0.43 | 3.1 |
| 24 | 0.79 | 0.1 | 4.83 | 1.9 |
| 25 | 0.58 | 0.93 | 2.49 | 1.9 |
| 26 | 0.72 | 1.02 | 2.03 | 2.0 |
| 27 | 0.81 | 2.22 | 1.7 | 7.2 |
| 28 | 0.56 | 5.34 | 0.58 | 9.2 |

**Table S4**: Two-dimensional Kullback-Leibler divergence values of the first two principal component distributions of aMD against cgMD simulations with different ENMs. The cgMD trajectories are projected on the aMD-derived PCs. A lower value corresponds to a higher similarity between both distributions.

| protein | Residues | ENM | AF-ENM | AF-ENM x 2 * | AF-ENM +6% P-W interaction |
|---|---|---|---|---|---|
| 24 | 1 - 70 | 0.8 | 1.1 | 1.2 | |
| 24 | 1 - 76 (all) | 1.1 | 1.8 | 0.7 | |
| 25 | 1 - 128 (all) | 4.5 | 1.4 | 1.3 | |
| 26 | 5 - 228 | 12.1 | 11.4 | 12.0 | |
| 26 | 1 - 238 (all) | 11.0 | 4.8 | 5.2 | |
| 27 | 1 - 152 (all) | 9.4 | 5.9 | 6.1 | |
| 28 | 1 - 210 (NTD) | 11.9 | 1.2 | | 1.2 |
| 28 | 275-524 (MD) | 9.0 | 4.8 | | 4.0 |
| 28 | 1 - 524 (all) | 10.5 | 4.4 | | 4.6 |

* AF-ENM with twice as strong force constants
